# Detecting genotype-population interaction effects by ancestry principal components

**DOI:** 10.1101/719948

**Authors:** Chenglong Yu, Guiyan Ni, Julius van der Werf, S. Hong Lee

## Abstract

Heterogeneity in the phenotypic mean and variance across populations is often observed for complex traits. One way to understand heterogeneous phenotypes lies in uncovering heterogeneity in genetic effects. Previous studies on genetic heterogeneity across populations were typically based on discrete groups of population stratified by different countries or cohorts, which ignored the difference of population characteristics for the individuals within each group and resulted in loss of information. Here we introduce a novel concept of genotype-by-population (G×P) interaction where population is defined by the first and second ancestry principal components (PCs), which are less likely to be confounded with country/cohort-specific factors. We applied a reaction norm model fitting each of 70 complex traits with significant SNP-heritability and the PCs as covariates to examine G×P interactions across diverse populations including white British and other white Europeans from the UK Biobank (*N* = 22,229). Our results demonstrated a significant population genetic heterogeneity for behavioural traits such as age first had sexual intercourse and qualifications. Our approach may shed light on the latent genetic architecture of complex traits that underlies the modulation of genetic effects across different populations.

## Introduction

Most human traits are polygenic and their phenotypes are typically influenced by numerous genes and environmental factors, and possibly by their interactions, e.g. genotype-environment (G×E) interaction ^1-4^. These traits have been termed as “complex traits”, which are distinguished from Mendelian traits that are shaped by a single or few major genes ^5^. Genome-wide association studies (GWAS) have successfully discovered thousands of associations between single-nucleotide polymorphisms (SNPs) and complex traits, which have revolutionized our understanding of the polygenic architecture of complex traits ^6-8^. Subsequently, in order to increase the power and precision to identify more causal variants, there have been numerous follow-up studies using meta-analyses of GWAS summary statistics or mega-analyses of multiple GWAS by combining diverse data sources that usually span across different nations or populations ^9, 10^. However, many human complex traits (e.g., height and body mass index (BMI)) are substantially differentiated among diverse populations ^11^. For instance, the mean height across European nations generally increases with latitude ^12^. Although across-population differences in the mean values are often observed for the phenotypes of complex traits, the underlying genetic and environmental bases remain largely unknown ^12^.

One way to understand such phenotypic heterogeneity lies in uncovering genetic differentiation for the traits captured by common variants across populations. Some studies ^12-15^ have focused on examining population genetic differentiation for several anthropometric, behavioural and psychiatric phenotypes, using whole-genome statistical methods such as applying bivariate genomic restricted maximum likelihood (GREML) to estimate genetic correlation between samples from the USA and Europe for height and BMI ^14^ or determining interaction of genotype by seven sampling populations for behavioural traits by a G-C interaction (GCI)-GREML approach ^15^. They reported significant evidence for G×E interaction in behavioural phenotypes (education and human reproductive behaviour) and BMI ^15^. The analytical method and designs used in their studies were based on discrete groups, which ignored the difference of population characteristics for the individuals within each group. Furthermore, the population groups used in their studies were classified according to their country origin, thus the results were likely to reflect heterogeneity across countries due to country-specific factors (e.g., trait definition and measurement ^16-18^, cultural and societal difference and socio-economic status). In addition, genetic measurement errors (e.g., due to the genotyping platform or imputation quality) across different cohorts may further cause confounding with genuine genetic heterogeneity across populations ^15^.

Principal component (PC) analysis provides a powerful tool to characterize populations and the first few PCs are typically used to control population stratifications in large-scale GWAS ^19^. PCs allow us to cluster individuals that are genetically similar to each other. Unlike discrete variables such as cohort and country, PCs are continuous variables that can differentiate individuals even within a cohort or a country according to their underlying genetic characteristics. Here, we introduce a novel concept of genotype-by-population (G×P) interaction where population is defined by the first and second PCs. It is of interest to test if different genotypes respond differently to the gradient of the first or second PC for complex traits using a whole-genome reaction norm model (RNM) ^20^, which has been recently introduced and allows fitting continuous environmental covariates, i.e. PCs in this study. RNM has been well established to estimate G×E interaction in agriculture ^21, 22^ and ecology ^23^. Furthermore, in this study we used the data source of UK BioBank (UKBB), which is a prospective cohort study with deep genetic and phenotypic data collected on approximately 500,000 individuals across the United Kingdom, aged between 40 and 69 at recruitment ^24, 25^.

Therefore, in our G×P interaction model applied to UKBB, the population characteristics for individuals are fully utilised and the findings are less likely to be confounded with country-specific factors or genetic measurement errors as mentioned above.

The aim of the study is to explore if there exists significant G×P interaction, which is also referred to genetic heterogeneity (heterogeneous genetic effects) across populations, for a wide range of complex traits. To do so, we applied the whole-genome RNM with PCs as continuous covariates to investigate G×P interactions for more than one hundred phenotypes using the UKBB data. The significant G×P interaction detected in this study may shed light on the latent genetic architecture of complex traits that underlies the modulation of genetic effects across different population backgrounds.

## Subjects and Methods

### Data and quality control (QC)

Our study was based on the UKBB data which contains approximately 500,000 individuals sampled across the United Kingdom ^25^. According to the ethnic background (data field 21000), there are currently 472,242 individuals with the white British ancestry and 17,038 individuals with any other white ethnic background (not with British or Irish ethnicity) in the UKBB participants. In order to match the sample size between the white British and the other white ethnic individuals, we randomly selected 17,000 individuals from the white British group, totalling 34,038 admixed European populations considered in this study. Using ancestry PCs provided by the UKBB, we examined a two-dimensional scatter plot of the first and second PC of the 17,000 white British and the 17,038 other white ethnic subjects (Figure 1A). It is shown that the white British group is situated within the group of the other white Europeans and we named the white British group as POP1 (*N*=17,000). As shown in Figures 1B and 1C, we used a geometric method by which we constructed a rectangle with maximums and minimums of PC1 and PC2 of the white British group as four sides and then group the individuals of the other white Europeans inside this rectangle, named as POP3 (*N*=9,809). The rest of the other white Europeans except POP3 were named as POP2 (*N*=7,229).

**Figure 1.**
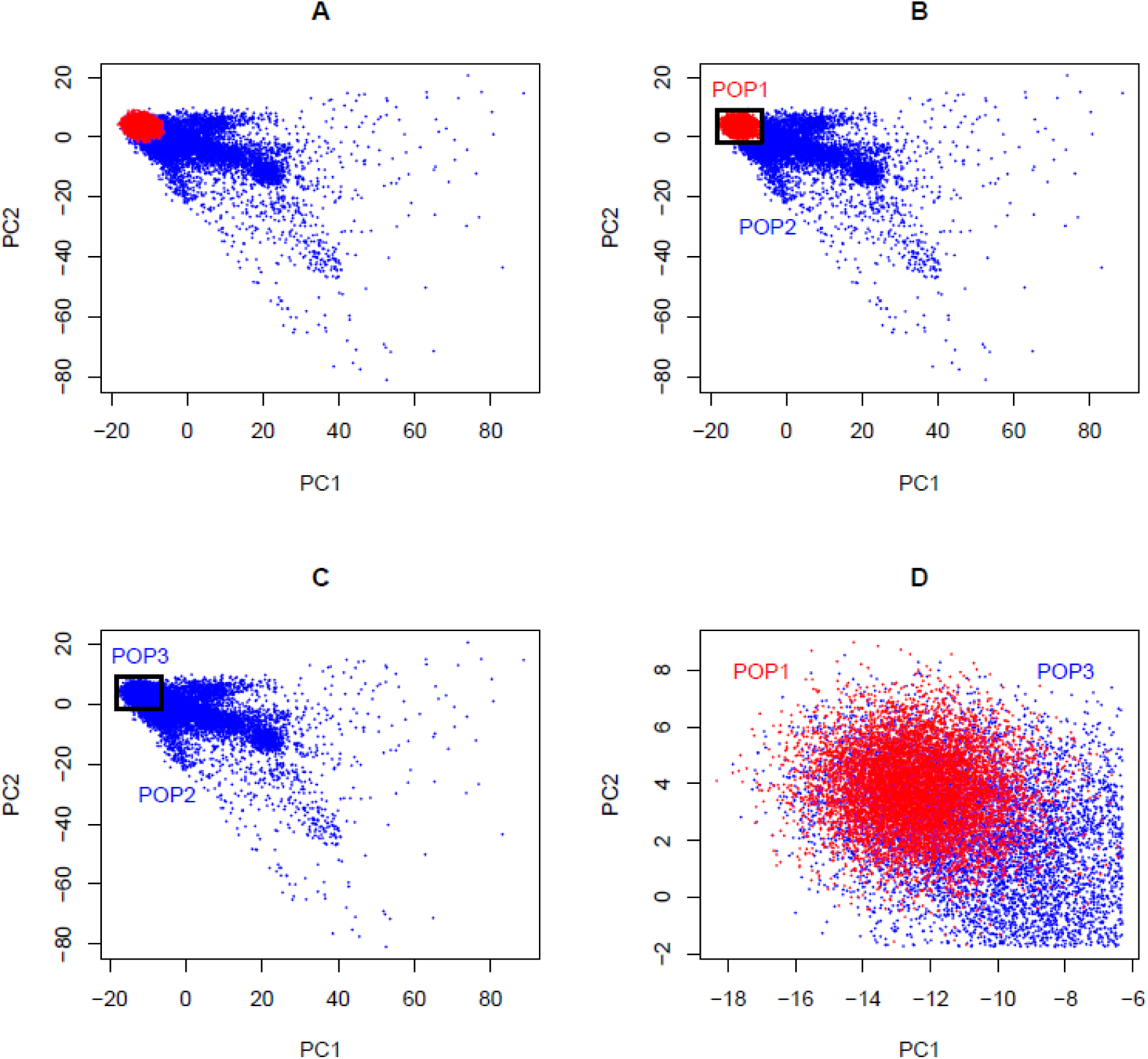
Two-dimensional scatter plots of PC1 and PC2 with red points representing white British individuals and blue points representing other white ethnic individuals from the UKBB. The white British group named as POP1 is situated within the group of the other white Europeans. As shown in (B) and (C), we used a geometric method by which we constructed a rectangle with maximums and minimums of PC1 and PC2 of POP1 as four sides and then group the individuals of the other white Europeans inside this rectangle, named as POP3. The rest of the other white Europeans except POP3 were named as POP2.

Our primary interest was to investigate G×P interaction where population was classified by ancestry PCs. For this purpose, we used three designs of combinations of the three groups, i.e. POP1+POP2 (Figure 1B), POP2+POP3 (Figure 1C) and POP1+POP3 (Figure 1D). It was noted that POP1+POP3 was a negative control as there was little population difference among them. To make sample size consistent across POP1 and POP2 in the design of POP1+POP2, we randomly selected 7,500 individuals from the 17,000 white British individuals and these were used as POP1 in the downstream analyses.

We extracted genetic data including around 92 million SNPs from the UKBB for all the individuals of POP1, POP2 and POP3. Stringent QC was applied to the combined data across POP1, POP2 and POP3. The QC criteria were to exclude 1) all duplicated and non-autosomal SNPs, 2) SNPs with INFO score < 0.6, 3) SNPs with call rate < 0.95; (4) individuals with missing rate > 0.05, 5) SNPs with Hardy-Weinberg equilibrium *p*-value < 0.0001, 6) SNPs with minor allele frequency < 0.01, and 7) SNPs with A/T alleles or G/C alleles. We also retained HapMap3 SNPs only as they are reliable and robust to bias in estimating SNP-heritability and genetic correlation ^15, 26, 27^. Hereafter, 1,133,957 common SNPs were remained for the G×P analyses. Moreover, we excluded one individual randomly selected from any pair with a genetic relationship > 0.05 (see Statistical models) to avoid bias due to confounding by shared environment among close relatives. After the QC, the sample sizes of POP1, POP2 and POP3 were reduced to 7487, 6913 and 7829.

### UKBB phenotypes

For current UKBB resource, we have access to 496 variables whose data types are categorical (multiple), categorical (single), continuous, integer, date, text and time. Here we focused on the variables of categorical (multiple), categorical (single), continuous and integer types, and categorized each variable as one of four value types: continuous, binary, ordered categorical and unordered categorical ^28^ (Table S1). Where a data field is measured at several time points we use the first occurrence only. It was noted that qualifications (data field 6138), a categorical (multiple) trait, was reorganised according to the underlying system ^29^. Briefly, the original and unordered seven categories were reclassified and ordered as 1) none, 2) O-levels or CSEs, 3) A-levels, NVQ, HND, HNC or other professional qualification, and 4) college or university degree. Then the continuous, binary and ordered categorical variables were selected and used as the main phenotypes in G×P interaction analyses.

Since there exist numerous “Not Available” (NA) records for individuals in UKBB, the limited sample sizes of some variables may lead to insufficient statistical power to perform our study. Hence, the variables with limited sample size should be excluded. As POP2 and POP3 have the same ethnic background, we only examined sample sizes of POP1 and POP2, and used the following thresholds to exclude the variable with: non-NA number in POP1 < 2,500 and non-NA number in POP2 < 2,500, and then we remain 199 variables whose sample size in POP1+POP2 > 5,000 as shown in Table S1. Note that some ambiguous values in variables such as “Do not know” or “Prefer not to answer” were treated as NA.

Among the 199 variables, we selected 128 variables as the main phenotypes (Table S2) in our proposed model to estimate G×P interactions where population difference was inferred from the first and second PCs. The other variables were used to control confounding effects owing to sex, age, year of birth, genotype batch and assessment centre (basic confounders adjusted for all the main phenotypes; the first 20 PCs were also used as basic confounders to account for population stratification) and Townsend deprivation index, smoking status, alcohol consumptions and many other variables (additional cofounders adjusted for some relevant phenotypes) or excluded if they were not likely to affect any of the main phenotypes (see the note of Table S2). The 128 main phenotypes could be classified into a number of criteria, 1) lifestyle and environment (alcohol, diet, electronic device use, sexual factors, sleep, smoking and sun exposure), 2) physical measures (anthropometry, blood pressure and bone-densitometry of heel), 3) early life factors, sociodemographics (education, employment and household), 4) health and medical history (eyesight, hearing, medical conditions and medication), 5) psychosocial factors (mental health), 6) female-specific factors, male-specific factors, 7) verbal interview (medical conditions) and 8) cognitive function (reaction time) (Table S2). Note that some phenotypes such as from sociodemographics (e.g., qualifications) can also be used as additional confounders for other phenotypes.

### Statistical models

#### *A linear mixed model without considering* G×P *interaction (baseline model)*

A standard linear mixed model assuming no G×P interaction can be written as

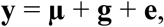

where **y** is an *n*×1 vector of phenotypes with *n* being the sample size, **µ** is an *n*×1 vector for fixed effects, **g** is an *n*×1 vector of total genetic effects of the individuals with 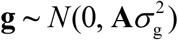 and **e** is an *n*×1 vector of residual effects with 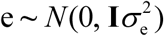, where 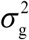 is the variance explained by all common SNPs and 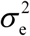 is the residual variance. In the GREML context ^30, 31^, **A** is a genomic relationship matrix (GRM) and **I** is an identity matrix. GRM can be estimated based on common SNPs across the genome and the elements of GRM can be defined as ^30, 32, 33^:

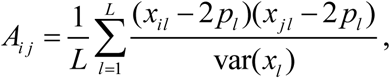

where *L* is the number of all common SNPs (*L* = 1,133,957 in this study), *x*_*il*_ denotes the number of copies of the reference allele for the *l*th SNP of the *i*th individual, *x*_*l*_ denotes all the numbers of copies of the reference allele across all the individuals, and *p*_*l*_ denotes the reference allele frequency of the *l*th SNP.

The variance-covariance matrix of the observed phenotypes (**V)** is

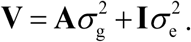

The SNP-based heritability, the proportion of the additive genetic variance explained by the genome-wide SNPs over the total phenotypic variance, is then referred as

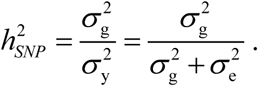

The phenotypes with significant SNP-based heritability from this baseline model will subsequently be investigated for G×P interaction.

#### G×P RNM method

In cases where G×P interaction exists across populations, the baseline model cannot account for heterogeneous genetic effects. We therefore applied RNM methods to detect heterogeneity across populations using the UKBB data. RNM and multivariate RNM (MRNM) have been demonstrated to perform better than the current state-of-the-art methods when detecting genotype-covariate and residual-covariate interactions in terms of simulation studies on type I error rate and power analyses ^20^. Here we focus on G×P interaction by considering PCs as covariates in the RNM:

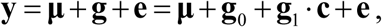

where **y, µ, g** and **e** are the same defined in the baseline model above, **g**_0_ and **g**_1_ are *n*×1 vectors of zero- and first-order random regression coefficients, respectively, **c** is an *n*×1 vector of covariate values of the *n* individuals (for which we used PC1 and PC2 values in this study). In the RNM, the random genetic effects, **g**, are regressed on the covariate gradient (reaction norm), which can be modelled with random regression coefficients, **g**_0_ and **g**_1_. The variance-covariance matrix of the random regression coefficients (**K**) is

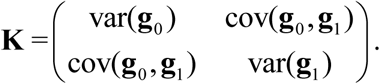

Then the variance-covariance matrix of genetic effects between *n* individuals (who have unique PC values) can be expressed as

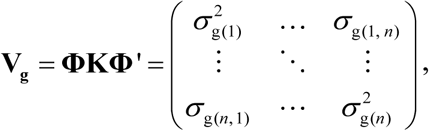

where 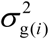 denotes the genetic variance at the *i*th covariate level, 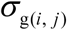 indicates the genetic covariance between the *i*th and *j*th covariate levels (*i* = 1, …, *n*, and *j* = 1, …, *n*), and 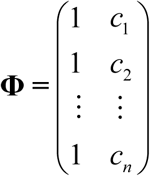 denotes the covariate matrix. This G×P RNM accounts for phenotypic plasticity and norms of reaction in response to different populations (represented by PC values) among samples.

The mathematical properties of **K** allow us to verify whether estimates of the parameters are reasonable or not. Specifically, estimated values in the matrix **K** should be within a valid parameter space:

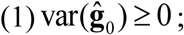

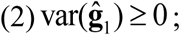

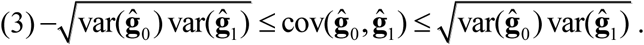

The estimates which violated one of above criteria were excluded for follow-up analyses. We obtained a p-value to detect G×P interaction using a likelihood ratio test (LRT) that compared the goodness of fitness of two models (GREML and G×P RNM), penalising the difference in the number of parameters between them.

We further tested if the significant G×P interactions were orthogonal (independent without confounding) to residual-population (R×P) interactions, i.e. residual heterogeneity across populations ^20^. Similarly, the R×P interaction can be detected by an R×P RNM:

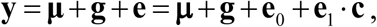

where **e**_0_ and **e**_1_ are vectors of zero- and first-order random regression coefficients when residual effects, **e**, are regressed on the covariate, **c**, i.e. an n vector of PC1 or PC2.

Furthermore, a full RNM model with both G×P and R×P interactions can be expressed as

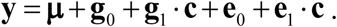

Since the G×P and R×P models are nested within the full model, LRT comparing the full and R×P or G×P model with an appropriate degree of freedom can determine the significance of orthogonal G×P or R×P interaction ^20^.

For the analyses showing a significant G×P interaction, we used rank-based INT phenotypes to check explicitly if the significance was due to phenotypic heteroscedasticity or normality assumption violation ^34^. The bias of RNM/MRNM estimates due to non-normality of phenotypic values can also be remedied by applying the rank-based INT ^20^. All models described above (i.e., GREML, bivariate GREML, RNM, MRNM) can be fitted using software MTG2 ^33^.

#### Spurious signals due to selection or collider bias

We used the UKBB data that have only a 5.5% response rate, i.e. selection. Consequently, the resulting sample may not be representative of the UK population as a whole and the selection may be associated with some of the phenotypes in the UKBB, causing selection or collider bias ^35, 36^. To test whether the G×P interaction effects detected by our method was genuine or spurious due to selection or collider bias, we conducted a series of simulation studies with phenotypes differentially selected for POP1 (white British) and POP2 (other white Europeans). A selection model using a logistic regression with a trait Y can be written as

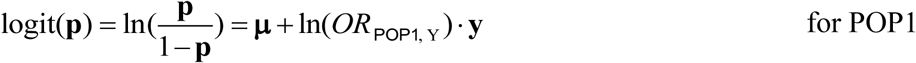

and

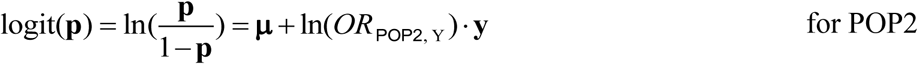

where **p** is a vector of participation probabilities in a study (e.g., UKBB questionnaire survey) for all individuals, **µ** is an overall mean vector which regulates the response rate, **y** is a vector of phenotypic values of the trait Y, *OR*_POP1,Y_ and *OR*_POP2,Y_ are selection odds ratios for POP1 and POP2, respectively. Then the sample selection bias can be simulated with varying selection odds ratios.

One hundred replicates of phenotypic values of the trait Y on POP1+POP2 (14,400 individuals) were simulated under a baseline model (GREML) that assumes no G×P interaction: **y** = **g** + **e**, where the variance-covariance structure between **g** and **e** was 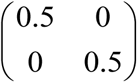. For each replicate, to avoid insufficient statistical power, we set **µ** as a vector of zeros which simulates a response rate of 50%. Letting us divide **y** and **p** into subsets according to specific populations (i.e. **y**_1_ and **p**_1_ for POP1 and **y**_2_ and **p**_2_ for POP2), we obtain the participation probability for each individual as

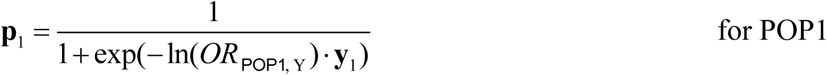

and

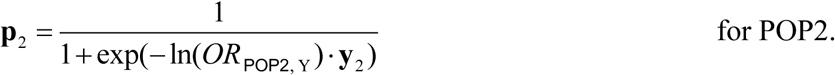

Then, individuals in each population are selected based on the participation probability. Specifically, we generate a uniform distribution vector **u**_1_ on (0, 1) with sample size of POP1, and compare the values of corresponding components in **p**_1_ and **u**_1_. The individuals having larger values in **p**_1_ than in **u**_1_ are assumed to participate in this study. Similarly, we can select individuals in POP2 by comparing **p**_2_ with a random number drawn from a uniform distribution (0, 1). Different combinations of selection odds ratios for POP1 and POP2 (e.g., *OR*_POP1,Y_ = 1 and *OR*_POP2,Y_ = 2) will generate selection bias associated with phenotypic values in the POP1+POP2 groups.

Since the phenotypic data was simulated under the null model, a significant G×P interaction detected from LRT comparing G×P RNM versus GREML was a type I error (false positive). This allowed us to investigate the type I error rate of G×P interaction due to selection bias attributed to various selection pressures (odds ratios) on POP1 and POP2. Using the same simulated data, we also applied a bivariate GREML ^37^ to test if estimated genetic correlation between POP1 and POP2 was significantly different from 1 (i.e. evidence of G×P interaction across POP1 and POP2) ^38^. This allowed us to assess the type I error rate of G×P interaction when using the bivariate GREML.

If two or more phenotypic variables simultaneously influence the probability of participation of individuals in a study, then investigating associations between those variables in the selected sample may induce collider bias ^36^. Therefore, we further considered the same selection model but including two traits to evaluate collider bias effects on the detection of G×P interaction across POP1 and POP2. The selection model with two traits Y and Z can be written as

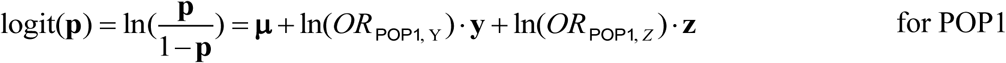

and

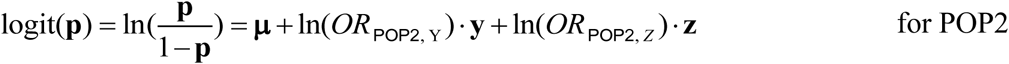

where **z** is a vector of phenotypic values of the trait Z, *OR*_POP1, *Z*_ and *OR*_POP2, *Z*_ are selection odds ratios with the trait Z for POP1 and POP2. The magnitude of collider bias depends on the levels of selection odds ratios for the two phenotypes.

We simulated 100 replicates of phenotypic values of the trait Z on POP1+POP2 under the null model of no G×P interaction: **z** = **α** + **β**, where the variance-covariance structure between **α** and **β** is 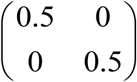. Since genetic components **g** and **α** are uncorrelated and residual components **e** and **β** are uncorrelated, the phenotypic variable **z** and previous simulated **y** = **g** + **e** are totally independent, but after selection we expect that the two variables will be associated because of a collider. Letting us divide **z** into subsets according to specific populations (i.e. **z**_1_ for POP1 and **z**_2_for POP2), the individuals can be selected based on

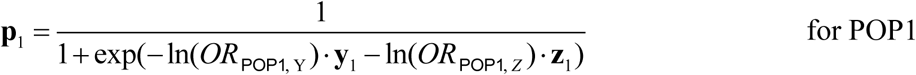

and

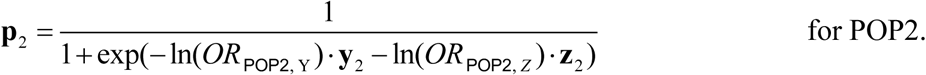

Similarly, we can select individuals in POP1 or POP2 by comparing **p**_1_ or **p**_2_ with a random number drawn from a uniform distribution (0, 1). Therefore, in terms of collider bias, different combinations of selection odds ratios for different traits and populations (e.g., *OR*_POP1,Y_ = 2, *OR*_POP2,Y_ = 3, *OR*_POP1, *Z*_ = 2 and *OR*_POP2, *Z*_ = 2) will generate collider bias in the POP1+POP2 groups. Similarly, we can examine G×P interaction by type I error rate analysis using G×P RNM and bivariate GREML methods, and assess collider bias effects for the two methods.

## Results

### Estimating SNP-based heritability for 128 phenotypes

We first applied the standard GREML model to estimate 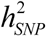 for the 128 phenotypes across POP1+POP2, POP2+POP3 and POP1+POP3, respectively. The phenotypes with significant 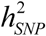 (Tables S3-5) were further investigated for G×P interaction effects using our G×P RNM approach.

### Genetic and residual correlations between phenotypes and PCs

The main response (**y**) and environmental covariates (**c**) are not always uncorrelated, for which multivariate RNM accounting for (genetic and residual) correlations between **y** and **c** should be used ^20^. We examined if there were non-negligible genetic and residual covariances between the main phenotypes and covariate (PC1 or PC2) for the complex traits with significant heritabilities (Tables S6-8). All genetic and residual covariances estimated by bivariate GREML were not significantly different from zero, and thus we used univariate RNM to detect the G×P interaction effects with covariate PC1/PC2 for those phenotypes.

### G×P interaction

For POP1+POP2, we fit the data of the 70 phenotypes with significant 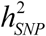 by modelling the G×P RNM with covariates PC1 and PC2, respectively (Tables S9 and S10). We excluded those estimates, which were not within the valid parameter space (see Statistical models), from the follow-up statistical test analyses, resulting in 29 and 32 traits remaining for PC1 (Table S9) and PC2 analyses (Table S10). We examined if there was significant G×P interaction and obtained p-values based on LRT comparing the fit to the data of the G×P RNM and null model. Significance level was determined by Bonferroni multiple testing correction: 0.05/140 = 3.57E-4 for the 70 phenotypes with covariates PC1 and PC2. Figures S1A and S1B show that significant G×P interactions were found for ten complex traits which are related to blood pressure (pulse rate, automated reading), bone-densitometry of heel (heel BMD T-score, automated; heel broadband ultrasound attenuation, direct entry; heel QUI, direct entry; heel BMD), diet (lamb/mutton intake), sexual factor (age first had sexual intercourse), sleep (sleep duration), smoking (ever smoked) and education (qualifications). For each of the ten traits, we further considered a multiple covariate model that fit PC1 and PC2 jointly (Table S11). However, G×P interactions were less significant than those obtained using the single covariate model fitting PC1 or PC2 separately (Figure S2), otherwise, the estimates were out of the valid parameter space. This was probably due the fact that there was collinearity between G×P interactions from PC1 and PC2.

In addition to the basic confounders for which the main phenotypes were initially adjusted (see Subjects and Methods), we further considered additional trait-specific confounders that might be relevant to some of traits (Table S2), e.g. Townsend deprivation index, smoking status, alcohol drinker status, etc. After controlling for additional trait-specific confounders, the G×P interactions in POP1+POP2 were still significant for bone-densitometry of heel (heel BMD T-score, automated; heel broadband ultrasound attenuation, direct entry; heel QUI, direct entry; heel BMD), age first had sexual intercourse and qualifications, whereas the signals disappeared for the other traits (Table S12).

We examined the distribution of phenotypic values after controlling additional confounders of the six traits with significant G×P interactions (Figure S3) and could not rule out the possibility that the interaction signals were due to non-normality (e.g. residual heteroscedasticity). We conducted the same analyses for the six traits using rank-based INT phenotypes (Table 1), which could control type I error rate due to a skewed and non-normal distribution of residual values ^20^. Indeed, phenotypic heteroscedasticity was remedied when using rank-based INT for the phenotypes of six traits as shown in Figures S4-9. We found that the interaction signals of age first had sexual intercourse and qualifications were remained significant even after applying rank-based INT phenotypes, however, the other traits were not significant anymore (Table 1).

**Table 1.** Genetic variance, interaction variance and their covariance component estimates for six phenotypes across POP1+POP2 with the covariates PC1 and PC2. The phenotypes were adjusted by basic plus additional confounders of fixed effects and transformed by rank-based INT. The estimates which were not within the valid parameter space are marked as “Excluded”. SE denotes standard error. DF denotes degree of freedom.

| UKBB data field | Phenotype | Covariate | $\text{var}(\mathbf{g}_0)$<br>(SE) | $\text{var}(\mathbf{g}_1)$<br>(SE) | $\text{cov}(\mathbf{g}_0, \mathbf{g}_1)$<br>(SE) | $\text{var}(\mathbf{e}_0)$<br>(SE) | *P*-value by LRT comparing with baseline model (DF = 2) |
| --- | --- | --- | --- | --- | --- | --- | --- |
| 78 | Heel bone mineral density (BMD) T-score, automated | PC1 | 0.3151(0.0459) | 0.0124(0.0110) | 0.0013(0.0120) | 0.6739(0.0456) | 0.1586 |
|  |  | PC2 | 0.3187(0.0460) | -0.0008(0.0047) | -0.0037(0.0110) | 0.6838(0.0450) | Excluded |
| 3144 | Heel Broadband ultrasound attenuation, direct entry | PC1 | 0.2754(0.0454) | 0.0094(0.0110) | 0.0087(0.0120) | 0.7161(0.0454) | 0.0789 |
|  |  | PC2 | 0.2774(0.0454) | 0.0006(0.0048) | -0.0024(0.0111) | 0.7232(0.0450) | 0.8987 |
| 3147 | Heel quantitative ultrasound index (QUI), direct entry | PC1 | 0.3151(0.0459) | 0.0124(0.0110) | 0.0013(0.0120) | 0.6739(0.0456) | 0.1597 |
|  |  | PC2 | 0.3187(0.0460) | -0.0009(0.0047) | -0.0037(0.0110) | 0.6839(0.0450) | Excluded |
| 3148 | Heel bone mineral density (BMD) | PC1 | 0.3070(0.0458) | 0.0107(0.0109) | 0.0046(0.0120) | 0.6836(0.0455) | 0.1315 |
|  |  | PC2 | 0.3106(0.0459) | -0.0016(0.0046) | -0.0069(0.0110) | 0.6926(0.0450) | Excluded |
| 2139 | Age first had sexual intercourse | PC1 | 0.1006(0.0266) | 0.0080(0.0078) | 0.0203(0.0087) | 0.8909(0.0290) | 5.16E-05 |
|  |  | PC2 | 0.1012(0.0266) | 0.0110(0.0057) | -0.0015(0.0087) | 0.8880(0.0286) | 0.0097 |
| 6138 | Qualifications | PC1 | 0.1194(0.0235) | 0.0706(0.0103) | -0.0791(0.0090) | 0.8124(0.0261) | 9.21E-18 |
|  |  | PC2 | 0.1778(0.0214) | 0.0360(0.0059) | 0.0833(0.0081) | 0.7885(0.0233) | 2.22E-24 |

For age first had sexual intercourse and qualifications that were shown to have significant G×P interactions, we further tested if the G×P interactions were orthogonal to R×P interactions, i.e. residual heterogeneity (see Subjects and Methods). Using the rank-based INT phenotypes adjusted for basic and additional confounders, we carried out an R×P model and a full model in which both G×P and R×P were fitted jointly. Subsequently, we conducted LRT to obtain p-values, comparing the full and nested models. A significant p-value from LRT between the full and R×P model indicates that G×P interaction is orthogonal to R×P interaction (see Subjects and Methods and Tables S13). For age first had sexual intercourse, although G×P or R×P interaction was significantly detected from the G×P or R×P model, it was shown that G×P interaction was not orthogonal to R×P (p-value = 0.88 for PC1 and 0.92 for PC2 in Table S13). For qualifications, on the other hand, it was shown that the G×P and R×P interactions were statistically independent (p-value = 4.15E-05 for PC1 and 0.003 for PC2 in Table S13).

For POP2+POP3, we conducted analyses using the same procedure as in the analyses of POP1+POP2. The POP3 individuals are very close to those in POP1 in terms of ancestry PC, but their ethnicities are not white British as in POP1 (see Subjects and Methods and Figure 1). Thirteen phenotypes demonstrated a significant genetic heterogeneity for covariate PC1 or PC2 as shown in Tables S14 and S15. After controlling for additional trait-specific confounders and transforming by rank-based INT (Table S16), the results for behavioural phenotypes age first had sexual intercourse (p-value = 7.86E-05 for PC1) and qualifications (p-value = 1.06E-15 for PC1) have demonstrated strong genetic heterogeneity signals, which are consistent with our findings for POP1+POP2. For qualifications, G×P interactions were significantly orthogonal to R×P interactions (p-value = 0.003 for PC1 in Table S17). We also found significant results across POP2+POP3 for anthropometric traits (waist circumference and weight) and diabetes diagnosed by doctor. However, these phenotypes were not discovered across POP1+POP2 with significant G×P interaction signals. We presented genetic variance, interaction variance and their covariance component estimates for these significant traits across POP2+POP3 in Table 2.

**Table 2.**
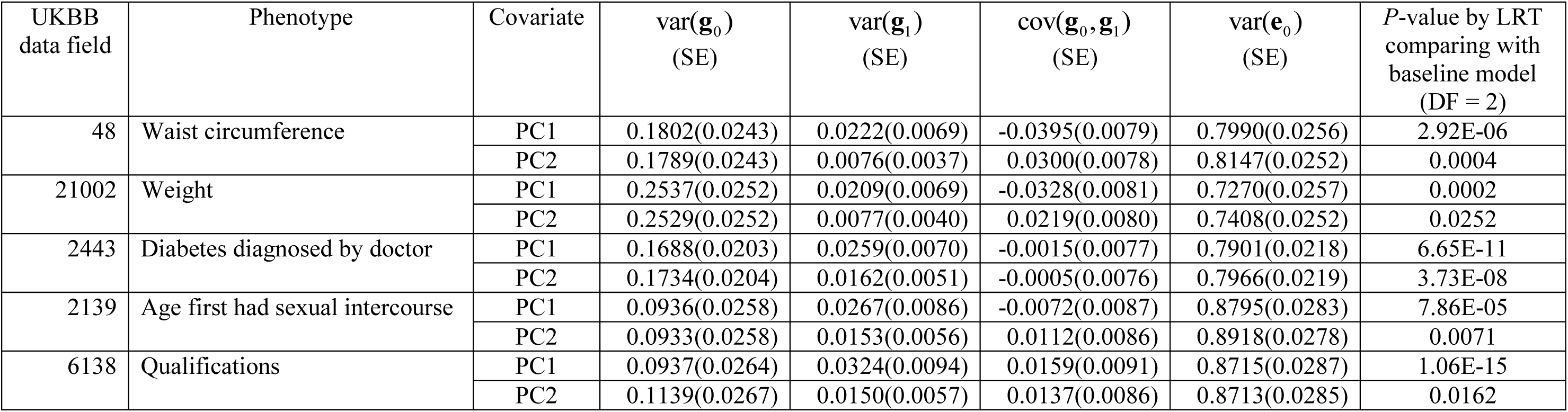
Genetic variance, interaction variance and their covariance component estimates for six phenotypes across POP2+POP3 with the covariates PC1 and PC2. The phenotypes were adjusted by basic plus additional confounders of fixed effects and transformed by rank-based INT. SE denotes standard error. DF denotes degree of freedom.

We also performed the same analyses on POP1+POP3, which is not a diverse population group as POP1+POP2 or POP2+POP3, and thus was used as a negative control group (see Methods). For several traits showing significant heterogeneous signals with covariate PC1 or PC2 after Bonferroni correction (see Tables S18 and S19), we further examined them by adding stringent confounders to correct for fixed effects and applying rank-based INT. The final results included no significant G×P interaction across POP1+POP3 (see Tables S20 and S21).

For the categorical phenotype qualifications, there were various ways to convert the seven UKBB categories into a continuous or a binary measure ^39, 40^. Following a previous study ^39^, we transformed the multiple categories (data fields: 6138.0.0 to 6138.0.5) into an educational year measure (Table S22). Based on this continuous phenotypic measure, we found significant genetic heterogeneity across POP1+POP2 and POP2+POP3 but no signal across POP1+POP3 (Table S23), which was consistent with our results obtained using four-level categories. We also examined G×P interactions for qualifications based on two types of binary measures (highest educational attainment versus other levels, and lowest educational attainment versus other levels)^40^. The results were consistent with those obtained using four-level qualifications, except that an unexpected significant signal across POP1+POP3 for covariate PC1 was detected based on the binary measure of “college or university degree” versus other six categories (Table S24).

### Testing effects of selection or collider bias

We examined the distribution of phenotypic values for age first had sexual intercourse and qualifications in which G×P interactions were consistently detected from both POP1+POP2 and POP2+POP3 (Tables S25 and S26). The distribution of age first had sexual intercourse is similar across POP1, POP2 and POP3. However, for qualifications, it is apparently shown that the subjects in POP2 and POP3 (other white Europeans) have higher qualification levels than those in POP1 (white British). Moreover, it is likely that the individuals in POP1 have higher educational levels than the general population of UK because individuals with higher educational levels are more likely to response to surveys from UKBB ^36^.

Our simulation studies testing for detecting spurious heterogeneity across POP1 and POP2 with multiple scenarios varying the level of selection odds ratios (see Supplementary Notes for details) have verified that (1) both G×P RNM and bivariate GREML are robust to the selection bias when using the same selection odds ratio across populations (Table 3); (2) only bivariate GREML is robust against the selection bias when using different selection odds ratios across populations (Table 3); (3) bivariate GREML is robust against the collider bias when estimating genetic correlation between POP1 and POP2, however it generates biased estimation of genetic correlation between the two traits (Table 4). It is noted that the level of selection odds ratios used in simulations is likely to reflect the real situation of qualifications, i.e. different selection pressure between POP1 and POP2 in UKBB (see Supplementary Notes and Tables S27).

**Table 3.** Simulation study results for selection bias on the phenotype Y across POP1+POP2. Different odds ratio combinations (*OR*_POP1,Y_ and *OR*_POP2,Y_) generated phenotypic values in POP1+POP2 with different selection bias levels. Type I error rates based on 100 simulation replicates were examined by G×P RNM and bivariate GREML respectively. The genetic correlations of the phenotype between POP1 and POP2 were estimated by bivariate GREML. SE denotes standard error.

| Selection scenarios in<br>POP1+POP2 | Type I error rate<br>by G×P RNM with PC1 | Type I error rate<br>by bivariate GREML | 100 estimated genetic correlations |  |
| --- | --- | --- | --- | --- |
|  |  |  | Mean | SE |
| $OR_{POP1, Y} = 1, OR_{POP2, Y} = 1$ | 5% | 0% | 0.9722 | 0.0145 |
| $OR_{POP1, Y} = 1, OR_{POP2, Y} = 2$ | 55% | 2% | 0.9876 | 0.0166 |
| $OR_{POP1, Y} = 2, OR_{POP2, Y} = 2$ | 1% | 0% | 1.0245 | 0.0160 |
| $OR_{POP1, Y} = 2, OR_{POP2, Y} = 3$ | 64% | 6% | 0.9882 | 0.0202 |

**Table 4.** Simulation study results for collider bias on two phenotypes Y and Z across POP1+POP2. Different odds ratio combinations and *OR*_POP1, Y_, *OR*_POP2, Y_, *OR*_POP1, *Z*_ and *OR*_POP2, *Z*_) generated phenotypes in POP1+POP2 with different selection bias levels. Type I error rates based on 100 simulation replicates were examined through estimated genetic correlations of the phenotype Y between POP1 and POP2 by bivariate GREML. SE denotes standard error.

| Selection scenarios with collider bias in POP1+POP2 | Type I error rate | Estimated genetic correlations of the phenotype Y between POP1 and POP2 |  | Estimated genetic correlations between Y and Z on selected POP1+POP2 |  |
| --- | --- | --- | --- | --- | --- |
|  |  | Mean | SE | Mean | SE |
| $OR_{POP1, Y} = 2$, $OR_{POP1, Z} = 2$,<br>$OR_{POP2, Y} = 3$, $OR_{POP2, Z} = 2$ | 1% | 1.0141 | 0.0189 | -0.2516 | 0.0032 |
| $OR_{POP1, Y} = 2$, $OR_{POP1, Z} = 2$,<br>$OR_{POP2, Y} = 3$, $OR_{POP2, Z} = 3$ | 2% | 1.0220 | 0.0165 | -0.2942 | 0.0031 |
| $OR_{POP1, Y} = 2$, $OR_{POP1, Z} = 3$,<br>$OR_{POP2, Y} = 3$, $OR_{POP2, Z} = 3$ | 2% | 1.0091 | 0.0187 | -0.3415 | 0.0036 |

For age first had sexual intercourse and qualifications, we confirmed our findings using bivariate GREML, a robust approach against selection bias (Table 5). The bivariate GREML results for qualifications indicated a significant genetic heterogeneity between POP1 and POP2 (p-value = 8.09E-04), and between POP2 and POP3 (p-value = 7.85E-04), but showed no genetic heterogeneity between POP1 and POP3. These results were consistent with our findings from the G×P RNM. For age first had sexual intercourse, the bivariate GREML detected a significant heterogeneity between POP2 and POP3 (p-value = 3.14E-05), however, there was no interaction signal between POP1 and POP3 (as expected). Unexpectedly, the bivariate GREML failed to find genetic heterogeneity across POP1+POP2 (Table 5) although RNM provided a significant signal.

**Table 5.** Genetic correlation estimates between population groups (POP1, POP2 and POP3) by bivariate GREML for two phenotypes. Here the phenotypes were adjusted by basic plus additional confounders of fixed effects and transformed by rank-based INT. SE denotes standard error. P-value was obtained through a Wald test under a null hypothesis that genetic correlation equals to 1.

| Phenotype | Genetic correlation<br>between POP1 and POP2 |  |  | Genetic correlation<br>between POP2 and POP3 |  |  | Genetic correlation<br>between POP1 and POP3 |  |  |
| --- | --- | --- | --- | --- | --- | --- | --- | --- | --- |
|  | Estimate | SE | P value | Estimate | SE | P value | Estimate | SE | P value |
| Qualifications | 0.2554 | 0.2223 | 8.09E-04 | 0.4795 | 0.1550 | 7.85E-04 | 0.5676 | 0.2743 | 0.1149 |
| Age first had sexual intercourse | 0.7418 | 0.3984 | 0.5169 | 0.0491 | 0.2284 | 3.14E-05 | 1.2176 | 0.3629 | 0.5488 |

As confirmed by the bivariate GREML, it was not likely that the findings for qualifications were spurious because of selection and collider bias. This was also evidenced by the fact that G×P RNM detected a significant interaction signal from POP2+POP3, noting that POP2 and POP3 were similarly distributed for qualifications (see Table S26). Similarly, the findings for age first had sexual intercourse were mostly robust whether using RNM or bivariate GREML except that there was no signal for POP1+POP2 when using the bivariate GREML, probably due to the lack of power. It was noted that the phenotypic distributions of age first had sexual intercourse were very similar across POP1, POP2 and POP3 (Table S25).

### Hidden heritability

For the significant traits qualifications and age first had sexual intercourse, we examined SNP-based heritabilities estimated by GREML and G×P RNM (see Table S28). The phenotypic values were adjusted for basic and additional confounders of fixed effects and transformed using rank-based INT. For POP1+POP2, the SNP-based heritability for qualifications estimated by G×P RNM increased by 28% (from 0.0998 to 0.1281) and 84% (from 0.0998 to 0.1840) with covariate PC1 and PC2, compared to those estimated by GREML. But there was no such apparent increase of estimated SNP-based heritability for POP2+POP3 and POP1+POP3 when comparing GREML and G×P RNM.

## Discussion

Previous results ^12-15^ were more likely to reflect heterogeneous genetic effects across nations or cohorts rather than populations as those designs were evidently confounded with country-specific factors (e.g., trait definition and measurement, cultural and societal difference). In this study, we focused on populations and proposed the new concept “genotype-population interaction” in which population is defined by the first and second ancestry PCs (each individual has a unique PC value). Using the RNM with whole-genome data from the UKBB, we have demonstrated significant G×P interaction effects for qualifications and age first had sexual intercourse across populations. Our findings corroborate the results in Tropf et al. ^15^ who reported that behavioural phenotypes (education and human reproductive behaviour) have significant G×E interactions across populations. For anthropometric phenotypes, height and BMI, our G×P RNM model did not detect any significant interaction signals^14^. However, the analyses of another two anthropometric traits (waist circumference and weight) have demonstrated significant genetic heterogeneity across the POP2+POP3 group (other white Europeans). Actually, the results by Tropf et al. ^15^ across seven populations have also revealed significant G×E interaction for BMI although the heterogeneity is not strong as for education and reproductive behaviours. Robinson et al. ^12^ also reported that, for BMI, environmental differences across Europe masked genetic differentiation. Thus, these findings may be consistent for some anthropometric phenotypes when using diverse European ancestry populations. From the POP2+POP3 analyses, we also found a significant G×P interaction for diagnosis of diabetes that is a binary response variable.

As the RNM has not been explicitly verified for binary traits, we also used bivariate GREML to estimate the genetic correlation between POP2 and POP3 for this disease trait and found no significant signal for genetic heterogeneity (estimate is 0.7988, SE = 0.2044, p-value = 0.3249). This might be due to the fact that there was no genuine interaction effects or that the bivariate GREML was simply underpowered. For the two binary measuring ways of qualifications (lowest educational attainment versus other levels, and highest educational attainment versus other levels), we also used bivariate GREML to examine genetic correlations between POP1, POP2 and POP3 (Table S29). The results for the binary phenotype of “none of the above” versus other six educational categories demonstrated significant genetic heterogeneity between POP1 and POP2 (p-value = 5.58E-05) and between POP2 and POP3 (p-value = 7.59E-05) but no significant signal between POP1 and POP3 (p-value = 0.0619), which were consistent with those obtained from the main analyses. For the binary data of “college or university degree” versus other six categories, the bivariate GREML indicated a marginally significant heterogeneity between POP1 and POP2 (p-value = 0.035) and no significant signal between POP2 and POP3 (p-value = 0.494), and POP1 and POP3 (p-value = 0.94). The reason that the genetic heterogeneity became weaker or disappeared is probably due to the fact that the bivariate GREML has less power compared to the RNM approach, and the phenotype categories reduced from four to two levels.

Our results imply that causal variants at multiple loci may not be universal but rather specific to populations for some complex traits. The results on qualifications across POP1+POP2 suggested that G×P interaction might be a reason for attenuation of SNP-based heritability when using data from different populations, for which we hold the same view as by Tropf et al. ^15^. This missing or hidden heritability issue ^41^ can produce lower predictive power of polygenic risk scores from large GWAS (usually generated from meta-analyses of different populations) compared with single homogenous population since the reference heritability obtained from the meta-analyses among several populations is smaller than that obtained from single homogenous population ^42^. Therefore, our findings suggest that large homogeneous population data sources (e.g., around 400,000 white British individuals in the UKBB) should be used to conduct genetic risk prediction for some specific traits such as human behaviors.

The current methods used for estimating G×E (or G×P) interactions, e.g. random regression (RR)-GREML ^22^ and GCI-GREML ^12, 15^, require that the main response should be stratified into multiple discrete groups according to covariate levels even for a continuous covariate ^13^. However, the arbitrary grouping ignores the difference of covariate values for the individuals within each group, and results in some loss of information. In contrast, the RNM allows us to fit a continuous covariate representing individuals uniquely (e.g. PC) in the model and produces unbiased estimates ^20^. In our results, bivariate GREML which labels the individuals into two discrete groups (POP1 and POP2) failed to find genetic heterogeneity for age first had sexual intercourse (Table S27), while RNM detected significant G×P interaction across POP1+POP2 (see Table 1). It may imply that G×P RNM is more powerful as it uses individual-level information represented by PC across populations, while bivariate GREML ignores such information within each stratified group. However, on the other hand, RNM may suffer from the selection bias when using different selection odds ratios across populations (Table 3) while bivariate GREML is robust against such selection and collider bias (Tables 3 and 4).

Residual-covariate interaction may result in heterogeneous residual variances across different covariate values, thus it is necessary to examine and distinguish genotype-covariate and residual-covariate interactions ^20^. Our results (Tables S13 and S17) provided cogent evidence of G×P and R×P interaction effects, which are (partially) independent without confounding, across populations for qualifications. However, for age first had sexual intercourse, there was no evidence showing that G×P interaction was orthogonal to R×P interaction from LRT comparing the full and nested models. Therefore, we could not rule out the possibility that the significant signal was mainly because of residual heterogeneity across populations. In order to disentangle G×P interaction from R×P interaction, the magnitude of G×P interaction should be large (e.g. qualifications) or sample size may have to be increased.

The previous results ^12-15^ were based on pooled data across different nations, and thus various trait definitions in phenotypic measure and genetic measurement errors across countries may generate artificial heterogeneity. In our study, however, we used the data resource standardized across one country (the United Kingdom) to rule out those cross-country factors and influences. The phenotypic definitions and measurement of complex traits in this cohort have been standardized nationwide. Moreover, UKBB utilized uniform standards of imputation and quality control for genotype data and provided genotyping batch information for each individual that was used as fixed effect adjusted in our models. Therefore, our results may reflect authentic G×P interaction effects across populations.

There are several limitations in this study. Firstly, we examined G×P interaction across populations using three data designs (POP1+POP2 and POP2+POP3 as primary data, and POP1+POP3 as a negative control), in which population is referred to the first and second ancestry PCs. As POP1 and POP3 are very close in terms of PCs, the individuals in the two primary groups POP1+POP2 and POP2+POP3 have common population structures (Figure 1). But both groups involve in different white ethnic backgrounds, i.e., POP1 may be closer to native British and POP2/POP3 is more likely to be descended from recent immigrants from many other European nations. Therefore, for our data designs, we cannot rule out the possibility that G×P interaction was confounded with immigration-specific factors such as socioeconomic attainment, social relations and cultural beliefs ^43^. We also notice that, in the UKBB data source, there are numerous samples with other ethnicities (e.g., Indian, Caribbean and African), thus future studies using our approach may aim to detect genotype-ethnicity interaction, which may uncover challenges for investigations into the genetic architecture of phenotypes across various ethnicities. Secondly, population defined by PCs in this study or by discrete groups in others ^14, 15^ includes both environmental and genetic information for individuals, thus the G×P interaction may not merely embody G×E interaction but also contains confounded genotype-by-genotype (G×G) interaction across populations. It may become a new challenge in the future to distinguish G×E and G×G in studies of genetic heterogeneity across populations. Thirdly, the sample size for people with other white ethnicity in UKBB (i.e., the sum of POP2 and POP3) is not large, thus the study may lack power for phenotypes with small SNP-based heritability such as behavioural traits. The phenotypes without significant heritability in the current samples were not investigated for G×P interaction, however, if boosting statistical power for those phenotypes, there may be new findings for heterogeneity across populations. Fourthly, the simulations on selection bias have demonstrated that the G×P RNM is not robust for data across populations with different selection odds ratios (see Table 3). Thus our approach is more preferable and restricted to data without selection bias or with the same selection pressure for populations. Finally, for genotypic information used in this study, we only examined common SNPs (minor allele frequency > 0.01). However, a recent study ^44^ reported that the missing heritability for height and BMI may be explained by rare genetic variants accessed from whole-genome sequence data. Therefore, can rare population-specific variants increase our understanding of genetic heterogeneity across populations? Further research is required to answer this question.

In conclusion, our study provided a paradigm shift tool in investigating genetic heterogeneity across populations. The new concept of G×P interaction with the use of ancestry PC is more plausible in explaining the genetic architecture of complex traits across heterogeneous populations. The G×P interaction effects on behavioural phenotypes (qualifications and age first had sexual intercourse) were found by a powerful approach based on technically homogeneous data (free of genetic measurement errors and cohort/country confounding factors), and these findings were validated in both data designs POP1+POP2 and POP2+POP3. The analyses performed in this study can be applied to dissect the genetic architecture of complex traits and diseases across populations, and the results from these analyses will provide important information and suggestion for studies of genomic risk prediction across Europeans.

## Supplemental Data

Supplemental data file includes supplemental notes, 9 figures and 29 tables and can be found with this article online.

## Acknowledgments

This research is supported by the Australian Research Council (DP190100766, FT160100229), and the Australian National Health and Medical Research Council (1087889). This research has been conducted using the UK Biobank Resource. UK Biobank (http://www.ukbiobank.ac.uk) Research Ethics Committee (REC) approval number is 11/NW/0382. Our reference number approved by UK Biobank is 14575.

## Declaration of Interests

The authors declare no conflict of interest.

## Web Resources

UK Biobank, http://www.ukbiobank.ac.uk

PLINK v1.90, https://www.cog-genomics.org/plink2

MTG2, https://sites.google.com/site/honglee0707/mtg2

