## Supplemental Data for "Detecting genotype-population interaction effects by ancestry principal components"

This file includes supplemental notes, 9 figures and 29 tables.

### Supplementary Notes

*Simulation study on selection bias*

We conducted a series of simulation studies by examining different odds ratio combinations for selection in POP1 and POP2 to evaluate influence of detecting G×P interaction due to selection bias (see Subjects and Methods). For the 100 replicates of phenotype simulated under a null model, we adjusted the values by first 20 PCs and ethnicity (POP1 or POP2) as fixed effects and transformed by rank-based INT. We then used G×P RNM with covariate PC1 and baseline GREML models to fit the 100 simulated replicates. Since the data was simulated under a null model, we consider a significant G×P interaction signal detected through LRT of G×P RNM versus GREML as a type I error (false positive). The results as shown in Table 3 indicated high type I error rates (> 50%) for selection scenarios of $OR_{\text{POP1, Y}} = 1$, $OR_{\text{POP2, Y}} = 2$ and $OR_{\text{POP1, Y}} = 2$, $OR_{\text{POP2, Y}} = 3$. But for same selection pressures ($OR_{\text{POP1, Y}} = 1$, $OR_{\text{POP2, Y}} = 1$ and $OR_{\text{POP1, Y}} = 2$, $OR_{\text{POP2, Y}} = 2$), G×P RNM can effectively control false positive rate. Type I error was also evaluated through genetic correlation estimated by bivariate GREML for the phenotype between POP1 and POP2 with the null assumption that genetic correlation = 1 implies no heterogeneity. As shown in Table

3, type I error rates assessed by bivariate GREML were controlled for all scenarios and the estimated genetic correlations were not significantly different from one for all selection scenarios.

To verify whether the selection odds ratios we simulated are comparable to real situation (qualifications), we performed linear regression including 1 (representing POP1) and 2 (representing POP2) as independent variable and educational levels as dependent variable on real data of qualifications across POP1+POP2. Similarly, we also performed 100 linear regressions for the 100 replicates simulated with selection odds ratio combination of $OR_{\text{POP1, Y}} = 2$ and $OR_{\text{POP2, Y}} = 3$. We then examined R-squared and p-values obtained from linear regressions for real data of qualifications and simulation data (Table S27). The regression R-squared values of simulations were similar to those of real data, which indicated that selection bias by $OR_{\text{POP1, Y}} = 2$ and $OR_{\text{POP2, Y}} = 3$ in simulation has been sufficient to reflect realistic situation of educational levels across POP1+POP2 in UKBB. Therefore, our simulation studies have verified that the G×P interaction of qualifications across populations we detected may not be due to selection bias at this scenario.

*Simulation study on collider bias*

We also performed simulation studies to evaluate collider bias effects on G×P interaction detected by bivariate GREML (see Subjects and Methods) as shown in Table 4. Type I error rates under the null hypothesis that genetic correlation of 1 implies no interaction were controlled for several combinations of $OR_{\text{POP1, Y}}$, $OR_{\text{POP2, Y}}$, $OR_{\text{POP1, }Z}$ and $OR_{\text{POP2, }Z}$, and meanwhile, significant negative genetic correlations between the two simulated phenotypes Y and Z demonstrated strong collider bias signal in selected POP1+POP2. Here we assume that the phenotype Z involves a sum of collider bias effects across all other traits on the main response Y. We further examined R-squared and p-values obtained from linear regressions for those simulation data with different collider bias levels (Table S27). The similar R-squared values indicated that our simulated collider bias settings have been sufficient to represent real situation of qualification levels. The results implied that collider bias may not be a cause for significant G×P interaction of qualifications across POP1+POP2.

**Figure S1 (A)**

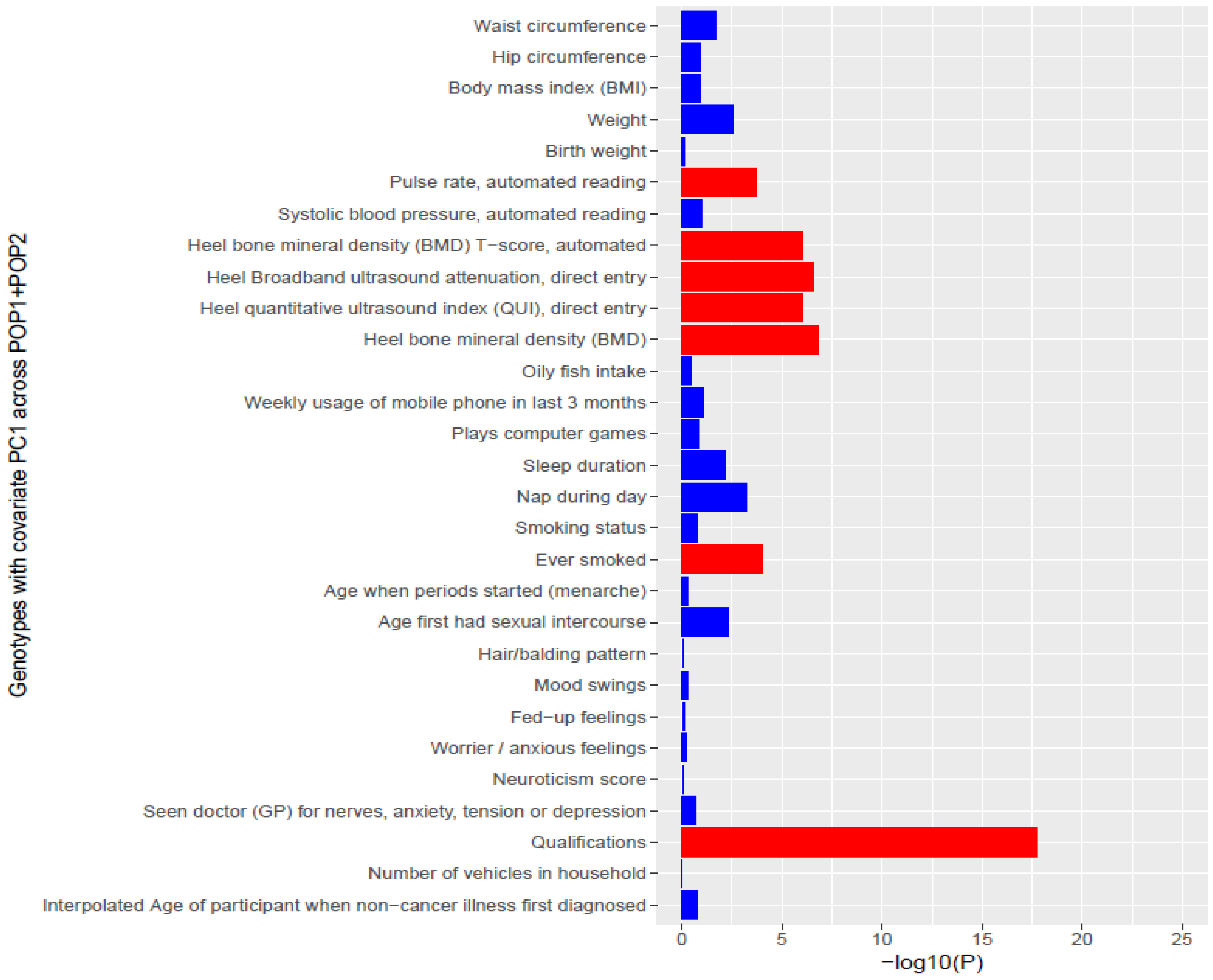
Waist circumference
Hip circumference
Body mass index (BMI)
Weight
Birth weight
Pulse rate, automated reading
Systolic blood pressure, automated reading
Heel bone mineral density (BMD) T−score, automated
Heel Broadband ultrasound attenuation, direct entry
Heel quantitative ultrasound index (QUI), direct entry
Heel bone mineral density (BMD)
Oily fish intake
Weekly usage of mobile phone in last 3 months
Plays computer games
Sleep duration
Nap during day
Smoking status
Ever smoked
Age when periods started (menarche)
Age first had sexual intercourse
Hair/balding pattern
Mood swings
Fed−up feelings
Worrier / anxious feelings
Neuroticism score
Seen doctor (GP) for nerves, anxiety, tension or depression
Qualifications
Number of vehicles in household
Interpolated Age of participant when non−cancer illness first diagnosed
Genotypes with covariate PC1 across POP1+POP2
0
5
10
15
20
25
−log10(P)

**Figure S1 (B)**

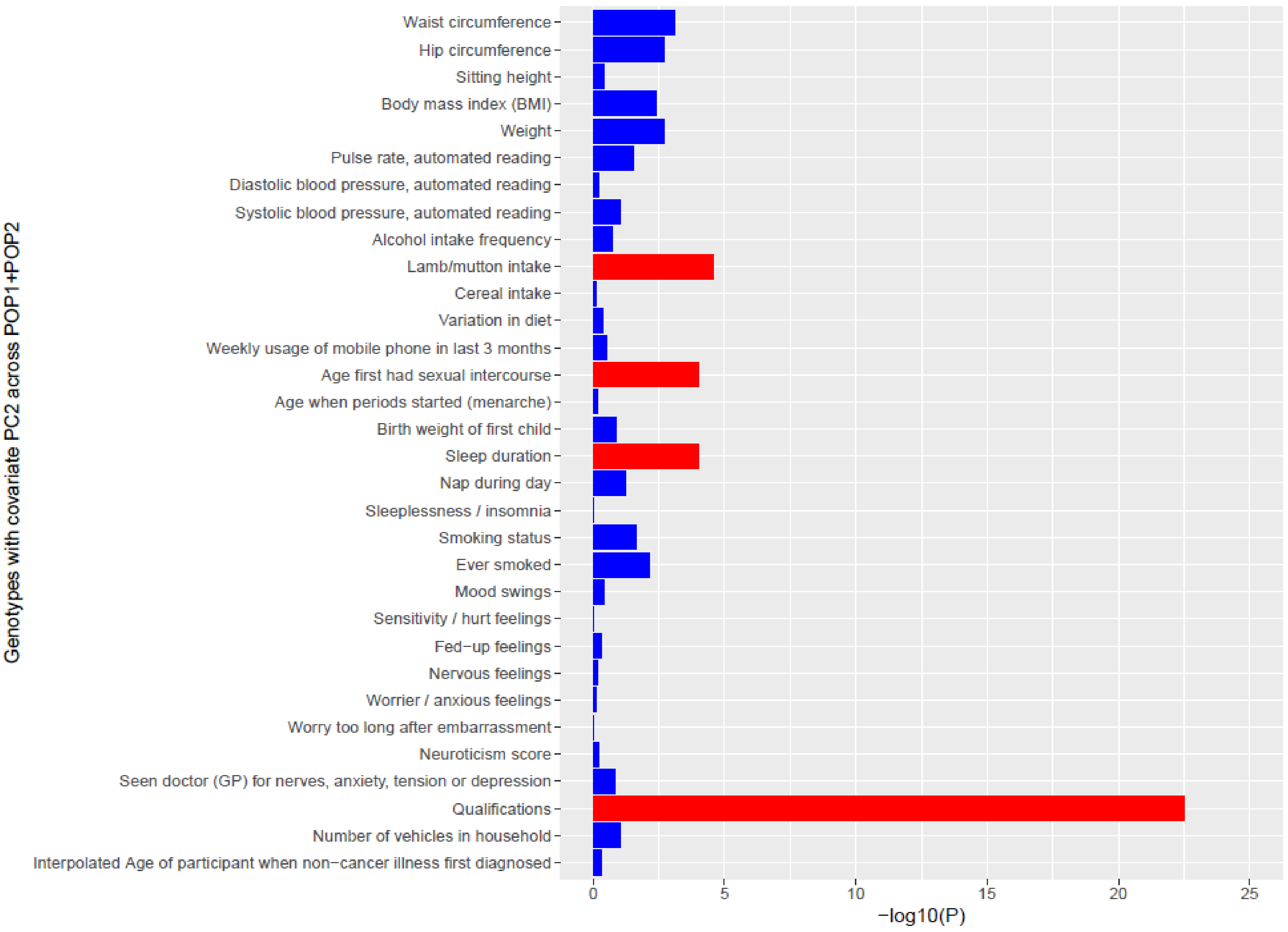
Genotypes with covariate PC2 across POP1+POP2
Waist circumference
Hip circumference
Sitting height
Body mass index (BMI)
Weight
Pulse rate, automated reading
Diastolic blood pressure, automated reading
Systolic blood pressure, automated reading
Alcohol intake frequency
Lamb/mutton intake
Cereal intake
Variation in diet
Weekly usage of mobile phone in last 3 months
Age first had sexual intercourse
Age when periods started (menarche)
Birth weight of first child
Sleep duration
Nap during day
Sleeplessness / insomnia
Smoking status
Ever smoked
Mood swings
Sensitivity / hurt feelings
Fed−up feelings
Nervous feelings
Worrier / anxious feelings
Worry too long after embarrassment
Neuroticism score
Seen doctor (GP) for nerves, anxiety, tension or depression
Qualifications
Number of vehicles in household
Interpolated Age of participant when non−cancer illness first diagnosed
0
5
10
15
20
25
−log10(P)

**Figure S1. Detecting G×P interaction for (A) 29 traits with covariate PC1 and (B) 32 traits with covariate PC2**. The phenotypic values for each trait were adjusted by basic confounders of fixed effects. Red colour indicates significant genetic heterogeneous signals (significance level was determined by Bonferroni multiple testing criterion: 0.05/140 = 3.57E-4).

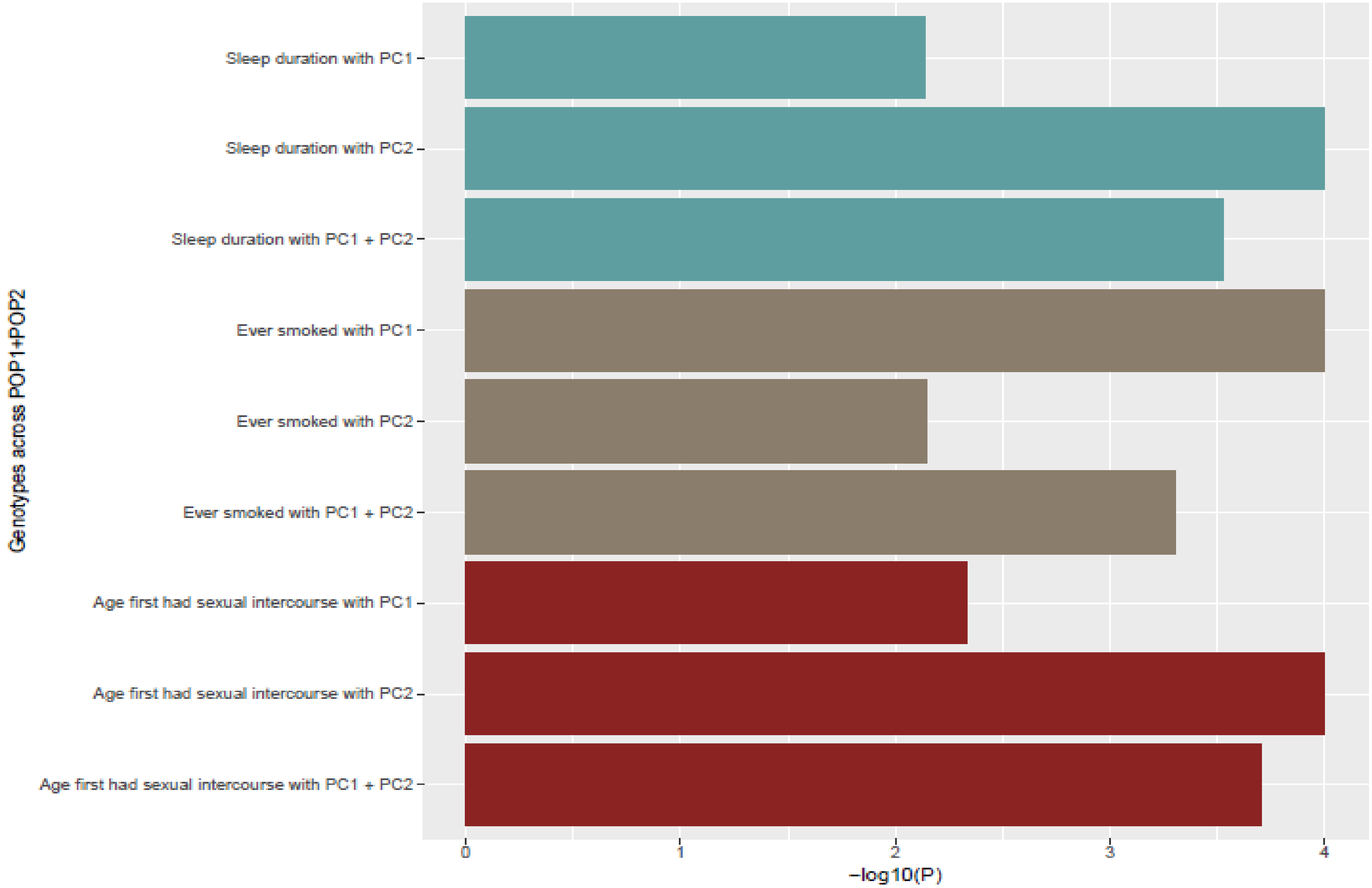


**Figure S2**. **Comparisons of G×P interaction patterns using PC1, PC2 or simultaneous PC1 and PC2 for three valid traits in Table S11**.

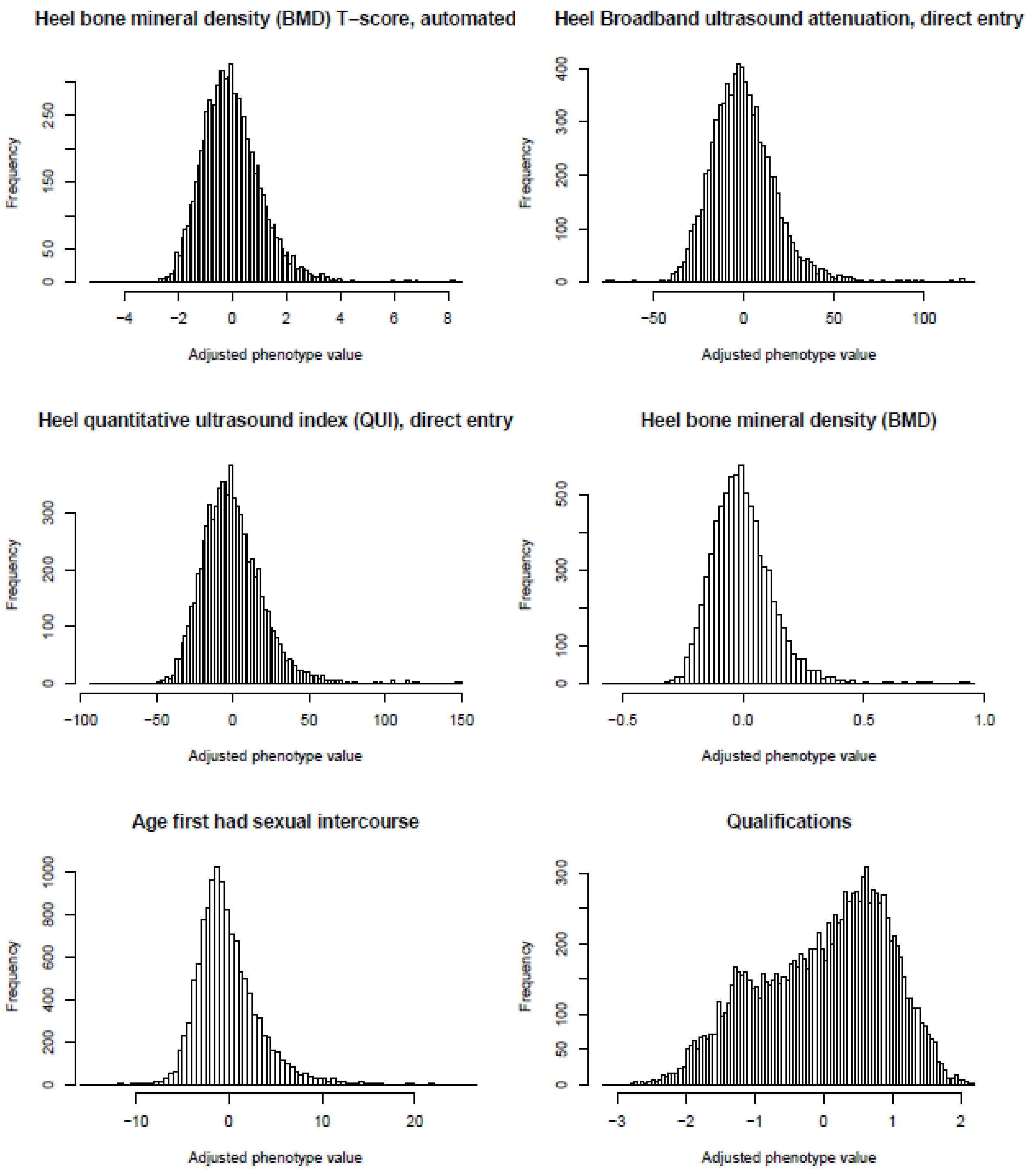


**Figure S3. Distributions of phenotypic values after controlling basic and additional confounders of the six traits with significant G×P interactions.**

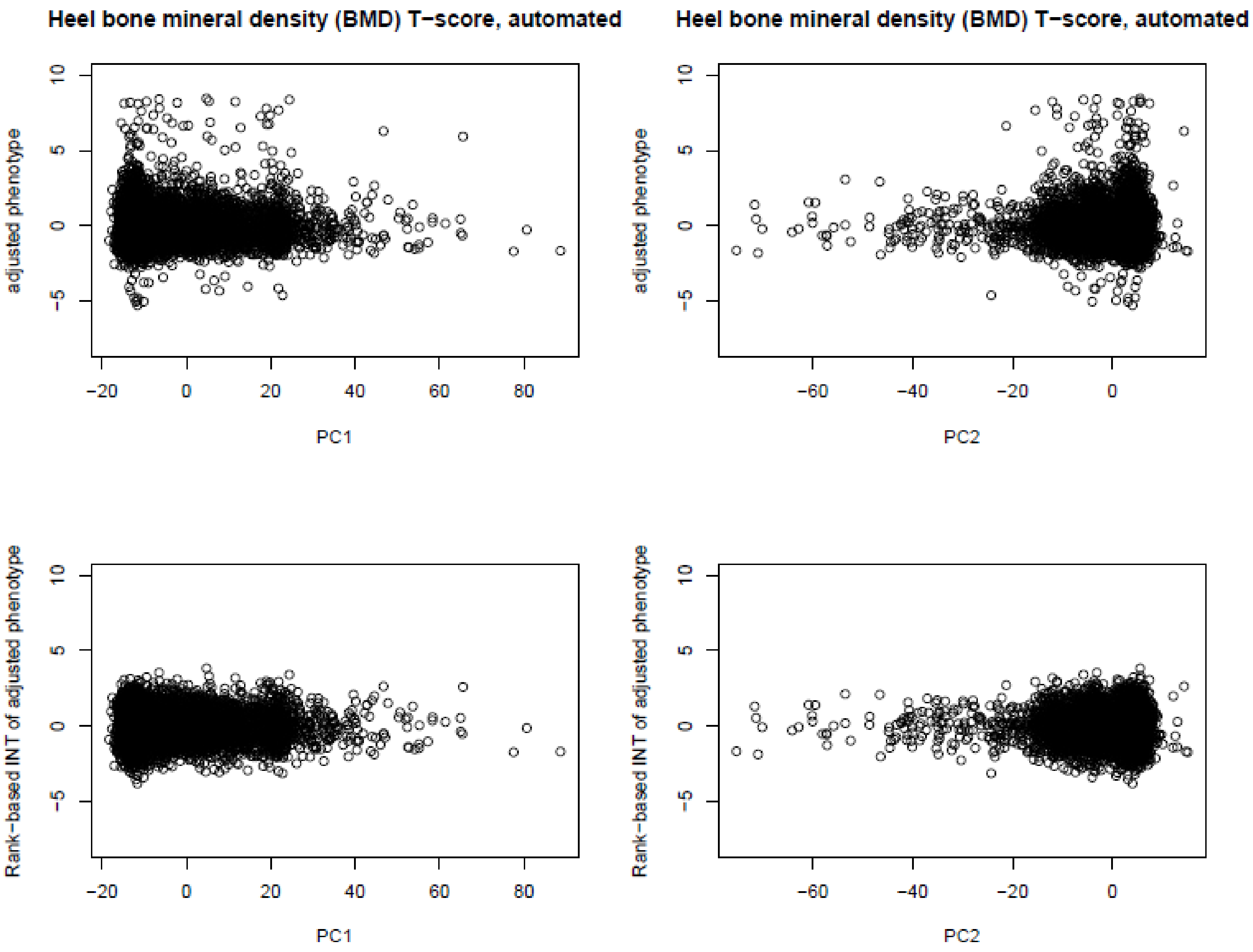


**Figure S4. Phenotypic heteroscedasticity of the trait "heel bone mineral density (BMD) T-score, automated" before and after rank-based INT.** The phenotypes were adjusted by basic and additional confounders of fixed effects.

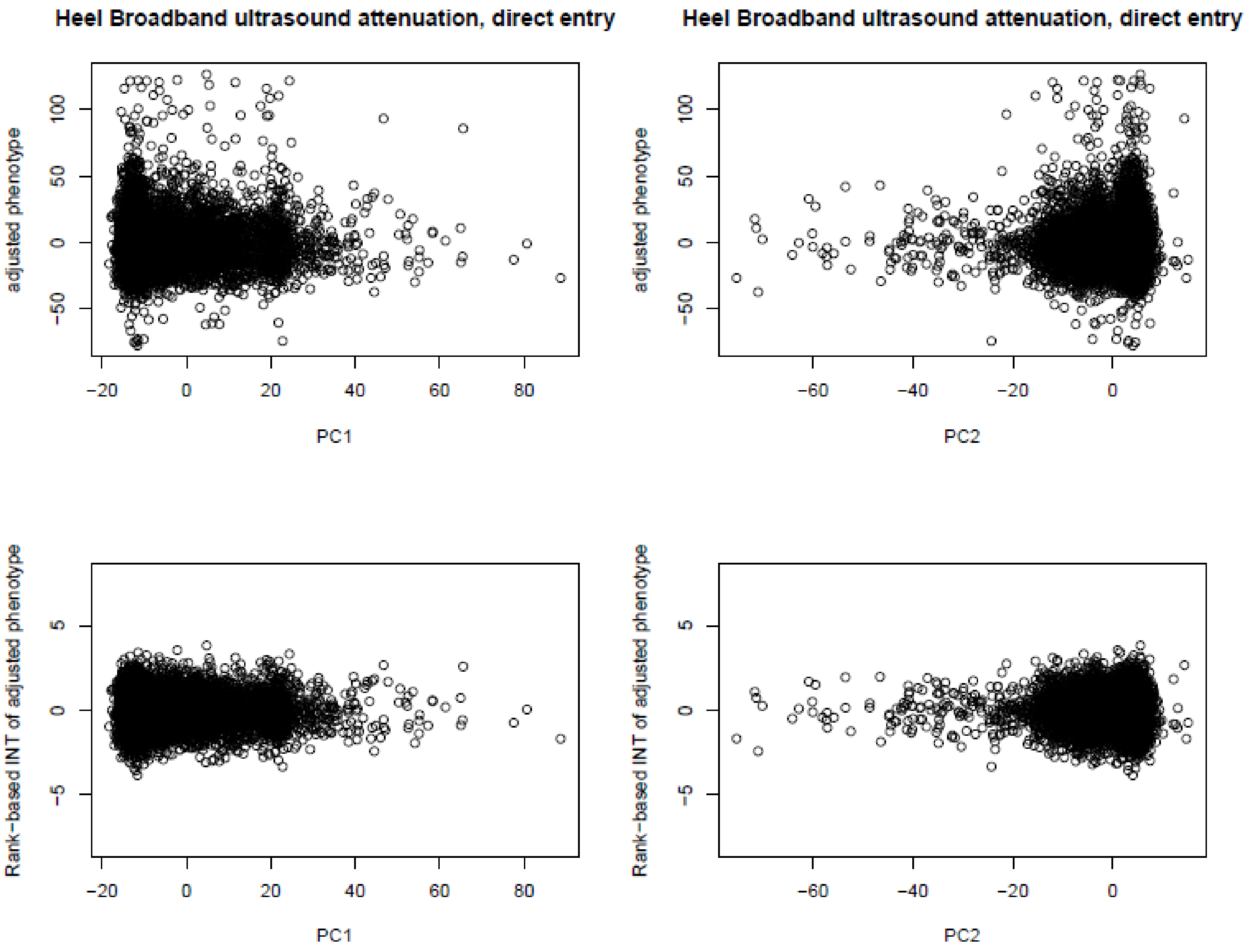


**Figure S5. Phenotypic heteroscedasticity of the trait "heel broadband ultrasound attenuation, direct entry" before and after rank-based INT.** The phenotypes were adjusted by basic and additional confounders of fixed effects.

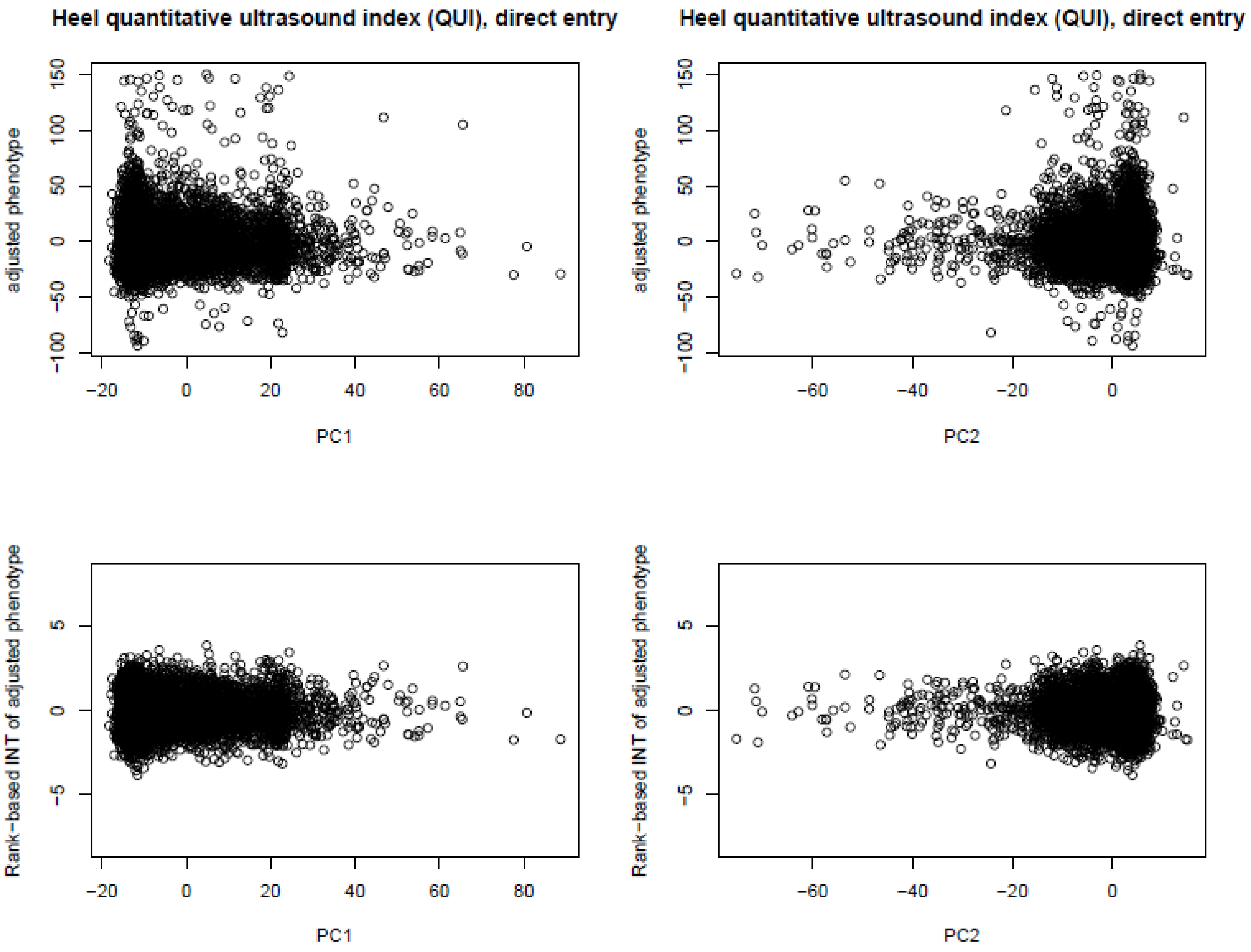


**Figure S6. Phenotypic heteroscedasticity of the trait "heel QUI, direct entry" before and after rank-based INT.** The phenotypes were adjusted by basic and additional confounders of fixed effects.

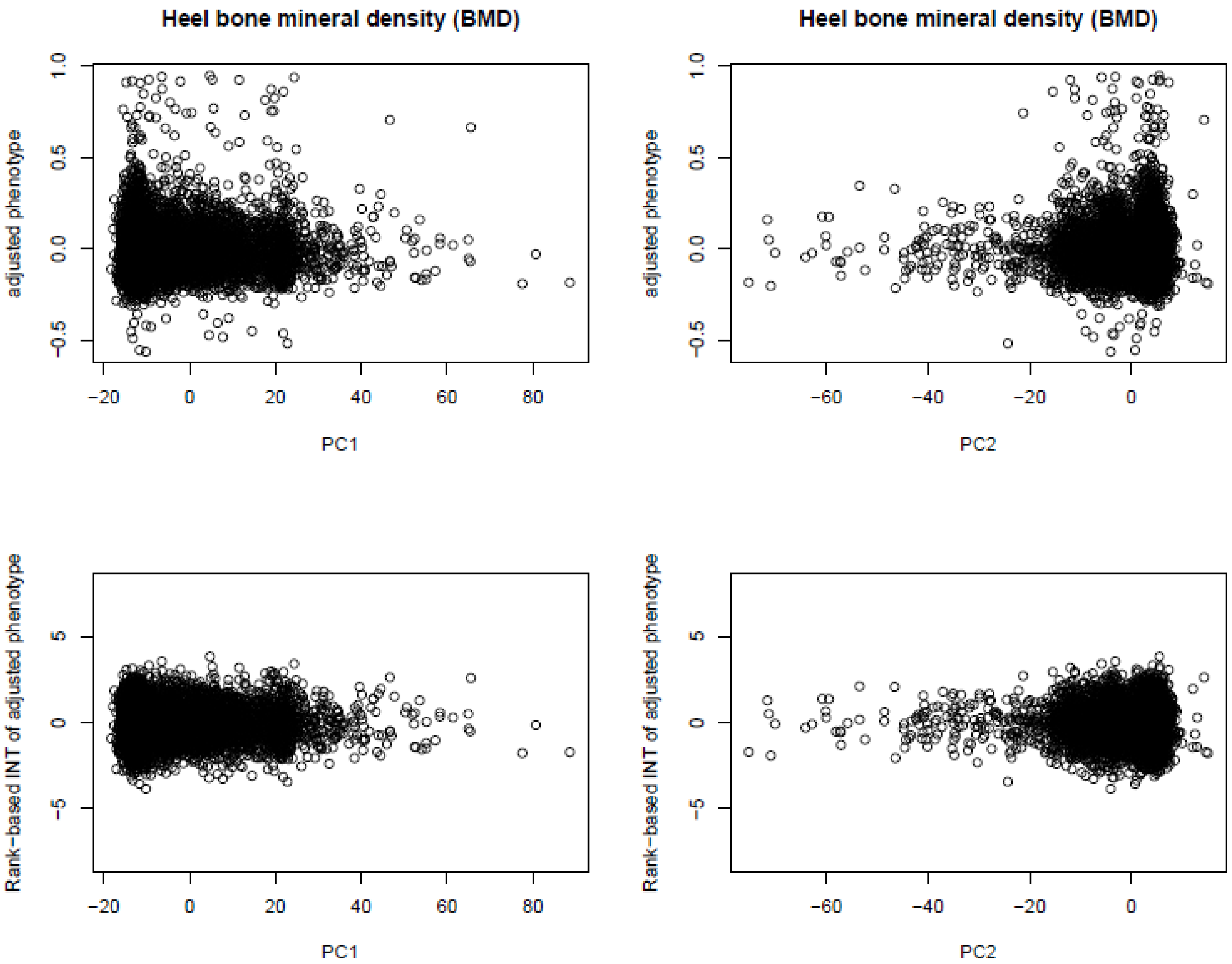


**Figure S7. Phenotypic heteroscedasticity of the trait "heel BMD" before and after rank-based INT.**

The phenotypes were adjusted by basic and additional confounders of fixed effects.

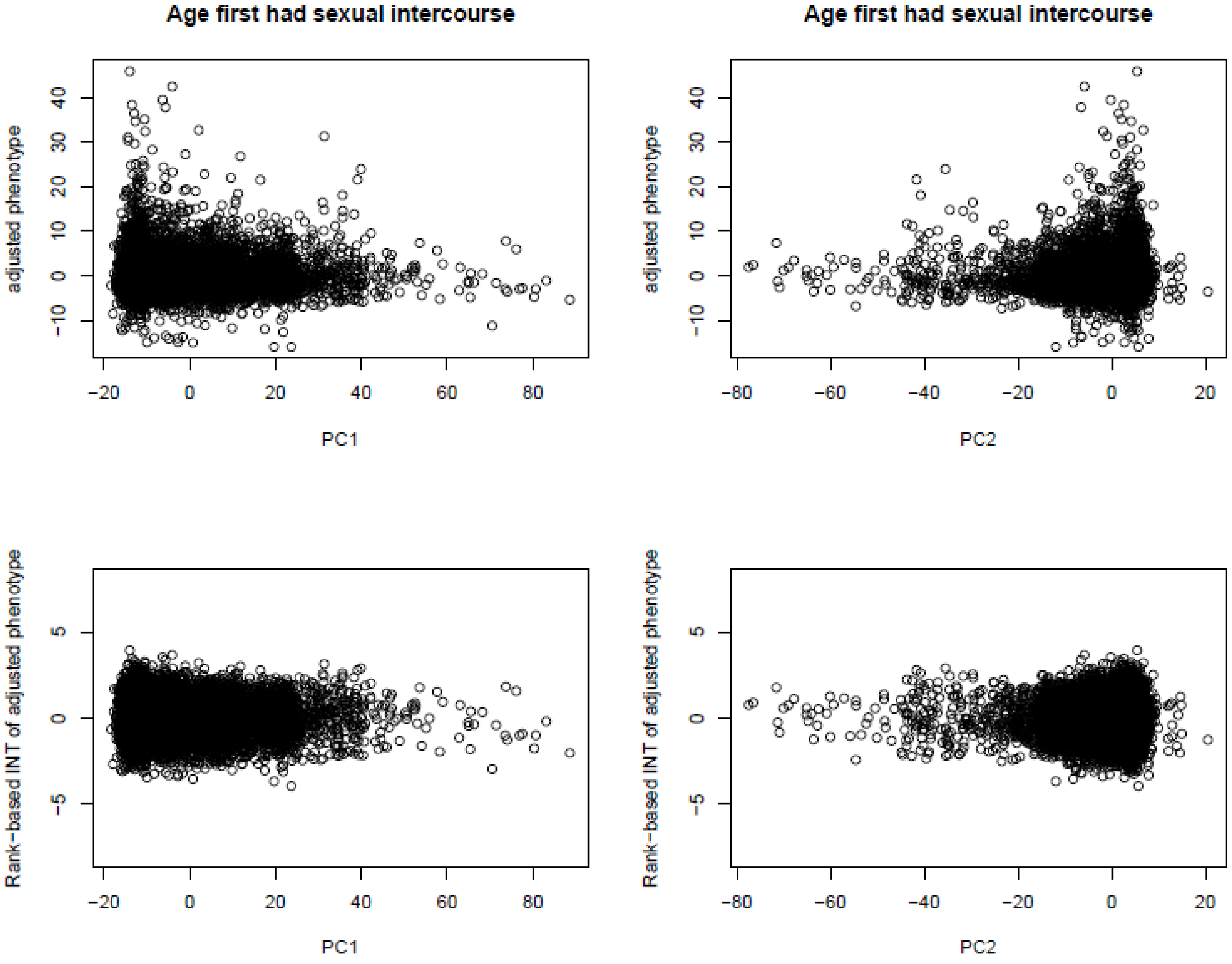


**Figure S8. Phenotypic heteroscedasticity of the trait "age first had sexual intercourse" before and after rank-based INT.** The phenotypes were adjusted by basic and additional confounders of fixed effects.

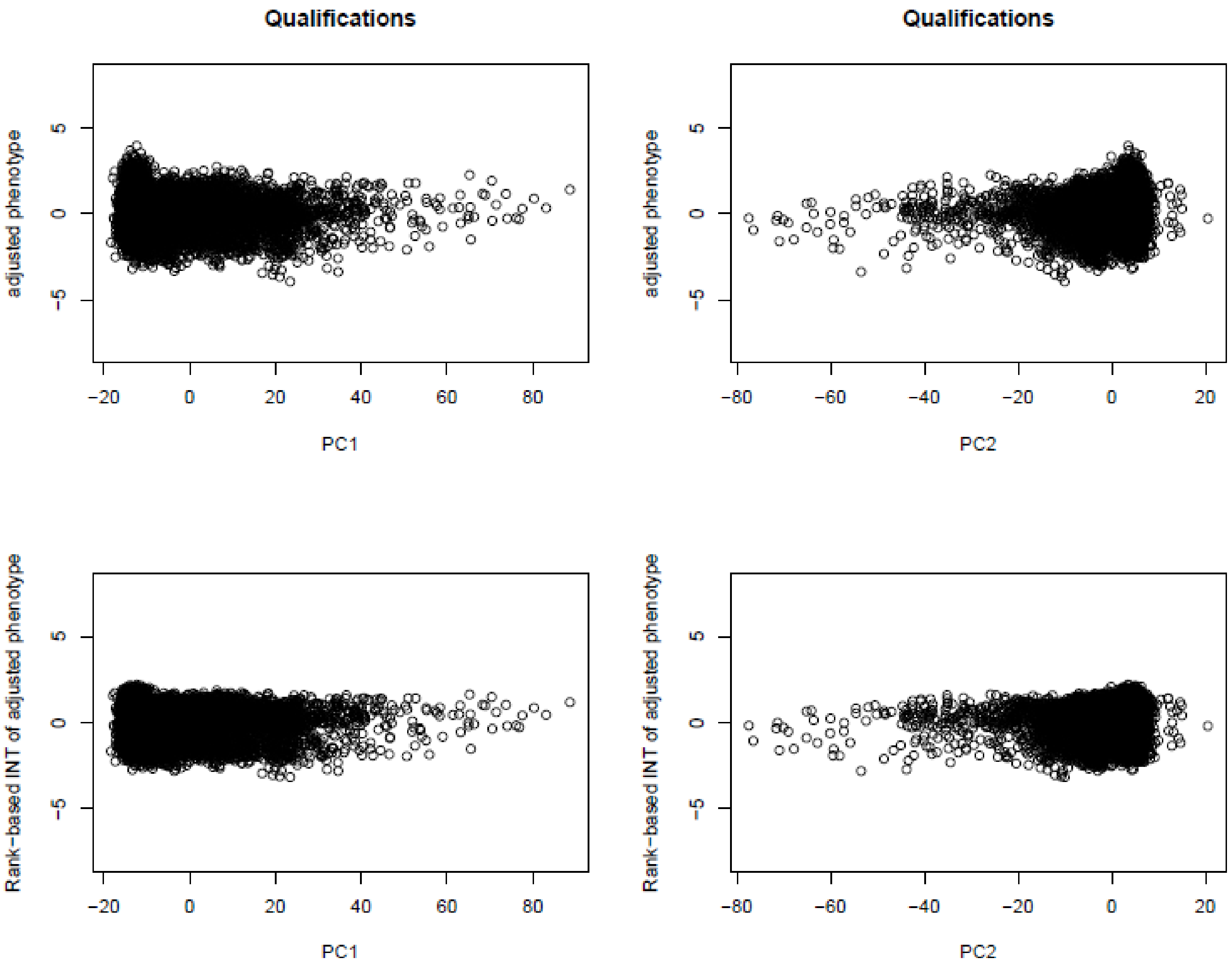


**Figure S9. Phenotypic heteroscedasticity of the trait "qualifications" before and after rank-based INT.**

The phenotypes were adjusted by basic and additional confounders of fixed effects.

**Table S1. The sample sizes of 199 UKBB variables of POP1, POP2 and POP1+POP2.** Here for each variable, non-NA number in POP1 is larger than 2,500, non-NA number in POP2 is larger than 2,500, non-NA number in POP1+POP2 is larger than 5,000.

| UKBB data field | UKBB variable description | Variable value type | Non-NA No. in POP1 | Non-NA No. in POP2 | Non-NA No. in POP1+POP2 |
|---|---|---|---|---|---|
| 19 | Heel ultrasound method | Unordered categorical | 4997 | 4045 | 9042 |
| 31 | Sex | Binary | 7487 | 6913 | 14400 |
| 34 | Year of birth | Continuous | 7487 | 6913 | 14400 |
| 48 | Waist circumference | Continuous | 7485 | 6888 | 14373 |
| 49 | Hip circumference | Continuous | 7487 | 6890 | 14377 |
| 50 | Standing height | Continuous | 7487 | 6886 | 14373 |
| 52 | Month of birth | Unordered categorical | 7487 | 6913 | 14400 |
| 54 | UK Biobank assessment centre | Unordered categorical | 7487 | 6913 | 14400 |
| 78 | Heel bone mineral density (BMD) T-score, automated | Continuous | 4331 | 3442 | 7773 |
| 87 | Non-cancer illness year/age first occurred | Continuous | 5621 | 4875 | 10496 |
| 102 | Pulse rate, automated reading | Continuous | 6989 | 6608 | 13597 |
| 120 | Birth weight known | Unordered categorical | 7487 | 6909 | 14396 |
| 132 | Job code | Unordered categorical | 4869 | 5105 | 9974 |
| 134 | Number of self-reported cancers | Continuous | 7487 | 6909 | 14396 |
| 135 | Number of self-reported non-cancer illnesses | Continuous | 7487 | 6909 | 14396 |
| 189 | Townsend deprivation index at recruitment | Continuous | 7487 | 6906 | 14393 |
| 403 | Number of times snap-button pressed | Continuous | 7469 | 6850 | 14319 |
| 404 | Duration to first press of snap-button in each round | Continuous | 3116 | 2812 | 5928 |
| 670 | Type of accommodation lived in | Unordered categorical | 7487 | 6913 | 14400 |
| 680 | Own or rent accommodation lived in | Unordered categorical | 7469 | 6886 | 14355 |
| 699 | Length of time at current address | Continuous | 7487 | 6913 | 14400 |
| 709 | Number in household | Continuous | 7469 | 6886 | 14355 |
| 728 | Number of vehicles in household | Ordered categorical | 7469 | 6886 | 14355 |
| 738 | Average total household income before tax | Ordered categorical | 7469 | 6879 | 14348 |
| 757 | Time employed in main current job | Continuous | 4220 | 4635 | 8855 |
| 767 | Length of working week for main job | Continuous | 4220 | 4635 | 8855 |
| 777 | Frequency of travelling from home to job workplace | Continuous | 4220 | 4631 | 8851 |
| 796 | Distance between home and job workplace | Continuous | 3953 | 4196 | 8149 |

| | | | | | |
|---|---|---|---|---|---|
| 806 | Job involves mainly walking or standing | Ordered categorical | 4220 | 4635 | 8855 |
| 816 | Job involves heavy manual or physical work | Ordered categorical | 4220 | 4635 | 8855 |
| 826 | Job involves shift work | Ordered categorical | 4220 | 4635 | 8855 |
| 845 | Age completed full time education | Continuous | 5241 | 2953 | 8194 |
| 1110 | Length of mobile phone use | Ordered categorical | 7487 | 6913 | 14400 |
| 1120 | Weekly usage of mobile phone in last 3 months | Ordered categorical | 6363 | 6062 | 12425 |
| 1130 | Hands-free device/speakerphone use with mobile phone in last 3 month | Ordered categorical | 6363 | 6062 | 12425 |
| 1140 | Difference in mobile phone use compared to two years previously | Ordered categorical | 6363 | 6062 | 12425 |
| 1150 | Usual side of head for mobile phone use | Binary | 6363 | 6062 | 12425 |
| 1160 | Sleep duration | Continuous | 7487 | 6913 | 14400 |
| 1170 | Getting up in morning | Ordered categorical | 7487 | 6906 | 14393 |
| 1180 | Morning/evening person (chronotype) | Ordered categorical | 7487 | 6906 | 14393 |
| 1190 | Nap during day | Ordered categorical | 7487 | 6913 | 14400 |
| 1200 | Sleeplessness / insomnia | Ordered categorical | 7487 | 6913 | 14400 |
| 1210 | Snoring | Binary | 7487 | 6913 | 14400 |
| 1220 | Daytime dozing / sleeping (narcolepsy) | Ordered categorical | 7487 | 6913 | 14400 |
| 1239 | Current tobacco smoking | Ordered categorical | 7487 | 6913 | 14400 |
| 1249 | Past tobacco smoking | Ordered categorical | 6574 | 6219 | 12793 |
| 1259 | Smoking/smokers in household | Unordered categorical | 6574 | 6212 | 12786 |
| 1269 | Exposure to tobacco smoke at home | Continuous | 6574 | 6219 | 12793 |
| 1279 | Exposure to tobacco smoke outside home | Continuous | 6574 | 6219 | 12793 |
| 1289 | Cooked vegetable intake | Continuous | 7487 | 6913 | 14400 |
| 1299 | Salad / raw vegetable intake | Continuous | 7487 | 6913 | 14400 |
| 1309 | Fresh fruit intake | Continuous | 7487 | 6913 | 14400 |
| 1319 | Dried fruit intake | Continuous | 7487 | 6913 | 14400 |
| 1329 | Oily fish intake | Ordered categorical | 7487 | 6913 | 14400 |
| 1339 | Non-oily fish intake | Ordered categorical | 7487 | 6913 | 14400 |
| 1349 | Processed meat intake | Ordered categorical | 7487 | 6913 | 14400 |
| 1359 | Poultry intake | Ordered categorical | 7487 | 6913 | 14400 |
| 1369 | Beef intake | Ordered categorical | 7487 | 6913 | 14400 |
| 1379 | Lamb/mutton intake | Ordered categorical | 7487 | 6913 | 14400 |
| 1389 | Pork intake | Ordered categorical | 7487 | 6913 | 14400 |
| 1408 | Cheese intake | Ordered categorical | 7306 | 6694 | 14000 |

| | | | | | |
|---|---|---|---|---|---|
| 1418 | Milk type used | Unordered categorical | 7487 | 6913 | 14400 |
| 1428 | Spread type | Unordered categorical | 7487 | 6906 | 14393 |
| 1438 | Bread intake | Continuous | 7487 | 6906 | 14393 |
| 1448 | Bread type | Unordered categorical | 7259 | 6596 | 13855 |
| 1458 | Cereal intake | Continuous | 7487 | 6913 | 14400 |
| 1468 | Cereal type | Unordered categorical | 6147 | 4944 | 11091 |
| 1478 | Salt added to food | Ordered categorical | 7487 | 6913 | 14400 |
| 1488 | Tea intake | Continuous | 7487 | 6913 | 14400 |
| 1498 | Coffee intake | Continuous | 7487 | 6913 | 14400 |
| 1508 | Coffee type | Unordered categorical | 5850 | 5744 | 11594 |
| 1518 | Hot drink temperature | Ordered categorical | 7487 | 6913 | 14400 |
| 1528 | Water intake | Continuous | 7487 | 6913 | 14400 |
| 1538 | Major dietary changes in the last 5 years | Unordered categorical | 7487 | 6913 | 14400 |
| 1548 | Variation in diet | Ordered categorical | 7487 | 6906 | 14393 |
| 1558 | Alcohol intake frequency. | Ordered categorical | 7487 | 6913 | 14400 |
| 1568 | Average weekly red wine intake | Continuous | 5272 | 4205 | 9477 |
| 1578 | Average weekly champagne plus white wine intake | Continuous | 5272 | 4205 | 9477 |
| 1588 | Average weekly beer plus cider intake | Continuous | 5272 | 4205 | 9477 |
| 1598 | Average weekly spirits intake | Continuous | 5272 | 4205 | 9477 |
| 1608 | Average weekly fortified wine intake | Continuous | 5272 | 4205 | 9477 |
| 1618 | Alcohol usually taken with meals | Ordered categorical | 5868 | 5138 | 11006 |
| 1628 | Alcohol intake versus 10 years previously | Ordered categorical | 6952 | 6293 | 13245 |
| 1647 | Country of birth (UK/elsewhere) | Unordered categorical | 7487 | 6913 | 14400 |
| 1677 | Breastfed as a baby | Binary | 7487 | 6913 | 14400 |
| 1687 | Comparative body size at age 10 | Ordered categorical | 7487 | 6913 | 14400 |
| 1697 | Comparative height size at age 10 | Ordered categorical | 7487 | 6913 | 14400 |
| 1707 | Handedness (chirality/laterality) | Binary | 7487 | 6913 | 14400 |
| 1717 | Skin colour | Ordered categorical | 7487 | 6913 | 14400 |
| 1747 | Hair colour (natural, before greying) | Unordered categorical | 7487 | 6913 | 14400 |
| 1767 | Adopted as a child | Binary | 7487 | 6913 | 14400 |
| 1777 | Part of a multiple birth | Binary | 7374 | 6823 | 14197 |
| 1787 | Maternal smoking around birth | Binary | 7374 | 6823 | 14197 |
| 1920 | Mood swings | Binary | 7487 | 6913 | 14400 |
| 1930 | Miserableness | Binary | 7487 | 6913 | 14400 |

| | | | | | |
|---|---|---|---|---|---|
| 1940 | Irritability | Binary | 7487 | 6913 | 14400 |
| 1950 | Sensitivity / hurt feelings | Binary | 7487 | 6913 | 14400 |
| 1960 | Fed-up feelings | Binary | 7487 | 6913 | 14400 |
| 1970 | Nervous feelings | Binary | 7487 | 6913 | 14400 |
| 1980 | Worrier / anxious feelings | Binary | 7487 | 6913 | 14400 |
| 1990 | Tense / 'highly strung' | Binary | 7487 | 6913 | 14400 |
| 2000 | Worry too long after embarrassment | Binary | 7487 | 6913 | 14400 |
| 2010 | Suffer from 'nerves' | Binary | 7487 | 6913 | 14400 |
| 2020 | Loneliness, isolation | Binary | 7487 | 6913 | 14400 |
| 2030 | Guilty feelings | Binary | 7487 | 6913 | 14400 |
| 2040 | Risk taking | Binary | 7487 | 6913 | 14400 |
| 2050 | Frequency of depressed mood in last 2 weeks | Ordered categorical | 7487 | 6913 | 14400 |
| 2060 | Frequency of unenthusiasm / disinterest in last 2 weeks | Ordered categorical | 7487 | 6913 | 14400 |
| 2070 | Frequency of tenseness / restlessness in last 2 weeks | Ordered categorical | 7487 | 6913 | 14400 |
| 2080 | Frequency of tiredness / lethargy in last 2 weeks | Ordered categorical | 7487 | 6913 | 14400 |
| 2090 | Seen doctor (GP) for nerves, anxiety, tension or depression | Binary | 7487 | 6913 | 14400 |
| 2100 | Seen a psychiatrist for nerves, anxiety, tension or depression | Binary | 7487 | 6913 | 14400 |
| 2129 | Answered sexual history questions | Binary | 7487 | 6913 | 14400 |
| 2139 | Age first had sexual intercourse | Continuous | 6939 | 6173 | 13112 |
| 2149 | Lifetime number of sexual partners | Continuous | 6866 | 6130 | 12996 |
| 2159 | Ever had same-sex intercourse | Binary | 6866 | 6130 | 12996 |
| 2207 | Wears glasses or contact lenses | Binary | 7487 | 6913 | 14400 |
| 2217 | Age started wearing glasses or contact lenses | Continuous | 6692 | 5654 | 12346 |
| 2227 | Other eye problems | Binary | 7487 | 6913 | 14400 |
| 2237 | Plays computer games | Ordered categorical | 7487 | 6913 | 14400 |
| 2247 | Hearing difficulty/problems | Binary | 7487 | 6906 | 14393 |
| 2257 | Hearing difficulty/problems with background noise | Binary | 7484 | 6911 | 14395 |
| 2375 | Relative age of first facial hair | Ordered categorical | 3502 | 2768 | 6270 |
| 2385 | Relative age voice broke | Ordered categorical | 3502 | 2768 | 6270 |
| 2395 | Hair/balding pattern | Ordered categorical | 3502 | 2768 | 6270 |
| 2405 | Number of children fathered | Continuous | 3502 | 2768 | 6270 |
| 2443 | Diabetes diagnosed by doctor | Binary | 7487 | 6913 | 14400 |
| 2453 | Cancer diagnosed by doctor | Binary | 7487 | 6913 | 14400 |
| 2463 | Fractured/broken bones in last 5 years | Binary | 7487 | 6913 | 14400 |

| | | | | | |
|---|---|---|---|---|---|
| 2473 | Other serious medical condition/disability diagnosed by doctor | Binary | 7487 | 6913 | 14400 |
| 2492 | Taking other prescription medications | Binary | 7487 | 6913 | 14400 |
| 2654 | Non-butter spread type details | Unordered categorical | 3970 | 2509 | 6479 |
| 2664 | Reason for reducing amount of alcohol drunk | Unordered categorical | 3075 | 2543 | 5618 |
| 2714 | Age when periods started (menarche) | Continuous | 3985 | 4142 | 8127 |
| 2734 | Number of live births | Continuous | 3985 | 4142 | 8127 |
| 2744 | Birth weight of first child | Continuous | 3200 | 3080 | 6280 |
| 2784 | Ever taken oral contraceptive pill | Binary | 3985 | 4142 | 8127 |
| 3081 | Foot measured for bone density | Binary | 4964 | 3984 | 8948 |
| 3082 | Fractured heel | Binary | 4964 | 3984 | 8948 |
| 3140 | Pregnant | Binary | 3985 | 4131 | 8116 |
| 3143 | Ankle spacing width | Continuous | 4331 | 3442 | 7773 |
| 3144 | Heel Broadband ultrasound attenuation, direct entry | Continuous | 4331 | 3442 | 7773 |
| 3147 | Heel quantitative ultrasound index (QUI), direct entry | Continuous | 4331 | 3442 | 7773 |
| 3148 | Heel bone mineral density (BMD) | Continuous | 4327 | 3442 | 7769 |
| 3393 | Hearing aid user | Binary | 4455 | 4457 | 8912 |
| 4079 | Diastolic blood pressure, automated reading | Continuous | 6989 | 6608 | 13597 |
| 4080 | Systolic blood pressure, automated reading | Continuous | 6988 | 6608 | 13596 |
| § 6138 | Qualifications | Ordered categorical | 7487 | 6906 | 14393 |
| 6141 | How are people in household related to participant | Unordered categorical | 6057 | 5485 | 11542 |
| 6142 | Current employment status | Unordered categorical | 7487 | 6913 | 14400 |
| 6143 | Transport type for commuting to job workplace | Unordered categorical | 3953 | 4196 | 8149 |
| 6144 | Never eat eggs, dairy, wheat, sugar | Unordered categorical | 7487 | 6906 | 14393 |
| 6145 | Illness, injury, bereavement, stress in last 2 years | Unordered categorical | 7487 | 6906 | 14393 |
| 6150 | Vascular/heart problems diagnosed by doctor | Unordered categorical | 7487 | 6913 | 14400 |
| 6152 | Blood clot, DVT, bronchitis, emphysema, asthma, rhinitis, eczema, allergy diagnosed by doctor | Unordered categorical | 7487 | 6913 | 14400 |
| 6153 | Medication for cholesterol, blood pressure, diabetes, or take exogenous hormones | Unordered categorical | 3985 | 4138 | 8123 |
| 6154 | Medication for pain relief, constipation, heartburn | Unordered categorical | 7487 | 6906 | 14393 |
| 6155 | Vitamin and mineral supplements | Unordered categorical | 7487 | 6906 | 14393 |
| 6177 | Medication for cholesterol, blood pressure or diabetes | Unordered categorical | 3502 | 2768 | 6270 |
| 6179 | Mineral and other dietary supplements | Unordered categorical | 7487 | 6906 | 14393 |
| 20002 | Non-cancer illness code, self-reported | Unordered categorical | 5621 | 4875 | 10496 |

| | | | | | |
|---|---|---|---|---|---|
| 20008 | Interpolated Year when non-cancer illness first diagnosed | Continuous | 5621 | 4875 | 10496 |
| 20009 | Interpolated Age of participant when non-cancer illness first diagnosed | Continuous | 5621 | 4875 | 10496 |
| 20013 | Method of recording time when non-cancer illness first diagnosed | Unordered categorical | 5621 | 4875 | 10496 |
| 20015 | Sitting height | Continuous | 7486 | 6881 | 14367 |
| 20022 | Birth weight | Continuous | 4273 | 3590 | 7863 |
| 20023 | Mean time to correctly identify matches | Continuous | 7440 | 6776 | 14216 |
| 20116 | Smoking status | Ordered categorical | 7487 | 6913 | 14400 |
| 20117 | Alcohol drinker status | Ordered categorical | 7487 | 6913 | 14400 |
| 20127 | Neuroticism score | Continuous | 6140 | 5065 | 11205 |
| 20160 | Ever smoked | Binary | 7487 | 6281 | 13768 |
| 21000 | Ethnic background | Unordered categorical | 7487 | 6913 | 14400 |
| 21001 | Body mass index (BMI) | Continuous | 7487 | 6878 | 14365 |
| 21002 | Weight | Continuous | 7487 | 6879 | 14366 |
| 21003 | Age when attended assessment centre | Continuous | 7487 | 6913 | 14400 |
| 21022 | Age at recruitment | Continuous | 7487 | 6913 | 14400 |
| 22000 | Genotype measurement batch | Unordered categorical | 7487 | 6913 | 14400 |
| 24003 | Nitrogen dioxide air pollution; 2010 | Continuous | 7379 | 6677 | 14056 |
| 24004 | Nitrogen oxides air pollution; 2010 | Continuous | 7379 | 6677 | 14056 |
| 24005 | Particulate matter air pollution (pm10); 2010 | Continuous | 6801 | 6392 | 13193 |
| 24006 | Particulate matter air pollution (pm2.5); 2010 | Continuous | 6801 | 6392 | 13193 |
| 24007 | Particulate matter air pollution (pm2.5) absorbance; 2010 | Continuous | 6801 | 6392 | 13193 |
| 24008 | Particulate matter air pollution 2.5-10um; 2010 | Continuous | 6801 | 6392 | 13193 |
| 24009 | Traffic intensity on the nearest road | Continuous | 7379 | 6677 | 14056 |
| 24010 | Inverse distance to the nearest road | Continuous | 7379 | 6677 | 14056 |
| 24011 | Traffic intensity on the nearest major road | Continuous | 7379 | 6677 | 14056 |
| 24012 | Inverse distance to the nearest major road | Continuous | 7379 | 6677 | 14056 |
| 24013 | Total traffic load on major roads | Continuous | 7379 | 6677 | 14056 |
| 24014 | Close to major road | Binary | 7379 | 6677 | 14056 |
| 24015 | Sum of road length of major roads within 100m | Continuous | 7379 | 6677 | 14056 |
| 24016 | Nitrogen dioxide air pollution; 2005 | Continuous | 7379 | 6677 | 14056 |
| 24017 | Nitrogen dioxide air pollution; 2006 | Continuous | 7379 | 6677 | 14056 |
| 24018 | Nitrogen dioxide air pollution; 2007 | Continuous | 7379 | 6677 | 14056 |
| 24019 | Particulate matter air pollution (pm10); 2007 | Continuous | 7356 | 6674 | 14030 |

| | | | | | |
|---|---|---|---|---|---|
| 24020 | Average daytime sound level of noise pollution | Continuous | 7379 | 6677 | 14056 |
| 24021 | Average evening sound level of noise pollution | Continuous | 7379 | 6677 | 14056 |
| 24022 | Average night-time sound level of noise pollution | Continuous | 7379 | 6677 | 14056 |
| 24023 | Average 16-hour sound level of noise pollution | Continuous | 7379 | 6677 | 14056 |
| 24024 | Average 24-hour sound level of noise pollution | Continuous | 7379 | 6677 | 14056 |

§: Qualifications (data field 6138) was reclassified and ordered as 1) none, 2) O-levels or CSEs, 3) A-levels, NVQ, HND, HNC or other professional qualification, and 4) college or university degree.

**Table S2. The use of 199 UKBB variables for current study.** 128 variables were selected as phenotypes. The other variables can be used as basic confounders, additional confounders or excluded.

| UKBB data field | UKBB variable description | UKBB category | Note |
|---|---|---|---|
| 31 | Sex | Baseline characteristics | Confounder (basic) |
| 34 | Year of birth | Baseline characteristics | Confounder (basic) |
| 52 | Month of birth | Baseline characteristics | Excluded |
| 189 | Townsend deprivation index at recruitment | Baseline characteristics | Confounder |
| 21022 | Age at recruitment | Baseline characteristics | Confounder (basic) |
| 22000 | Genotype measurement batch | Genomics - Genotyping process and sample QC | Confounder (basic) |
| 54 | UK Biobank assessment centre | Reception - Recruitment | Confounder (basic) |
| 21003 | Age when attended assessment centre | Reception - Recruitment | Confounder (basic), same as "21022" |
| 48 | Waist circumference | Physical measures - Anthropometry | **Phenotype** |
| 49 | Hip circumference | Physical measures - Anthropometry | **Phenotype** |
| 50 | Standing height | Physical measures - Anthropometry | **Phenotype** |
| 20015 | Sitting height | Physical measures - Anthropometry | **Phenotype** |
| 21001 | Body mass index (BMI) | Physical measures - Anthropometry | **Phenotype** |
| 21002 | Weight | Physical measures - Anthropometry | **Phenotype** |
| 102 | Pulse rate, automated reading | Physical measures - Blood pressure | **Phenotype** |
| 4079 | Diastolic blood pressure, automated reading | Physical measures - Blood pressure | **Phenotype** |
| 4080 | Systolic blood pressure, automated reading | Physical measures - Blood pressure | **Phenotype** |
| 19 | Heel ultrasound method | Physical measures - Bone-densitometry of heel | Confounder |
| 78 | Heel bone mineral density (BMD) T-score, automated | Physical measures - Bone-densitometry of heel | **Phenotype** |
| 3081 | Foot measured for bone density | Physical measures - Bone-densitometry of heel | Confounder |
| 3082 | Fractured heel | Physical measures - Bone-densitometry of heel | Confounder |
| 3143 | Ankle spacing width | Physical measures - Bone-densitometry of heel | **Phenotype** |
| 3144 | Heel Broadband ultrasound attenuation, direct entry | Physical measures - Bone-densitometry of heel | **Phenotype** |
| 3147 | Heel quantitative ultrasound index (QUI), direct entry | Physical measures - Bone-densitometry of heel | **Phenotype** |
| 3148 | Heel bone mineral density (BMD) | Physical measures - Bone-densitometry of heel | **Phenotype** |
| 1558 | Alcohol intake frequency. | Lifestyle and environment - Alcohol | **Phenotype** |
| 1568 | Average weekly red wine intake | Lifestyle and environment - Alcohol | **Phenotype** |
| 1578 | Average weekly champagne plus white wine intake | Lifestyle and environment - Alcohol | **Phenotype** |

| | | | |
|---|---|---|---|
| 1588 | Average weekly beer plus cider intake | Lifestyle and environment - Alcohol | **Phenotype** |
| 1598 | Average weekly spirits intake | Lifestyle and environment - Alcohol | **Phenotype** |
| 1608 | Average weekly fortified wine intake | Lifestyle and environment - Alcohol | **Phenotype** |
| 1618 | Alcohol usually taken with meals | Lifestyle and environment - Alcohol | **Phenotype** |
| 1628 | Alcohol intake versus 10 years previously | Lifestyle and environment - Alcohol | **Phenotype** |
| 2664 | Reason for reducing amount of alcohol drunk | Lifestyle and environment - Alcohol | Confounder |
| 20117 | Alcohol drinker status | Lifestyle and environment - Alcohol | **Phenotype** |
| 6144 | Never eat eggs, dairy, wheat, sugar | Lifestyle and environment - Diet | Confounder |
| 1289 | Cooked vegetable intake | Lifestyle and environment - Diet | **Phenotype** |
| 1299 | Salad / raw vegetable intake | Lifestyle and environment - Diet | **Phenotype** |
| 1309 | Fresh fruit intake | Lifestyle and environment - Diet | **Phenotype** |
| 1319 | Dried fruit intake | Lifestyle and environment - Diet | **Phenotype** |
| 1329 | Oily fish intake | Lifestyle and environment - Diet | **Phenotype** |
| 1339 | Non-oily fish intake | Lifestyle and environment - Diet | **Phenotype** |
| 1349 | Processed meat intake | Lifestyle and environment - Diet | **Phenotype** |
| 1359 | Poultry intake | Lifestyle and environment - Diet | **Phenotype** |
| 1369 | Beef intake | Lifestyle and environment - Diet | **Phenotype** |
| 1379 | Lamb/mutton intake | Lifestyle and environment - Diet | **Phenotype** |
| 1389 | Pork intake | Lifestyle and environment - Diet | **Phenotype** |
| 1408 | Cheese intake | Lifestyle and environment - Diet | **Phenotype** |
| 1418 | Milk type used | Lifestyle and environment - Diet | Confounder |
| 1428 | Spread type | Lifestyle and environment - Diet | Confounder |
| 1438 | Bread intake | Lifestyle and environment - Diet | **Phenotype** |
| 1448 | Bread type | Lifestyle and environment - Diet | Confounder |
| 1458 | Cereal intake | Lifestyle and environment - Diet | **Phenotype** |
| 1468 | Cereal type | Lifestyle and environment - Diet | Confounder |
| 1478 | Salt added to food | Lifestyle and environment - Diet | **Phenotype** |
| 1488 | Tea intake | Lifestyle and environment - Diet | **Phenotype** |
| 1498 | Coffee intake | Lifestyle and environment - Diet | **Phenotype** |
| 1508 | Coffee type | Lifestyle and environment - Diet | Confounder |
| 1518 | Hot drink temperature | Lifestyle and environment - Diet | **Phenotype** |
| 1528 | Water intake | Lifestyle and environment - Diet | **Phenotype** |
| 1538 | Major dietary changes in the last 5 years | Lifestyle and environment - Diet | Confounder |
| 1548 | Variation in diet | Lifestyle and environment - Diet | **Phenotype** |

| | | | |
|---|---|---|---|
| 2654 | Non-butter spread type details | Lifestyle and environment - Diet | Confounder |
| 1110 | Length of mobile phone use | Lifestyle and environment - Electronic device use | **Phenotype** |
| 1120 | Weekly usage of mobile phone in last 3 months | Lifestyle and environment - Electronic device use | **Phenotype** |
| 1130 | Hands-free device/speakerphone use with mobile phone in last 3 month | Lifestyle and environment - Electronic device use | **Phenotype** |
| 1140 | Difference in mobile phone use compared to two years previously | Lifestyle and environment - Electronic device use | **Phenotype** |
| 1150 | Usual side of head for mobile phone use | Lifestyle and environment - Electronic device use | **Phenotype** |
| 2237 | Plays computer games | Lifestyle and environment - Electronic device use | **Phenotype** |
| 2129 | Answered sexual history questions | Lifestyle and environment - Sexual factors | Confounder |
| 2139 | Age first had sexual intercourse | Lifestyle and environment - Sexual factors | **Phenotype** |
| 2149 | Lifetime number of sexual partners | Lifestyle and environment - Sexual factors | **Phenotype** |
| 2159 | Ever had same-sex intercourse | Lifestyle and environment - Sexual factors | **Phenotype** |
| 1160 | Sleep duration | Lifestyle and environment - Sleep | **Phenotype** |
| 1170 | Getting up in morning | Lifestyle and environment - Sleep | **Phenotype** |
| 1180 | Morning/evening person (chronotype) | Lifestyle and environment - Sleep | **Phenotype** |
| 1190 | Nap during day | Lifestyle and environment - Sleep | **Phenotype** |
| 1200 | Sleeplessness / insomnia | Lifestyle and environment - Sleep | **Phenotype** |
| 1210 | Snoring | Lifestyle and environment - Sleep | **Phenotype** |
| 1220 | Daytime dozing / sleeping (narcolepsy) | Lifestyle and environment - Sleep | **Phenotype** |
| 1239 | Current tobacco smoking | Lifestyle and environment - Smoking | **Phenotype** |
| 1249 | Past tobacco smoking | Lifestyle and environment - Smoking | **Phenotype** |
| 1259 | Smoking/smokers in household | Lifestyle and environment - Smoking | Confounder |
| 1269 | Exposure to tobacco smoke at home | Lifestyle and environment - Smoking | Confounder |
| 1279 | Exposure to tobacco smoke outside home | Lifestyle and environment - Smoking | Confounder |
| 20116 | Smoking status | Lifestyle and environment - Smoking | **Phenotype** |
| 20160 | Ever smoked | Lifestyle and environment - Smoking | **Phenotype** |
| 1717 | Skin colour | Lifestyle and environment - Sun exposure | **Phenotype** |
| 1747 | Hair colour (natural, before greying) | Lifestyle and environment - Sun exposure | Confounder |
| 1647 | Country of birth (UK/elsewhere) | Early life factors | Confounder |
| 1677 | Breastfed as a baby | Early life factors | **Phenotype** |
| 1687 | Comparative body size at age 10 | Early life factors | **Phenotype** |
| 1697 | Comparative height size at age 10 | Early life factors | **Phenotype** |
| 1707 | Handedness (chirality/laterality) | Early life factors | **Phenotype** |

| | | | |
|---|---|---|---|
| 1767 | Adopted as a child | Early life factors | **Phenotype** |
| 1777 | Part of a multiple birth | Early life factors | **Phenotype** |
| 1787 | Maternal smoking around birth | Early life factors | **Phenotype** |
| 20022 | Birth weight | Early life factors | **Phenotype** |
| 120 | Birth weight known | Early life factors | Confounder |
| 845 | Age completed full time education | Sociodemographics - Education | **Phenotype** |
| 6138 | Qualifications | Sociodemographics - Education | **Phenotype** (reorganized) |
| 132 | Job code | Sociodemographics - Employment | Confounder |
| 757 | Time employed in main current job | Sociodemographics - Employment | **Phenotype** |
| 767 | Length of working week for main job | Sociodemographics - Employment | **Phenotype** |
| 777 | Frequency of travelling from home to job workplace | Sociodemographics - Employment | **Phenotype** |
| 796 | Distance between home and job workplace | Sociodemographics - Employment | **Phenotype** |
| 806 | Job involves mainly walking or standing | Sociodemographics - Employment | **Phenotype** |
| 816 | Job involves heavy manual or physical work | Sociodemographics - Employment | **Phenotype** |
| 826 | Job involves shift work | Sociodemographics - Employment | **Phenotype** |
| 6142 | Current employment status | Sociodemographics - Employment | Confounder |
| 6143 | Transport type for commuting to job workplace | Sociodemographics - Employment | Confounder |
| 21000 | Ethnic background | Sociodemographics - Ethnicity | Have been used for sampling |
| 670 | Type of accommodation lived in | Sociodemographics - Household | Confounder |
| 680 | Own or rent accommodation lived in | Sociodemographics - Household | Confounder |
| 699 | Length of time at current address | Sociodemographics - Household | **Phenotype** |
| 709 | Number in household | Sociodemographics - Household | **Phenotype** |
| 728 | Number of vehicles in household | Sociodemographics - Household | **Phenotype** |
| 738 | Average total household income before tax | Sociodemographics - Household | **Phenotype** |
| 6141 | How are people in household related to participant | Sociodemographics - Household | Confounder |
| 2714 | Age when periods started (menarche) | Female-specific factors | **Phenotype** |
| 2734 | Number of live births | Female-specific factors | **Phenotype** |
| 2744 | Birth weight of first child | Female-specific factors | **Phenotype** |
| 2784 | Ever taken oral contraceptive pill | Female-specific factors | Confounder |
| 2375 | Relative age of first facial hair | Male-specific factors | **Phenotype** |
| 2385 | Relative age voice broke | Male-specific factors | **Phenotype** |
| 2395 | Hair/balding pattern | Male-specific factors | **Phenotype** |
| 2405 | Number of children fathered | Male-specific factors | **Phenotype** |
| 2207 | Wears glasses or contact lenses | Health and medical history - Eyesight | **Phenotype** |

| | | | |
|---|---|---|---|
| 2217 | Age started wearing glasses or contact lenses | Health and medical history - Eyesight | **Phenotype** |
| 2227 | Other eye problems | Health and medical history - Eyesight | **Phenotype** |
| 2247 | Hearing difficulty/problems | Health and medical history - Hearing | **Phenotype** |
| 2257 | Hearing difficulty/problems with background noise | Health and medical history - Hearing | **Phenotype** |
| 3393 | Hearing aid user | Health and medical history - Hearing | **Phenotype** |
| 2443 | Diabetes diagnosed by doctor | Health and medical history - Medical conditions | **Phenotype** |
| 2453 | Cancer diagnosed by doctor | Health and medical history - Medical conditions | **Phenotype** |
| 2463 | Fractured/broken bones in last 5 years | Health and medical history - Medical conditions | **Phenotype** |
| 2473 | Other serious medical condition/disability diagnosed by doctor | Health and medical history - Medical conditions | **Phenotype** |
| 6150 | Vascular/heart problems diagnosed by doctor | Health and medical history - Medical conditions | Confounder |
| 6152 | Blood clot, DVT, bronchitis, emphysema, asthma, rhinitis, eczema, allergy diagnosed by doctor | Health and medical history - Medical conditions | Confounder |
| 6153 | Medication for cholesterol, blood pressure, diabetes, or take exogenous hormones | Health and medical history - Medical conditions | Confounder |
| 2492 | Taking other prescription medications | Health and medical history - Medication | Confounder |
| 6155 | Vitamin and mineral supplements | Health and medical history - Medication | Confounder |
| 6177 | Medication for cholesterol, blood pressure or diabetes | Health and medical history - Medication | Confounder |
| 6179 | Mineral and other dietary supplements | Health and medical history - Medication | Confounder |
| 6154 | Medication for pain relief, constipation, heartburn | Health and medical history - Medication | Confounder |
| 403 | Number of times snap-button pressed | Cognitive function - Reaction time | **Phenotype** |
| 404 | Duration to first press of snap-button in each round | Cognitive function - Reaction time | **Phenotype** |
| 20023 | Mean time to correctly identify matches | Cognitive function - Reaction time | **Phenotype** |
| 1920 | Mood swings | Psychosocial factors - Mental health | **Phenotype** |
| 1930 | Miserableness | Psychosocial factors - Mental health | **Phenotype** |
| 1940 | Irritability | Psychosocial factors - Mental health | **Phenotype** |
| 1950 | Sensitivity / hurt feelings | Psychosocial factors - Mental health | **Phenotype** |
| 1960 | Fed-up feelings | Psychosocial factors - Mental health | **Phenotype** |
| 1970 | Nervous feelings | Psychosocial factors - Mental health | **Phenotype** |
| 1980 | Worrier / anxious feelings | Psychosocial factors - Mental health | **Phenotype** |
| 1990 | Tense / 'highly strung' | Psychosocial factors - Mental health | **Phenotype** |
| 2000 | Worry too long after embarrassment | Psychosocial factors - Mental health | **Phenotype** |
| 2010 | Suffer from 'nerves' | Psychosocial factors - Mental health | **Phenotype** |
| 2020 | Loneliness, isolation | Psychosocial factors - Mental health | **Phenotype** |

| | | | |
|---|---|---|---|
| 2030 | Guilty feelings | Psychosocial factors - Mental health | **Phenotype** |
| 2040 | Risk taking | Psychosocial factors - Mental health | **Phenotype** |
| 2050 | Frequency of depressed mood in last 2 weeks | Psychosocial factors - Mental health | **Phenotype** |
| 2060 | Frequency of unenthusiasm / disinterest in last 2 weeks | Psychosocial factors - Mental health | **Phenotype** |
| 2070 | Frequency of tenseness / restlessness in last 2 weeks | Psychosocial factors - Mental health | **Phenotype** |
| 2080 | Frequency of tiredness / lethargy in last 2 weeks | Psychosocial factors - Mental health | **Phenotype** |
| 2090 | Seen doctor (GP) for nerves, anxiety, tension or depression | Psychosocial factors - Mental health | **Phenotype** |
| 2100 | Seen a psychiatrist for nerves, anxiety, tension or depression | Psychosocial factors - Mental health | **Phenotype** |
| 6145 | Illness, injury, bereavement, stress in last 2 years | Psychosocial factors - Mental health | Confounder |
| 20127 | Neuroticism score | Psychosocial factors - Mental health | **Phenotype** |
| 87 | Non-cancer illness year/age first occurred | Verbal interview - Medical conditions | Excluded |
| 134 | Number of self-reported cancers | Verbal interview - Medical conditions | **Phenotype** |
| 135 | Number of self-reported non-cancer illnesses | Verbal interview - Medical conditions | **Phenotype** |
| 3140 | Pregnant | Verbal interview - Medical conditions | Confounder |
| 20002 | Non-cancer illness code, self-reported | Verbal interview - Medical conditions | Confounder |
| 20008 | Interpolated Year when non-cancer illness first diagnosed | Verbal interview - Medical conditions | Confounder |
| 20009 | Interpolated Age of participant when non-cancer illness first diagnosed | Verbal interview - Medical conditions | **Phenotype** |
| 20013 | Method of recording time when non-cancer illness first diagnosed | Verbal interview - Medical conditions | Confounder |
| 24003 | Nitrogen dioxide air pollution; 2010 | Residential air pollution | Confounder |
| 24004 | Nitrogen oxides air pollution; 2010 | Residential air pollution | Confounder |
| 24005 | Particulate matter air pollution (pm10); 2010 | Residential air pollution | Confounder |
| 24006 | Particulate matter air pollution (pm2.5); 2010 | Residential air pollution | Confounder |
| 24007 | Particulate matter air pollution (pm2.5) absorbance; 2010 | Residential air pollution | Confounder |
| 24008 | Particulate matter air pollution 2.5-10um; 2010 | Residential air pollution | Confounder |
| 24009 | Traffic intensity on the nearest road | Residential air pollution | Confounder |
| 24010 | Inverse distance to the nearest road | Residential air pollution | Confounder |
| 24011 | Traffic intensity on the nearest major road | Residential air pollution | Confounder |
| 24012 | Inverse distance to the nearest major road | Residential air pollution | Confounder |
| 24013 | Total traffic load on major roads | Residential air pollution | Confounder |
| 24014 | Close to major road | Residential air pollution | Confounder |

| | | | |
|---|---|---|---|
| 24015 | Sum of road length of major roads within 100m | Residential air pollution | Confounder |
| 24016 | Nitrogen dioxide air pollution; 2005 | Residential air pollution | Confounder |
| 24017 | Nitrogen dioxide air pollution; 2006 | Residential air pollution | Confounder |
| 24018 | Nitrogen dioxide air pollution; 2007 | Residential air pollution | Confounder |
| 24019 | Particulate matter air pollution (pm10); 2007 | Residential air pollution | Confounder |
| 24020 | Average daytime sound level of noise pollution | Residential noise pollution | Confounder |
| 24021 | Average evening sound level of noise pollution | Residential noise pollution | Confounder |
| 24022 | Average night-time sound level of noise pollution | Residential noise pollution | Confounder |
| 24023 | Average 16-hour sound level of noise pollution | Residential noise pollution | Confounder |
| 24024 | Average 24-hour sound level of noise pollution | Residential noise pollution | Confounder |

**Table S3**. **Seventy UKBB phenotypes with significant SNP-based heritability estimated by GREML model across POP1+POP2.** SE denotes standard error.

| Phenotypes | UKBB data field | $h^2_{SNP}$ (SE) |
|---|---|---|
| *Physical measures* - *Anthropometry*: | | |
| Waist circumference | 48 | 0.162(0.024) |
| Hip circumference | 49 | 0.187(0.025) |
| Standing height | 50 | 0.495(0.025) |
| Sitting height | 20015 | 0.335(0.025) |
| Body mass index (BMI) | 21001 | 0.204(0.025) |
| Weight | 21002 | 0.216(0.025) |
| *Physical measures* - *Blood pressure*: | | |
| Pulse rate, automated reading | 102 | 0.155(0.025) |
| Diastolic blood pressure, automated reading | 4079 | 0.122(0.025) |
| Systolic blood pressure, automated reading | 4080 | 0.134(0.025) |
| *Physical measures* - *Bone-densitometry of heel*: | | |
| Heel bone mineral density (BMD) T-score, automated | 78 | 0.268(0.045) |
| Ankle spacing width | 3143 | 0.176(0.044) |
| Heel Broadband ultrasound attenuation, direct entry | 3144 | 0.233(0.044) |
| Heel quantitative ultrasound index (QUI), direct entry | 3147 | 0.268(0.045) |
| Heel bone mineral density (BMD) | 3148 | 0.256(0.045) |
| *lifestyle and environment - Alcohol:* | | |
| Alcohol intake frequency | 1558 | 0.067(0.023) |
| Average weekly spirits intake | 1598 | 0.087(0.035) |
| *Lifestyle and environment - Diet:* | | |
| Fresh fruit intake | 1309 | 0.080(0.024) |
| Oily fish intake | 1329 | 0.083(0.023) |
| Beef intake | 1369 | 0.071(0.024) |
| Lamb/mutton intake | 1379 | 0.050(0.023) |
| Cereal intake | 1458 | 0.127(0.024) |
| Salt added to food | 1478 | 0.097(0.024) |
| Tea intake | 1488 | 0.054(0.023) |
| Hot drink temperature | 1518 | 0.118(0.024) |
| Water intake | 1528 | 0.121(0.024) |

| | | |
|---|---|---|
| Variation in diet | 1548 | 0.067(0.024) |
| *Lifestyle and environment - Electronic device use:* | | |
| Weekly usage of mobile phone in last 3 months | 1120 | 0.069(0.027) |
| Plays computer games | 2237 | 0.098(0.024) |
| *Lifestyle and environment - Sexual factors:* | | |
| Age first had sexual intercourse | 2139 | 0.103(0.027) |
| Lifetime number of sexual partners | 2149 | 0.170(0.031) |
| Ever had same-sex intercourse | 2159 | 0.056(0.026) |
| *Lifestyle and environment - Sleep*: | | |
| Sleep duration | 1160 | 0.074(0.023) |
| Getting up in morning | 1170 | 0.095(0.024) |
| Morning/evening person (chronotype) | 1180 | 0.086(0.026) |
| Nap during day | 1190 | 0.086(0.024) |
| Sleeplessness / insomnia | 1200 | 0.103(0.024) |
| *Lifestyle and environment - Smoking*: | | |
| Current tobacco smoking | 1239 | 0.045(0.023) |
| Past tobacco smoking | 1249 | 0.114(0.027) |
| Smoking status | 20116 | 0.141(0.024) |
| Ever smoked | 20160 | 0.133(0.025) |
| *Lifestyle and environment - Sun exposure*: | | |
| Skin colour | 1717 | 0.112(0.024) |
| *Early life factors*: | | |
| Comparative body size at age 10 | 1687 | 0.126(0.025) |
| Comparative height size at age 10 | 1697 | 0.243(0.025) |
| Birth weight | 20022 | 0.113(0.042) |
| *Female-specific factors*: | | |
| Age when periods started (menarche) | 2714 | 0.240(0.043) |
| Birth weight of first child | 2744 | 0.158(0.055) |
| *Male-specific factors*: | | |
| Relative age of first facial hair | 2375 | 0.187(0.055) |
| Relative age voice broke | 2385 | 0.165(0.058) |
| Hair/balding pattern | 2395 | 0.347(0.055) |
| *Psychosocial factors - Mental health*: | | |
| Mood swings | 1920 | 0.081(0.024) |
| Irritability | 1940 | 0.083(0.025) |

| | | |
|---|---|---|
| Sensitivity / hurt feelings | 1950 | 0.052(0.024) |
| Fed-up feelings | 1960 | 0.069(0.024) |
| Nervous feelings | 1970 | 0.066(0.024) |
| Worrier / anxious feelings | 1980 | 0.104(0.024) |
| Tense / 'highly strung' | 1990 | 0.069(0.025) |
| Worry too long after embarrassment | 2000 | 0.134(0.025) |
| Neuroticism score | 20127 | 0.170(0.031) |
| Guilty feelings | 2030 | 0.100(0.024) |
| Risk taking | 2040 | 0.094(0.025) |
| Frequency of depressed mood in last 2 weeks | 2050 | 0.092(0.025) |
| Frequency of unenthusiasm / disinterest | 2060 | 0.067(0.024) |
| Frequency of tenseness / restlessness | 2070 | 0.094(0.025) |
| Frequency of tiredness / lethargy in last 2 weeks | 2080 | 0.101(0.025) |
| Seen doctor (GP) for nerves, anxiety, tension or depression | 2090 | 0.064(0.023) |
| *Cognitive function - Reaction time*: | | |
| Number of times snap-button pressed | 403 | 0.053(0.024) |
| *Health and medical history - Eyesight*: | | |
| Age started wearing glasses or contact lenses | 2217 | 0.098(0.028) |
| *Sociodemographics - Education*: | | |
| Qualifications | 6138 | 0.162(0.024) |
| *Sociodemographics - Household*: | | |
| Number of vehicles in household | 728 | 0.048(0.023) |
| *Verbal interview - Medical conditions*: | | |
| Interpolated Age of participant when non-cancer illness first diagnosed | 20009 | 0.094(0.034) |

**Table S4**. **Fifty-eight UKBB phenotypes with significant SNP-based heritability estimated by GREML model across POP2+POP3**. SE denotes standard error.

| Phenotypes | UKBB data field | $h^2_{SNP}$ (SE) |
|---|---|---|
| *Physical measures - Anthropometry*: | | |
| Waist circumference | 48 | 0.205(0.024) |
| Hip circumference | 49 | 0.201(0.024) |
| Standing height | 50 | 0.493(0.024) |
| Sitting height | 20015 | 0.365(0.024) |
| Body mass index (BMI) | 21001 | 0.229(0.024) |
| Weight | 21002 | 0.251(0.024) |
| *Physical measures - Blood pressure*: | | |
| Pulse rate, automated reading | 102 | 0.121(0.024) |
| Diastolic blood pressure, automated reading | 4079 | 0.140(0.025) |
| Systolic blood pressure, automated reading | 4080 | 0.130(0.025) |
| *Physical measures - Bone-densitometry of heel*: | | |
| Heel bone mineral density (BMD) T-score, automated | 78 | 0.207(0.045) |
| Ankle spacing width | 3143 | 0.291(0.046) |
| Heel Broadband ultrasound attenuation, direct entry | 3144 | 0.193(0.045) |
| Heel quantitative ultrasound index (QUI), direct entry | 3147 | 0.207(0.045) |
| Heel bone mineral density (BMD) | 3148 | 0.212(0.045) |
| *lifestyle and environment - Alcohol:* | | |
| Alcohol intake frequency | 1558 | 0.110(0.023) |
| Average weekly spirits intake | 1598 | 0.076(0.035) |
| Alcohol intake versus 10 years previously | 1628 | 0.063(0.025) |
| *Lifestyle and environment - Diet:* | | |
| Fresh fruit intake | 1309 | 0.066(0.023) |
| Dried fruit intake | 1319 | 0.057(0.023) |
| Oily fish intake | 1329 | 0.049(0.022) |
| Non-oily fish intake | 1339 | 0.047(0.022) |
| Processed meat intake | 1349 | 0.085(0.023) |
| Salt added to food | 1478 | 0.098(0.023) |
| Hot drink temperature | 1518 | 0.087(0.023) |
| Water intake | 1528 | 0.105(0.023) |

| | | |
|---|---|---|
| Variation in diet | 1548 | 0.059(0.022) |
| *Lifestyle and environment - Electronic device use:* | | |
| Length of mobile phone use | 1110 | 0.074(0.023) |
| Hands-free device/speakerphone use with mobile phone in last 3 month | 1130 | 0.053(0.027) |
| Plays computer games | 2237 | 0.048(0.023) |
| *Lifestyle and environment - Sexual factors:* | | |
| Age first had sexual intercourse | 2139 | 0.074(0.025) |
| *Lifestyle and environment - Sleep*: | | |
| Sleep duration | 1160 | 0.081(0.023) |
| Getting up in morning | 1170 | 0.120(0.023) |
| Morning/evening person (chronotype) | 1180 | 0.138(0.025) |
| Nap during day | 1190 | 0.050(0.023) |
| Sleeplessness / insomnia | 1200 | 0.048(0.023) |
| *Lifestyle and environment - Smoking*: | | |
| Past tobacco smoking | 1249 | 0.088(0.026) |
| Smoking status | 20116 | 0.081(0.024) |
| *Lifestyle and environment - Sun exposure*: | | |
| Skin colour | 1717 | 0.163(0.024) |
| *Early life factors*: | | |
| Comparative body size at age 10 | 1687 | 0.177(0.024) |
| Comparative height size at age 10 | 1697 | 0.259(0.025) |
| *Female-specific factors*: | | |
| Age when periods started (menarche) | 2714 | 0.198(0.039) |
| Number of live births | 2734 | 0.119(0.037) |
| Birth weight of first child | 2744 | 0.141(0.053) |
| *Male-specific factors*: | | |
| Relative age of first facial hair | 2375 | 0.197(0.061) |
| Hair/balding pattern | 2395 | 0.426(0.059) |
| *Psychosocial factors - Mental health*: | | |
| Irritability | 1940 | 0.049(0.023) |
| Fed-up feelings | 1960 | 0.060(0.023) |
| Worry too long after embarrassment | 2000 | 0.048(0.024) |
| Neuroticism score | 20127 | 0.057(0.029) |
| Frequency of tenseness / restlessness | 2070 | 0.064(0.024) |

| | | |
|---|---|---|
| Frequency of tiredness / lethargy in last 2 weeks | 2080 | 0.048(0.023) |
| *Cognitive function - Reaction time*:<br>Duration to first press of snap-button in each round | <br>404 | <br>0.115(0.056) |
| *Health and medical history*:<br>Age started wearing glasses or contact lenses<br>Hearing difficulty/problems<br>Diabetes diagnosed by doctor | <br>2217<br>2247<br>2443 | <br>0.177(0.029)<br>0.067(0.024)<br>0.092(0.023) |
| *Sociodemographics - Education*:<br>Qualifications | <br>6138 | <br>0.162(0.024) |
| *Sociodemographics - Household*:<br>Length of time at current address<br>Average total household income before tax | <br>699<br>738 | <br>0.059(0.023)<br>0.057(0.026) |

**Table S5**. **Seventy UKBB phenotypes with significant SNP-based heritability estimated by GREML model across POP1+POP3**. SE denotes standard error.

| Phenotypes | UKBB data field | $h^2_{SNP}$ (SE) |
|---|---|---|
| *Physical measures - Anthropometry*: | | |
| Waist circumference | 48 | 0.182(0.023) |
| Hip circumference | 49 | 0.212(0.023) |
| Standing height | 50 | 0.548(0.023) |
| Sitting height | 20015 | 0.378(0.023) |
| Body mass index (BMI) | 21001 | 0.215(0.023) |
| Weight | 21002 | 0.240(0.023) |
| *Physical measures - Blood pressure*: | | |
| Pulse rate, automated reading | 102 | 0.148(0.024) |
| Diastolic blood pressure, automated reading | 4079 | 0.141(0.024) |
| Systolic blood pressure, automated reading | 4080 | 0.136(0.024) |
| *Physical measures - Bone-densitometry of heel*: | | |
| Heel bone mineral density (BMD) T-score, automated | 78 | 0.303(0.040) |
| Ankle spacing width | 3143 | 0.232(0.040) |
| Heel Broadband ultrasound attenuation, direct entry | 3144 | 0.307(0.040) |
| Heel quantitative ultrasound index (QUI), direct entry | 3147 | 0.303(0.040) |
| Heel bone mineral density (BMD) | 3148 | 0.308(0.041) |
| *lifestyle and environment - Alcohol:* | | |
| Alcohol intake frequency | 1558 | 0.093(0.022) |
| *Lifestyle and environment - Diet:* | | |
| Fresh fruit intake | 1309 | 0.048(0.021) |
| Dried fruit intake | 1319 | 0.044(0.022) |
| Oily fish intake | 1329 | 0.058(0.021) |
| Non-oily fish intake | 1339 | 0.045(0.021) |
| Processed meat intake | 1349 | 0.059(0.021) |
| Poultry intake | 1359 | 0.054(0.021) |
| Beef intake | 1369 | 0.046(0.022) |
| Lamb/mutton intake | 1379 | 0.046(0.021) |
| Cheese intake | 1408 | 0.048(0.022) |
| Cereal intake | 1458 | 0.065(0.021) |

| | | |
|---|---|---|
| Salt added to food<br>Hot drink temperature<br>Water intake | 1478<br>1518<br>1528 | 0.084(0.022)<br>0.090(0.022)<br>0.061(0.022) |
| *Lifestyle and environment - Electronic device use:*<br>Length of mobile phone use<br>Plays computer games | <br>1110<br>2237 | <br>0.070(0.022)<br>0.069(0.022) |
| *Lifestyle and environment - Sexual factors:*<br>Age first had sexual intercourse | <br>2139 | <br>0.146(0.025) |
| *Lifestyle and environment - Sleep*:<br>Sleep duration<br>Getting up in morning<br>Morning/evening person (chronotype)<br>Nap during day<br>Sleeplessness / insomnia<br>Snoring | <br>1160<br>1170<br>1180<br>1190<br>1200<br>1210 | <br>0.077(0.022)<br>0.119(0.022)<br>0.162(0.025)<br>0.094(0.022)<br>0.098(0.022)<br>0.076(0.024) |
| *Lifestyle and environment - Smoking*:<br>Current tobacco smoking<br>Past tobacco smoking<br>Smoking status<br>Ever smoked | <br>1239<br>1249<br>20116<br>20160 | <br>0.055(0.022)<br>0.092(0.025)<br>0.116(0.022)<br>0.093(0.023) |
| *Lifestyle and environment - Sun exposure*:<br>Skin colour | <br>1717 | <br>0.180(0.023) |
| *Early life factors*:<br>Comparative body size at age 10<br>Comparative height size at age 10<br>Birth weight | <br>1687<br>1697<br>20022 | <br>0.130(0.023)<br>0.279(0.023)<br>0.118(0.038) |
| *Female-specific factors*:<br>Age when periods started (menarche)<br>Number of live births | <br>2714<br>2734 | <br>0.183(0.038)<br>0.099(0.038) |
| *Male-specific factors*:<br>Relative age of first facial hair<br>Hair/balding pattern | <br>2375<br>2395 | <br>0.210(0.053)<br>0.436(0.053) |
| *Psychosocial factors - Mental health*:<br>Mood swings<br>Miserableness | <br>1920<br>1930 | <br>0.117(0.022)<br>0.060(0.022) |

| | | |
|---|---|---|
| Irritability | 1940 | 0.099(0.023) |
| Sensitivity / hurt feelings | 1950 | 0.055(0.023) |
| Fed-up feelings | 1960 | 0.099(0.022) |
| Nervous feelings | 1970 | 0.056(0.022) |
| Worrier / anxious feelings | 1980 | 0.095(0.022) |
| Tense / 'highly strung' | 1990 | 0.113(0.023) |
| Worry too long after embarrassment | 2000 | 0.077(0.023) |
| Neuroticism score | 20127 | 0.108(0.027) |
| Guilty feelings | 2030 | 0.069(0.022) |
| Risk taking | 2040 | 0.068(0.023) |
| Frequency of tiredness / lethargy in last 2 weeks | 2080 | 0.079(0.022) |
| Seen doctor (GP) for nerves, anxiety, tension or depression | 2090 | 0.046 (0.021) |
| *Cognitive function* - *Reaction time*: | | |
| Mean time to correctly identify matches | 20023 | 0.071(0.022) |
| *Health and medical history*: | | |
| Age started wearing glasses or contact lenses | 2217 | 0.141(0.026) |
| Diabetes diagnosed by doctor | 2443 | 0.049(0.021) |
| Fractured/broken bones in last 5 years | 2463 | 0.050(0.022) |
| *Sociodemographics* - *Education*: | | |
| Qualifications | 6138 | 0.173(0.022) |
| *Sociodemographics* - *Household*: | | |
| Average total household income before tax | 738 | 0.060(0.024) |
| *Verbal interview* - *Medical conditions*: | | |
| Number of self-reported non-cancer illnesses | 135 | 0.078(0.022) |

**Table S6**. **Genetic and residual covariances between 70 UKBB phenotypes and PC1/PC2 estimated by bivariate GREML model across POP1+POP2**. SE denotes standard error.

| UKBB data field | Phenotypes | Genetic covariance with PC1 (SE) | Genetic covariance with PC2 (SE) | Residual covariance with PC1 (SE) | Residual covariance with PC2 (SE) |
|---|---|---|---|---|---|
| | *Physical measures* - *Anthropometry*: | | | | |
| 48 | Waist circumference | 0.0041(0.0189) | 0.0007(0.017) | -0.0176(0.0178) | -0.0033(0.0167) |
| 49 | Hip circumference | 0.0085(0.0194) | -0.0045(0.0174) | -0.0208(0.0180) | 0.009(0.0168) |
| 50 | Standing height | -0.0015(0.0225) | -0.0031(0.0204) | 0.0164(0.0187) | 0.0164(0.0175) |
| 20015 | Sitting height | -0.0050(0.0213) | -0.0064(0.0192) | 0.0188(0.0187) | 0.0188(0.0174) |
| 21001 | Body mass index (BMI) | 0.0073(0.0197) | 0.0015(0.0177) | -0.0206(0.0181) | -0.0092(0.0169) |
| 21002 | Weight | 0.0044(0.0199) | -0.003(0.0178) | -0.0028(0.0182) | 0.005(0.017) |
| | *Physical measures* - *Blood pressure*: | | | | |
| 102 | Pulse rate, automated reading | -0.0035(0.0191) | 0.0073(0.0172) | 0.0220(0.0181) | -0.0219(0.017) |
| 4079 | Diastolic blood pressure, automated reading | -0.0004(0.0184) | -0.0066(0.0166) | 0.0002(0.0179) | 0.0222(0.0168) |
| 4080 | Systolic blood pressure, automated reading | -0.0070(0.0187) | -0.005(0.0169) | 0.0222(0.0180) | 0.0133(0.0169) |
| | *Physical measures* - *Bone-densitometry of heel*: | | | | |
| 78 | Heel bone mineral density (BMD) T-score, automated | 0.0011(0.0259) | -0.0118(0.0234) | -0.0011(0.0244) | 0.029(0.0229) |
| 3143 | Ankle spacing width | 0.0044(0.0240) | -0.0043(0.0219) | -0.0048(0.0233) | 0.0072(0.0221) |
| 3144 | Heel Broadband ultrasound attenuation, direct entry | 0.0062(0.0252) | -0.0073(0.0229) | -0.0294(0.0241) | 0.0212(0.0227) |
| 3147 | Heel quantitative ultrasound index (QUI), direct entry | 0.0011(0.0259) | -0.0118(0.0234) | -0.0011(0.0244) | 0.029(0.0229) |
| 3148 | Heel bone mineral density (BMD) | 0.0017(0.0257) | -0.0123(0.0232) | -0.0034(0.0243) | 0.0291(0.0228) |
| | *lifestyle and environment - Alcohol:* | | | | |
| 1558 | Alcohol intake frequency | 0.0028(0.0163) | 0.0086(0.0149) | -0.0065(0.0165) | -0.0223(0.0157) |
| 1598 | Average weekly spirits intake | -0.0044(0.0203) | -0.0038(0.0188) | 0.0175(0.0204) | 0.0173(0.0196) |
| | *Lifestyle and environment* - *Diet:* | | | | |
| 1309 | Fresh fruit intake | 0.0012(0.0169) | 0.0002(0.0154) | -0.0082(0.0168) | -0.0023(0.0159) |
| 1329 | Oily fish intake | -0.0044(0.0169) | 0.0042(0.0154) | 0.0180(0.0168) | -0.0151(0.0159) |
| 1369 | Beef intake | 0.0059(0.0165) | 0.0012(0.0151) | -0.0223(0.0166) | -0.0046(0.0158) |
| 1379 | Lamb/mutton intake | 0.0042(0.0156) | 0.0066(0.0145) | -0.0226(0.0161) | -0.0226(0.0155) |
| 1458 | Cereal intake | 0.0023(0.0182) | -0.0024(0.0164) | -0.0017(0.0174) | 0.0027(0.0164) |
| 1478 | Salt added to food | 0.0047(0.0174) | -0.0021(0.0158) | -0.0219(0.0171) | 0.0082(0.0162) |

| | | | | | |
|---|---|---|---|---|---|
| 1488 | Tea intake | -0.0012(0.0157) | 0.0052(0.0145) | 0.0012(0.0161) | -0.0198(0.0154) |
| 1518 | Hot drink temperature | -0.0076(0.0181) | -0.005(0.0163) | 0.0218(0.0175) | 0.0218(0.0164) |
| 1528 | Water intake | -0.0012(0.0181) | -0.0091(0.0163) | -0.0088(0.0175) | 0.0217(0.0164) |
| 1548 | Variation in diet | -0.0085(0.0164) | -0.0057(0.0151) | 0.0224(0.0167) | 0.0224(0.0159) |
| | *Lifestyle and environment - Electronic device use:* | | | | |
| 1120 | Weekly usage of mobile phone in last 3 months | 0.0039(0.0172) | -0.0043(0.0159) | -0.0242(0.0176) | 0.0189(0.0169) |
| 2237 | Plays computer games | 0.0039(0.0174) | -0.0023(0.0157) | -0.0219(0.0170) | 0.0112(0.0161) |
| | *Lifestyle and environment - Sexual factors:* | | | | |
| 2139 | Age first had sexual intercourse | 0.0039(0.0184) | 0.0002(0.0168) | -0.0236(0.0181) | -0.0003(0.0172) |
| 2149 | Lifetime number of sexual partners | -0.0100(0.0209) | -0.0066(0.0189) | 0.0241(0.0200) | 0.024(0.0187) |
| 2159 | Ever had same-sex intercourse | -0.0064(0.0165) | -0.0019(0.0155) | 0.0238(0.0171) | 0.0065(0.0165) |
| | *Lifestyle and environment - Sleep*: | | | | |
| 1160 | Sleep duration | -0.0058(0.0166) | -0.0066(0.0152) | 0.0223(0.0166) | 0.0222(0.0158) |
| 1170 | Getting up in morning | -0.0027(0.0174) | 0.0003(0.0157) | 0.0166(0.0171) | -0.0023(0.0161) |
| 1180 | Morning/evening person (chronotype) | -0.0043(0.0176) | 0.0005(0.0161) | 0.0234(0.0176) | -0.0003(0.0167) |
| 1190 | Nap during day | -0.0050(0.0171) | 0.0096(0.0156) | 0.0221(0.0170) | -0.0221(0.0161) |
| 1200 | Sleeplessness / insomnia | 0.0001(0.0176) | 0.0061(0.0159) | 0.0002(0.0172) | -0.0219(0.0162) |
| | *Lifestyle and environment - Smoking*: | | | | |
| 1239 | Current tobacco smoking | -0.0047(0.0153) | 0.0005(0.0143) | 0.0226(0.0159) | -0.0043(0.0154) |
| 1249 | Past tobacco smoking | 0.0092(0.0186) | 0.0068(0.0169) | -0.0232(0.0182) | -0.0221(0.0172) |
| 20116 | Smoking status | -0.0083(0.0185) | -0.0006(0.0166) | 0.0214(0.0176) | -0.0024(0.0165) |
| 20160 | Ever smoked | -0.0076(0.0186) | -0.0052(0.0168) | 0.0221(0.0178) | 0.0165(0.0168) |
| | *Lifestyle and environment - Sun exposure*: | | | | |
| 1717 | Skin colour | 0.009(0.0179) | -0.0007(0.0162) | -0.0219(0.0174) | 0.0041(0.0164) |
| | *Early life factors*: | | | | |
| 1687 | Comparative body size at age 10 | 0.0048(0.0184) | -0.006(0.0166) | -0.0219(0.0177) | 0.0219(0.0166) |
| 1697 | Comparative height size at age 10 | 0.0043(0.0203) | 0.0022(0.0182) | -0.0135(0.0184) | -0.0015(0.0171) |
| 20022 | Birth weight | 0.0047(0.0221) | 0.006(0.0206) | -0.0303(0.0223) | -0.0184(0.0214) |
| | *Female-specific factors*: | | | | |
| 2714 | Age when periods started (menarche) | -0.0073(0.0247) | -0.0079(0.0222) | 0.0291(0.0236) | 0.0097(0.0221) |
| 2744 | Birth weight of first child | -0.0023(0.0256) | -0.0076(0.0236) | 0.0059(0.0257) | 0.0349(0.0245) |
| | *Male-specific factors*: | | | | |
| 2375 | Relative age of first facial hair | 0.0042(0.026) | -0.0115(0.0242) | -0.0331(0.026) | 0.0345(0.0249) |
| 2385 | Relative age voice broke | -0.0022(0.0259) | 0.004(0.0244) | 0.0024(0.0263) | -0.0148(0.0254) |
| 2395 | Hair/balding pattern | -0.0032(0.0284) | 0.0027(0.0258) | 0.0014(0.0269) | 0.0016(0.0254) |

| | | | | | |
|---|---|---|---|---|---|
| | *Psychosocial factors - Mental health*: | | | | |
| 1920 | Mood swings | 0.0011(0.0171) | 0.001(0.0157) | -0.0083(0.0171) | -0.0057(0.0162) |
| 1940 | Irritability | -0.0065(0.0173) | 0.0034(0.0159) | 0.0227(0.0173) | -0.0115(0.0165) |
| 1950 | Sensitivity / hurt feelings | 0.0046(0.0159) | 0.0017(0.0148) | -0.0229(0.0164) | -0.0098(0.0158) |
| 1960 | Fed-up feelings | -0.0034(0.0167) | 0.0045(0.0153) | 0.0212(0.0169) | -0.0219(0.0161) |
| 1970 | Nervous feelings | -0.0038(0.0165) | 0.004(0.0152) | 0.0226(0.0167) | -0.0227(0.016) |
| 1980 | Worrier / anxious feelings | 0.0007(0.0178) | 0.0059(0.0161) | 0.0001(0.0174) | -0.0223(0.0164) |
| 1990 | Tense / 'highly strung' | -0.002(0.0168) | 0.0031(0.0155) | 0.0141(0.017) | -0.0153(0.0163) |
| 2000 | Worry too long after embarrassment | 0.0047(0.0187) | 0.0022(0.0169) | -0.0221(0.0179) | -0.0111(0.0168) |
| 20127 | Neuroticism score | -0.0008(0.0208) | 0.0048(0.019) | 0.0176(0.0199) | -0.0244(0.0187) |
| 2030 | Guilty feelings | 0.0074(0.0177) | 0.0011(0.0161) | -0.0223(0.0173) | -0.0051(0.0164) |
| 2040 | Risk taking | -0.0018(0.0176) | 0.0072(0.016) | 0.0006(0.0174) | -0.0226(0.0164) |
| 2050 | Frequency of depressed mood in last 2 weeks | 0.0012(0.0175) | -0.0076(0.016) | -0.0112(0.0173) | 0.0226(0.0164) |
| 2060 | Frequency of unenthusiasm / disinterest | -0.0001(0.0166) | 0.0009(0.0153) | -0.0004(0.0168) | -0.0052(0.0161) |
| 2070 | Frequency of tenseness / restlessness | 0.0046(0.0176) | 0.0073(0.0161) | -0.0227(0.0174) | -0.0226(0.0165) |
| 2080 | Frequency of tiredness / lethargy in last 2 weeks | -0.0051(0.0177) | 0.0074(0.0161) | 0.0223(0.0174) | -0.0223(0.0164) |
| 2090 | Seen doctor (GP) for nerves, anxiety, tension or depression | -0.0043(0.0163) | 0.0005(0.015) | 0.0111(0.0165) | 0.0001(0.0158) |
| | *Cognitive function - Reaction time*: | | | | |
| 403 | Number of times snap-button pressed | -0.0032(0.0158) | -0.0003(0.0146) | 0.0112(0.0164) | 0.0057(0.0157) |
| | *Health and medical history - Eyesight*: | | | | |
| 2217 | Age started wearing glasses or contact lenses | -0.0053(0.0186) | -0.006(0.017) | 0.0242(0.0184) | 0.0211(0.0175) |
| | *Sociodemographics - Education*: | | | | |
| 6138 | Qualifications | -0.0063(0.0189) | -0.0096(0.017) | 0.0213(0.0178) | 0.0213(0.0166) |
| | *Sociodemographics - Household*: | | | | |
| 728 | Number of vehicles in household | -0.0045(0.0155) | -0.0012(0.0145) | 0.0227(0.0162) | 0.0057(0.0156) |
| | *Verbal interview - Medical conditions*: | | | | |
| 20009 | Interpolated Age of participant when non-cancer illness first diagnosed | 0.0006(0.0194) | -0.0065(0.0179) | -0.0009(0.0197) | 0.025(0.0188) |

**Table S7**. **Genetic and residual covariances between 58 UKBB phenotypes and PC1/PC2 estimated by bivariate GREML model across POP2+POP3**. SE denotes standard error.

| UKBB data field | Phenotypes | Genetic covariance with PC1 (SE) | Genetic covariance with PC2 (SE) | Residual covariance with PC1 (SE) | Residual covariance with PC2 (SE) |
|---|---|---|---|---|---|
| | *Physical measures - Anthropometry*: | | | | |
| 48 | Waist circumference | 0.0003(0.0192) | 0.004(0.0171) | 0.0015(0.0177) | -0.0083(0.0163) |
| 49 | Hip circumference | 0.0038(0.0192) | 0.0048(0.0181) | -0.0019(0.0178) | -0.0079(0.0165) |
| 50 | Standing height | 0.0045(0.0218) | 0.0003(0.0197) | 0.0068(0.0181) | -0.0067(0.0169) |
| 20015 | Sitting height | 0.0048(0.0209) | 0.002(0.0187) | 0.0076(0.0181) | -0.0074(0.0168) |
| 21001 | Body mass index (BMI) | 0.0032(0.0195) | 0.0056(0.0183) | -0.0082(0.0178) | -0.0078(0.0165) |
| 21002 | Weight | 0.0016(0.0199) | 0.0045(0.0185) | 0.0082(0.018) | -0.0077(0.0166) |
| | *Physical measures - Blood pressure*: | | | | |
| 102 | Pulse rate, automated reading | -0.0043(0.0178) | 0.0045(0.016) | 0.0091(0.0173) | -0.0091(0.0162) |
| 4079 | Diastolic blood pressure, automated reading | 0.0009(0.0183) | -0.0005(0.0164) | -0.009(0.0175) | 0.0087(0.0164) |
| 4080 | Systolic blood pressure, automated reading | -0.0027(0.018) | -0.0015(0.0162) | 0.009(0.0174) | 0.0072(0.0163) |
| | *Physical measures - Bone-densitometry of heel*: | | | | |
| 78 | Heel bone mineral density (BMD) T-score, automated | 0.0107(0.0241) | -0.0096(0.023) | -0.0124(0.0234) | 0.0121(0.0224) |
| 3143 | Ankle spacing width | 0.0015(0.0257) | 0.0014(0.0231) | 0.0024(0.0243) | -0.0121(0.0227) |
| 3144 | Heel Broadband ultrasound attenuation, direct entry | 0.0092(0.0239) | -0.0086(0.0234) | -0.0124(0.0233) | 0.012(0.0225) |
| 3147 | Heel quantitative ultrasound index (QUI), direct entry | 0.0107(0.0241) | -0.0096(0.023) | -0.0124(0.0234) | 0.0121(0.0224) |
| 3148 | Heel bone mineral density (BMD) | 0.0032(0.0243) | -0.0098(0.0231) | -0.0123(0.0235) | 0.012(0.0224) |
| | *lifestyle and environment - Alcohol:* | | | | |
| 1558 | Alcohol intake frequency | 0.0024(0.0172) | 0.0037(0.0153) | -0.0089(0.0168) | -0.0088(0.0155) |
| 1598 | Average weekly spirits intake | 0.001(0.0194) | -0.004(0.0181) | -0.003(0.0198) | 0.0114(0.019) |
| 1628 | Alcohol intake versus 10 years previously | -0.0012(0.0164) | 0.0038(0.0151) | 0.0011(0.0168) | -0.0096(0.016) |
| | *Lifestyle and environment - Diet:* | | | | |
| 1309 | Fresh fruit intake | -0.0036(0.0158) | 0.0004(0.0145) | 0.0091(0.016) | -0.0022(0.0153) |
| 1319 | Dried fruit intake | -0.0031(0.0156) | 0.007(0.0187) | 0.0092(0.0161) | -0.0077(0.0162) |
| 1329 | Oily fish intake | 0.0015(0.0151) | 0.004(0.0141) | -0.0091(0.0157) | -0.0092(0.0151) |
| 1339 | Non-oily fish intake | 0.0022(0.015) | 0.0039(0.0139) | -0.0092(0.0157) | -0.0092(0.015) |
| 1349 | Processed meat intake | -0.001(0.0165) | 0.0032(0.0149) | 0.009(0.0164) | -0.009(0.0154) |

| | | | | | |
|---|---|---|---|---|---|
| 1478 | Salt added to food | 0.0048(0.0167) | -0.0038(0.0153) | -0.0089(0.0165) | 0.0089(0.0157) |
| 1518 | Hot drink temperature | -0.0027(0.0167) | 0.0013(0.0151) | 0.0033(0.0166) | -0.0002(0.0156) |
| 1528 | Water intake | -0.0034(0.0171) | -0.0037(0.0154) | 0.0028(0.0168) | 0.0089(0.0157) |
| 1548 | Variation in diet | -0.0027(0.0156) | 0.0038(0.0143) | 0.009(0.0159) | -0.0092(0.0152) |
| | *Lifestyle and environment - Electronic device use:* | | | | |
| 1110 | Length of mobile phone use | 0.002(0.0162) | 0.0032(0.0148) | -0.0091(0.0163) | -0.006(0.0155) |
| 1130 | Hands-free device/speakerphone use with mobile phone in last 3 month | 0.0043(0.0163) | -0.0059(0.0171) | -0.01(0.017) | 0.0095(0.0169) |
| 2237 | Plays computer games | 0.0037(0.0151) | 0.0031(0.014) | -0.0091(0.0157) | -0.0092(0.0151) |
| | *Lifestyle and environment - Sexual factors:* | | | | |
| 2139 | Age first had sexual intercourse | 0.0034(0.0169) | -0.0029(0.0156) | -0.0097(0.0171) | 0.0067(0.0164) |
| | *Lifestyle and environment - Sleep*: | | | | |
| 1160 | Sleep duration | -0.0077(0.0161) | -0.0063(0.0187) | 0.0089(0.016) | 0.0076(0.0162) |
| 1170 | Getting up in morning | -0.0043(0.0175) | 0.0069(0.0157) | 0.0088(0.0169) | -0.0088(0.0159) |
| 1180 | Morning/evening person (chronotype) | -0.0003(0.0184) | -0.0064(0.0186) | 0.0034(0.0177) | 0.0084(0.0169) |
| 1190 | Nap during day | -0.0037(0.0153) | 0.0045(0.0141) | 0.0092(0.0159) | -0.0091(0.0151) |
| 1200 | Sleeplessness / insomnia | 0.0016(0.0151) | -0.0026(0.014) | -0.0092(0.0157) | 0.0092(0.0151) |
| | *Lifestyle and environment - Smoking*: | | | | |
| 1249 | Past tobacco smoking | -0.0064(0.0175) | 0.0037(0.0158) | -0.0095(0.0174) | -0.0095(0.0164) |
| 20116 | Smoking status | -0.0044(0.0167) | -0.0005(0.015) | 0.009(0.0166) | -0.0008(0.0157) |
| | *Lifestyle and environment - Sun exposure*: | | | | |
| 1717 | Skin colour | 0.0003(0.0185) | -0.0012(0.0166) | -0.0086(0.0173) | 0.0086(0.0161) |
| | *Early life factors*: | | | | |
| 1687 | Comparative body size at age 10 | -0.0041(0.0188) | -0.0029(0.0169) | 0.0087(0.0176) | 0.0086(0.0164) |
| 1697 | Comparative height size at age 10 | -0.0011(0.02) | 0.0022(0.0179) | 0.0082(0.018) | -0.0082(0.0168) |
| | *Female-specific factors*: | | | | |
| 2714 | Age when periods started (menarche) | -0.0081(0.0227) | -0.0057(0.0206) | 0.0114(0.0218) | 0.0115(0.0206) |
| 2734 | Number of live births | 0.0013(0.0208) | 0.004(0.0189) | -0.0116(0.0208) | -0.0116(0.0197) |
| 2744 | Birth weight of first child | -0.0014(0.0244) | -0.0046(0.0226) | 0.0138(0.0248) | 0.0142(0.0236) |
| | *Male-specific factors*: | | | | |
| 2375 | Relative age of first facial hair | 0.0038(0.0263) | -0.0081(0.0243) | -0.0151(0.0267) | 0.015(0.0255) |
| 2395 | Hair/balding pattern | -0.0081(0.0293) | 0.0074(0.0264) | 0.0073(0.0277) | -0.0138(0.0259) |
| | *Psychosocial factors - Mental health*: | | | | |
| 1940 | Irritability | -0.0005(0.0154) | -0.0019(0.0145) | 0.0017(0.0161) | 0.006(0.0155) |
| 1960 | Fed-up feelings | 0.0017(0.0158) | -0.0034(0.0146) | -0.0093(0.0162) | 0.0093(0.0155) |

| | | | | | |
|---|---|---|---|---|---|
| 2000 | Worry too long after embarrassment | 0.0046(0.0045) | -0.0027(0.0143) | -0.0094(0.0049) | 0.0095(0.0154) |
| 20127 | Neuroticism score | 0.0028(0.0171) | 0.0032(0.0161) | -0.0106(0.0179) | -0.0106(0.0173) |
| 2070 | Frequency of tenseness / restlessness | 0.0015(0.0161) | 0.0037(0.0144) | -0.0094(0.0164) | -0.0094(0.0153) |
| 2080 | Frequency of tiredness / lethargy in last 2 weeks | -0.0009(0.0153) | 0.0045(0.0142) | 0.0028(0.0159) | -0.0094(0.0153) |
| | *Cognitive function - Reaction time*: | | | | |
| 404 | Duration to first press of snap-button in each round | 0.0064(0.0238) | 0.0094(0.0238) | -0.0144(0.0247) | -0.0139(0.0242) |
| | *Health and medical history*: | | | | |
| 2217 | Age started wearing glasses or contact lenses | 0.0061(0.02) | -0.0055(0.0179) | -0.0096(0.0189) | 0.0095(0.0177) |
| 2247 | Hearing difficulty/problems | 0.003(0.0162) | 0.0006(0.0148) | -0.0094(0.0164) | -0.0008(0.0156) |
| 2443 | Diabetes diagnosed by doctor | -0.0046(0.0168) | 0.002(0.0152) | 0.009(0.0166) | -0.0058(0.0157) |
| | *Sociodemographics - Education*: | | | | |
| 6138 | Qualifications | -0.0031(0.0183) | -0.0061(0.0164) | 0.0086(0.0172) | 0.0086(0.016) |
| | *Sociodemographics - Household*: | | | | |
| 699 | Length of time at current address | -0.0055(0.0206) | -0.0005(0.0143) | 0.0078(0.0173) | -0.0005(0.0152) |
| 738 | Average total household income before tax | -0.0015(0.0164) | -0.0033(0.0154) | 0.0099(0.017) | 0.0099(0.0164) |

**Table S8. Genetic and residual covariances between 70 UKBB phenotypes and PC1/PC2 estimated by bivariate GREML model across POP1+POP3.** SE denotes standard error.

| UKBB data field | Phenotypes | Genetic covariance with PC1 (SE) | Genetic covariance with PC2 (SE) | Residual covariance with PC1 (SE) | Residual covariance with PC2 (SE) |
|---|---|---|---|---|---|
| | *Physical measures - Anthropometry*: | | | | |
| 48 | Waist circumference | -0.0187(0.1147) | 0.0878(0.1339) | 0.0368(0.1025) | -0.0945(0.1111) |
| 49 | Hip circumference | -0.0234(0.0909) | 0.054(0.1061) | 0.0383(0.0802) | -0.0554(0.087) |
| 50 | Standing height | -0.0004(0.0682) | -0.07(0.0781) | 0.0132(0.0535) | 0.064(0.0579) |
| 20015 | Sitting height | 0.0048(0.0378) | -0.0333(0.0437) | -0.0003(0.0313) | 0.0335(0.034) |
| 21001 | Body mass index (BMI) | -0.0065(0.0467) | 0.0386(0.0545) | 0.0113(0.0412) | -0.0398(0.0446) |
| 21002 | Weight | -0.0201(0.141) | 0.0719(0.1643) | 0.0387(0.1229) | -0.0758(0.1334) |
| | *Physical measures - Blood pressure*: | | | | |
| 102 | Pulse rate, automated reading | -0.0656(0.1079) | -0.0625(0.1258) | 0.0803(0.0985) | 0.0668(0.1063) |
| 4079 | Diastolic blood pressure, automated reading | -0.0581(0.0982) | 0.0352(0.1146) | 0.065(0.09) | -0.0238(0.0971) |
| 4080 | Systolic blood pressure, automated reading | -0.0595(0.1685) | 0.014(0.1966) | 0.0614(0.1548) | 0.0069(0.1671) |
| | *Physical measures - Bone-densitometry of heel*: | | | | |
| 78 | Heel bone mineral density (BMD) T-score, automated | -0.007(0.0152) | 0.016(0.0176) | 0.0085(0.0136) | -0.0158(0.0147) |
| 3143 | Ankle spacing width | -0.0088(0.0483) | 0.0348(0.0559) | 0.0107(0.0441) | -0.0384(0.0473) |
| 3144 | Heel Broadband ultrasound attenuation, direct entry | -0.082(0.2359) | 0.2011(0.273) | 0.111(0.2111) | -0.1974(0.2271) |
| 3147 | Heel quantitative ultrasound index (QUI), direct entry | -0.1238(0.2696) | 0.2827(0.312) | 0.1502(0.2415) | -0.2802(0.2597) |
| 3148 | Heel bone mineral density (BMD) | -0.0008(0.0017) | 0.0019(0.002) | 0.001(0.0015) | -0.0018(0.0016) |
| | *lifestyle and environment - Alcohol:* | | | | |
| 1558 | Alcohol intake frequency | 0.0066(0.0132) | 0.0102(0.0154) | -0.0079(0.0124) | -0.0109(0.0133) |
| | *Lifestyle and environment - Diet:* | | | | |
| 1309 | Fresh fruit intake | -0.0034(0.0135) | -0.0155(0.0157) | 0.0039(0.013) | 0.0181(0.0139) |
| 1319 | Dried fruit intake | 0.0165(0.0162) | -0.0263(0.0189) | -0.0214(0.0157) | 0.0299(0.0168) |
| 1329 | Oily fish intake | 0.0055(0.0077) | -0.0001(0.009) | -0.0074(0.0074) | 0.0001(0.0079) |
| 1339 | Non-oily fish intake | 0.0009(0.0066) | -0.002(0.0077) | -0.0012(0.0064) | 0.0024(0.0068) |
| 1349 | Processed meat intake | -0.0004(0.0088) | 0.0024(0.0103) | 0.0007(0.0084) | -0.0027(0.009) |
| 1359 | Poultry intake | -0.0044(0.0077) | 0.0028(0.0089) | 0.0062(0.0074) | -0.0033(0.0079) |
| 1369 | Beef intake | -0.0046(0.007) | 0.0045(0.0082) | 0.0064(0.0068) | -0.0051(0.0073) |
| 1379 | Lamb/mutton intake | -0.0014(0.006) | -0.0026(0.0069) | 0.0019(0.0057) | 0.0031(0.0061) |
| 1408 | Cheese intake | 0.0011(0.0093) | -0.0122(0.0108) | -0.0014(0.009) | 0.0138(0.0096) |

| | | | | | |
|---|---|---|---|---|---|
| 1458 | Cereal intake | -0.0079(0.0242) | 0.0034(0.0282) | 0.0102(0.023) | -0.0048(0.0247) |
| 1478 | Salt added to food | -0.0017(0.0076) | -0.0004(0.0088) | 0.0023(0.0071) | 0.0008(0.0077) |
| 1518 | Hot drink temperature | -0.0011(0.0051) | 0.0004(0.0059) | 0.0011(0.0048) | -0.0005(0.0051) |
| 1528 | Water intake | 0.0036(0.0205) | -0.0192(0.0239) | -0.0047(0.0196) | 0.0216(0.021) |
| | *Lifestyle and environment - Electronic device use:* | | | | |
| 1110 | Length of mobile phone use | 0.0008(0.0114) | 0.002(0.0133) | -0.0011(0.0108) | -0.0023(0.0116) |
| 2237 | Plays computer games | -0.0001(0.0044) | 0.0014(0.0051) | 0.0003(0.0042) | -0.0016(0.0045) |
| | *Lifestyle and environment - Sexual factors:* | | | | |
| 2139 | Age first had sexual intercourse | -0.021(0.0366) | 0.0219(0.0429) | 0.0241(0.0335) | -0.023(0.0364) |
| | *Lifestyle and environment - Sleep*: | | | | |
| 1160 | Sleep duration | -0.0027(0.0093) | 0.0112(0.0109) | 0.0037(0.0088) | -0.0124(0.0095) |
| 1170 | Getting up in morning | 0.002(0.007) | 0.0013(0.0081) | -0.0025(0.0064) | -0.001(0.0069) |
| 1180 | Morning/evening person (chronotype) | 0.0029(0.0093) | -0.0005(0.0109) | -0.0029(0.0085) | 0.0004(0.0092) |
| 1190 | Nap during day | 0.0014(0.0052) | -0.001(0.006) | -0.0019(0.0048) | 0.001(0.0052) |
| 1200 | Sleeplessness / insomnia | 0.0009(0.0063) | 0.0014(0.0074) | -0.001(0.0059) | -0.0019(0.0064) |
| 1210 | Snoring | 0.0037(0.0042) | 0.0003(0.0049) | -0.0049(0.004) | -0.0003(0.0043) |
| | *Lifestyle and environment - Smoking*: | | | | |
| 1239 | Current tobacco smoking | -0.0017(0.0036) | 0.0002(0.0041) | 0.0023(0.0034) | -0.0003(0.0037) |
| 1249 | Past tobacco smoking | -0.003(0.0122) | -0.0008(0.0143) | 0.0039(0.0116) | 0.0008(0.0125) |
| 20116 | Smoking status | -0.0003(0.0063) | -0.0008(0.0073) | 0.0006(0.0058) | 0.0008(0.0063) |
| 20160 | Ever smoked | 0.0019(0.0043) | -0.0008(0.0051) | -0.0024(0.0041) | 0.0004(0.0044) |
| | *Lifestyle and environment - Sun exposure*: | | | | |
| 1717 | Skin colour | 0.0002(0.0055) | 0.0016(0.0064) | -0.0015(0.0049) | -0.0012(0.0053) |
| | *Early life factors*: | | | | |
| 1687 | Comparative body size at age 10 | 0.0016(0.0031) | 0.0007(0.0036) | -0.002(0.0028) | -0.0007(0.0031) |
| 1697 | Comparative height size at age 10 | -0.0007(0.0034) | -0.0006(0.0039) | 0.0007(0.0029) | 0.0004(0.0031) |
| 20022 | Birth weight | 0.002(0.0075) | 0.0011(0.0087) | -0.0021(0.0071) | -0.0012(0.0076) |
| | *Female-specific factors*: | | | | |
| 2714 | Age when periods started (menarche) | -0.0126(0.0185) | -0.0272(0.0218) | 0.0176(0.0175) | 0.0305(0.0189) |
| 2734 | Number of live births | 0.0012(0.0128) | -0.0093(0.0151) | -0.0023(0.0125) | 0.0103(0.0134) |
| | *Male-specific factors*: | | | | |
| 2375 | Relative age of first facial hair | 0.0034(0.0065) | -0.0021(0.0074) | -0.0044(0.0059) | 0.002(0.0063) |
| 2395 | Hair/balding pattern | -0.006(0.0164) | 0.0022(0.0188) | 0.007(0.0143) | -0.0019(0.0154) |
| | *Psychosocial factors - Mental health*: | | | | |
| 1920 | Mood swings | 0.0012(0.0044) | -0.0003(0.0052) | -0.0014(0.0041) | 0.0002(0.0044) |

| | | | | | |
|---|---|---|---|---|---|
| 1930 | Miserableness | 0.0018(0.0042) | 0.0058(0.0049) | -0.0024(0.004) | -0.0066(0.0043) |
| 1940 | Irritability | 0.0004(0.0041) | -0.0005(0.0047) | -0.0002(0.0038) | 0.0006(0.0041) |
| 1950 | Sensitivity / hurt feelings | 0.0004(0.0042) | 0.0029(0.0049) | -0.0006(0.004) | -0.0035(0.0043) |
| 1960 | Fed-up feelings | -0.0002(0.0043) | 0.0013(0.005) | 0.0005(0.004) | -0.0017(0.0043) |
| 1970 | Nervous feelings | 0.0011(0.0036) | -0.0001(0.0042) | -0.0015(0.0034) | 0(0.0037) |
| 1980 | Worrier / anxious feelings | 0.001(0.0044) | 0.0007(0.0051) | -0.0014(0.0041) | -0.0009(0.0044) |
| 1990 | Tense / 'highly strung' | -0.0003(0.0035) | 0.0008(0.0041) | 0.0004(0.0032) | -0.001(0.0035) |
| 2000 | Worry too long after embarrassment | 0.0016(0.0044) | -0.0025(0.0051) | -0.0021(0.0041) | 0.0028(0.0044) |
| 20127 | Neuroticism score | 0.0078(0.0315) | -0.0095(0.0371) | -0.0092(0.0295) | 0.0082(0.032) |
| 2030 | Guilty feelings | 0.0016(0.0039) | -0.0015(0.0046) | -0.002(0.0037) | 0.0016(0.004) |
| 2040 | Risk taking | -0.0007(0.004) | -0.0001(0.0046) | 0.001(0.0038) | 0.0001(0.0041) |
| 2080 | Frequency of tiredness / lethargy in last 2 weeks | -0.0055(0.0073) | 0.0008(0.0086) | 0.007(0.0069) | -0.0011(0.0075) |
| 2090 | Seen doctor (GP) for nerves, anxiety, tension or depression | 0.001(0.0039) | 0.0032(0.0046) | -0.0013(0.0038) | -0.0037(0.0041) |
| | *Cognitive function - Reaction time*: | | | | |
| 20023 | Mean time to correctly identify matches | -0.159(0.91) | 0.5117(1.0597) | 0.226(0.863) | -0.6117(0.9266) |
| | *Health and medical history*: | | | | |
| 2217 | Age started wearing glasses or contact lenses | 0.015(0.1613) | -0.1278(0.1888) | -0.0136(0.1481) | 0.1421(0.1604) |
| 2443 | Diabetes diagnosed by doctor | 0.0001(0.0018) | -0.0004(0.0021) | -0.0001(0.0017) | 0.0004(0.0018) |
| 2463 | Fractured/broken bones in last 5 years | 0.0008(0.0026) | -0.0045(0.003) | -0.0012(0.0025) | 0.0052(0.0026) |
| | *Sociodemographics - Education*: | | | | |
| 6138 | Qualifications | 0.004(0.0093) | -0.0085(0.0108) | -0.0062(0.0083) | 0.0096(0.009) |
| | *Sociodemographics - Household*: | | | | |
| 738 | Average total household income before tax | -0.0039(0.0101) | -0.0088(0.0117) | 0.0048(0.0096) | 0.0101(0.0103) |
| | *Verbal interview - Medical conditions*: | | | | |
| 135 | Number of self-reported non-cancer illnesses | 0.0001(0.0156) | -0.0047(0.0182) | 0.0015(0.0147) | 0.0049(0.0158) |

**Table S9. Genetic variance, interaction variance and their covariance component estimates for 70 UKBB phenotypes with significant heritabilities across POP1+POP2 with the covariate PC1**. The estimates which were not within the valid parameter space were excluded for follow-up analyses as shown in the column "Note". SE denotes standard error. DF denotes degree of freedom.

| UKBB data field | $\mathrm{var}(\mathbf{g}_0)$ (SE) | $\mathrm{var}(\mathbf{g}_1)$ (SE) | $\mathrm{cov}(\mathbf{g}_0, \mathbf{g}_1)$ (SE) | $\mathrm{var}(\mathbf{e}_0)$ (SE) | Note | *P*-value by LRT comparing with baseline model (DF =2) |
|---|---|---|---|---|---|---|
| 48 | 22.8234(3.4160) | 1.0785(0.8583) | -2.8866(1.0905) | 115.2992(3.6088) | | 0.0220 |
| 49 | 16.1120(2.1416) | 0.3507(0.5114) | -1.2321(0.6705) | 69.6114(2.2349) | | 0.1322 |
| 50 | 19.3852(1.0527) | -0.0938(0.2146) | 0.2898(0.3109) | 19.9075(0.9541) | Excluded | |
| 20015 | 4.5722(0.3539) | -0.1247(0.0649) | 0.1097(0.1028) | 9.1564(0.3431) | Excluded | |
| 21001 | 4.6117(0.5676) | 0.2398(0.1522) | -0.3711(0.1822) | 17.8795(0.5895) | | 0.1285 |
| 21002 | 43.0297(4.9553) | 2.8555(1.3309) | -5.5148(1.5958) | 151.7953(5.1194) | | 0.0029 |
| 102 | 19.6191(3.1902) | 1.5100(0.8223) | -3.8713(0.9883) | 103.7211(3.3880) | | 0.0002 |
| 4079 | 13.3974(2.7753) | -0.4101(0.6394) | 0.6785(0.8590) | 95.8879(2.9699) | Excluded | |
| 4080 | 43.8004(8.2129) | 1.7493(2.1498) | -4.9706(2.5723) | 274.5824(8.8145) | | 0.1119 |
| 78 | 0.4026(0.0676) | 0.0263(0.0189) | 0.0416(0.0185) | 1.0632(0.0679) | | 9.39E-07 |
| 3143 | 2.7502(0.6788) | -0.0278(0.1648) | 0.4103(0.1829) | 12.7605(0.7007) | Excluded | |
| 3144 | 85.2801(16.0444) | 6.6743(4.6119) | 10.4970(4.4641) | 267.7478(16.2238) | | 2.87E-07 |
| 3147 | 126.3220(21.2130) | 8.2347(5.9147) | 13.0431(5.8075) | 333.5802(21.2942) | | 9.40E-07 |
| 3148 | 0.0048(0.0008) | 0.0003(0.0002) | 0.0006(0.0002) | 0.0135(0.0008) | | 1.58E-07 |
| 1558 | 0.1547(0.0528) | -0.0036(0.0125) | 0.0264(0.0168) | 2.1348(0.0587) | Excluded | |
| 1598 | 1.9868(0.7932) | 6.5790(0.4870) | -4.4794(0.4369) | 31.0538(1.0070) | Excluded | |
| 1309 | 0.0183(0.0348) | -0.0224(0.0072) | 0.0472(0.0112) | 2.0163(0.0402) | Excluded | |
| 1329 | 0.0669(0.0187) | 0.0049(0.0049) | -0.0090(0.0061) | 0.7365(0.0208) | | 0.3285 |
| 1369 | 0.0500(0.0167) | -0.0032(0.0043) | 0.0220(0.0055) | 0.6616(0.0184) | Excluded | |
| 1379 | 0.0275(0.0117) | -0.0027(0.0034) | 0.0212(0.0040) | 0.4843(0.0130) | Excluded | |
| 1458 | 0.9810(0.1847) | -0.0285(0.0422) | 0.0126(0.0576) | 6.7208(0.1981) | Excluded | |
| 1478 | 0.0819(0.0195) | -0.0030(0.0049) | 0.0313(0.0063) | 0.7404(0.0212) | Excluded | |
| 1488 | 0.7017(0.0510) | 0.6597(0.0380) | -0.7585(0.0436) | 6.4842(0.1076) | Excluded | |
| 1518 | 0.0435(0.0084) | -0.0033(0.0016) | 0.0113(0.0024) | 0.3076(0.0089) | Excluded | |
| 1528 | 0.1598(0.0832) | -0.0406(0.0174) | 0.1469(0.0256) | 4.2004(0.0949) | Excluded | |
| 1548 | 0.0248(0.0086) | -0.0017(0.0020) | 0.0038(0.0027) | 0.3413(0.0095) | Excluded | |
| 1120 | 0.1099(0.0425) | 0.0136(0.0120) | 0.0077(0.0134) | 1.4478(0.0469) | | 0.0799 |
| 2237 | 0.0249(0.0061) | 0.0039(0.0020) | -0.0034(0.0021) | 0.2304(0.0068) | | 0.1408 |
| 2139 | 1.5527(0.4033) | 0.3563(0.1283) | -0.1990(0.1348) | 13.1435(0.4406) | | 0.0047 |

| | | | | | | |
|---|---|---|---|---|---|---|
| 2149 | -36.1697(19.0655) | -5.4504(2.0441) | 36.1221(5.8221) | 1143.5765(23.37) | Excluded | |
| 2159 | 0.0000(0.0006) | -0.0003(0.0001) | 0.0010(0.0002) | 0.0297(0.0006) | Excluded | |
| 1160 | 0.0834(0.0267) | 0.0218(0.0091) | -0.0063(0.0093) | 1.0469(0.0299) | | 0.0073 |
| 1170 | 0.0601(0.0146) | -0.0041(0.0037) | 0.0183(0.0048) | 0.5567(0.0159) | Excluded | |
| 1180 | 0.0822(0.0244) | -0.0101(0.0053) | 0.0252(0.0075) | 0.8692(0.0265) | Excluded | |
| 1190 | 0.0291(0.0080) | 0.0072(0.0024) | -0.0096(0.0026) | 0.3010(0.0089) | | 0.0006 |
| 1200 | 0.0539(0.0124) | -0.0027(0.0030) | 0.0052(0.0039) | 0.4670(0.0134) | Excluded | |
| 1239 | 0.0004(0.0023) | -0.0012(0.0005) | 0.0036(0.0008) | 0.1299(0.0027) | Excluded | |
| 1249 | 0.2571(0.0390) | 0.0632(0.0133) | -0.1324(0.0146) | 1.4763(0.0418) | Excluded | |
| 20116 | 0.0700(0.0121) | 0.0057(0.0035) | -0.0032(0.0040) | 0.4228(0.0130) | | 0.1870 |
| 20160 | 0.0303(0.0058) | 0.0047(0.0016) | -0.0082(0.0019) | 0.1972(0.0063) | | 0.0001 |
| 1717 | 0.0263(0.0075) | -0.0023(0.0017) | 0.0162(0.0024) | 0.3254(0.0083) | Excluded | |
| 1687 | 0.0146(0.0029) | -0.0005(0.0008) | 0.0026(0.0009) | 0.1034(0.0031) | Excluded | |
| 1697 | 0.0270(0.0028) | -0.0001(0.0007) | -0.0001(0.0009) | 0.0839(0.0029) | Excluded | |
| 20022 | 0.0490(0.0186) | 0.0026(0.0035) | -0.0020(0.0047) | 0.3900(0.0194) | | 0.6955 |
| 2714 | 0.6343(0.1138) | 0.0031(0.0209) | -0.0263(0.0285) | 1.9979(0.1153) | | 0.5074 |
| 2744 | 0.2355(0.0859) | -0.0113(0.0150) | 0.0631(0.0206) | 1.3442(0.0887) | Excluded | |
| 2375 | 0.0399(0.0112) | -0.0033(0.0020) | 0.0117(0.0028) | 0.1671(0.0114) | Excluded | |
| 2385 | 0.0139(0.0058) | -0.0027(0.0008) | 0.0079(0.0014) | 0.0930(0.0060) | Excluded | |
| 2395 | 0.4116(0.0668) | 0.0069(0.0120) | -0.0088(0.0153) | 0.7675(0.0659) | | 0.8129 |
| 1920 | 0.0191(0.0058) | 0.0015(0.0015) | -0.0021(0.0018) | 0.2195(0.0064) | | 0.4826 |
| 1940 | 0.0177(0.0052) | -0.0023(0.0011) | 0.0056(0.0016) | 0.1925(0.0056) | Excluded | |
| 1950 | 0.0125(0.0057) | -0.0007(0.0013) | 0.0012(0.0018) | 0.2248(0.0063) | Excluded | |
| 1960 | 0.0160(0.0056) | 0.0012(0.0015) | -0.0011(0.0018) | 0.2152(0.0062) | | 0.7256 |
| 1970 | 0.0127(0.0044) | -0.0017(0.0010) | 0.0050(0.0014) | 0.1741(0.0049) | Excluded | |
| 1980 | 0.0247(0.0057) | 0.0001(0.0014) | -0.0013(0.0018) | 0.2115(0.0062) | | 0.6595 |
| 1990 | 0.0052(0.0031) | -0.0011(0.0007) | 0.0055(0.0010) | 0.1351(0.0035) | Excluded | |
| 2000 | 0.0326(0.0062) | -0.0003(0.0014) | 0.0001(0.0019) | 0.2118(0.0066) | Excluded | |
| 20127 | 1.7563(0.3177) | 0.0084(0.0695) | 0.0255(0.0929) | 8.5585(0.3316) | | 0.8753 |
| 2030 | 0.0204(0.0050) | -0.0011(0.0012) | 0.0026(0.0016) | 0.1843(0.0054) | Excluded | |
| 2040 | 0.0124(0.0043) | -0.0019(0.0009) | 0.0072(0.0012) | 0.1779(0.0047) | Excluded | |
| 2050 | 0.0123(0.0072) | -0.0025(0.0017) | 0.0136(0.0023) | 0.3273(0.0081) | Excluded | |
| 2060 | 0.0113(0.0077) | -0.0022(0.0017) | 0.0157(0.0024) | 0.3693(0.0087) | Excluded | |
| 2070 | 0.0106(0.0071) | -0.0031(0.0017) | 0.0116(0.0022) | 0.3240(0.0080) | Excluded | |
| 2080 | 0.0707(0.0174) | 0.0007(0.0048) | 0.0157(0.0058) | 0.6382(0.0190) | Excluded | |
| 2090 | 0.0135(0.0050) | 0.0006(0.0012) | -0.0027(0.0016) | 0.2007(0.0056) | | 0.1977 |
| 403 | 0.2533(0.0046) | -0.0006(0.0040) | -0.0029(0.0009) | 0.2533(0.0046) | Excluded | |

| | | | | | | |
|---|---|---|---|---|---|---|
| 2217 | 27.0236(7.4244) | -2.3572(1.1177) | -0.3735(2.0760) | 242.8999(7.9144) | Excluded | |
| 6138 | 0.1583(0.0246) | 0.0768(0.0101) | -0.0688(0.0086) | 0.7846(0.0265) | | 1.96E-18 |
| 728 | 0.0338(0.0165) | 0.0009(0.0045) | -0.0014(0.0054) | 0.6696(0.0185) | | 0.9670 |
| 20009 | 19.0474(6.7765) | 1.0624(1.3542) | -3.2719(1.8475) | 181.4122(7.2165) | | 0.1779 |

**Table S10**. **Genetic variance, interaction variance and their covariance component estimates for 70 UKBB phenotypes with significant heritabilities across POP1+POP2 with the covariate PC2.** The estimates which were not within the valid parameter space were excluded for follow-up analyses as shown in the column "Note". SE denotes standard error. DF denotes degree of freedom.

| UKBB data field | $\text{var}(\mathbf{g}_0)$ (SE) | $\text{var}(\mathbf{g}_1)$ (SE) | $\text{cov}(\mathbf{g}_0, \mathbf{g}_1)$ (SE) | $\text{var}(\mathbf{e}_0)$ (SE) | Note | *P*-value by LRT comparing with baseline model (DF =2) |
|---|---|---|---|---|---|---|
| 48 | 22.6677(3.4011) | 0.6079(0.4699) | 3.5679(1.0350) | 115.9131(3.5460) | | 0.0008 |
| 49 | 16.0363(2.1375) | 0.4629(0.3077) | 2.2087(0.6575) | 69.5760(2.2019) | | 0.0020 |
| 50 | 19.4046(1.0518) | -0.0410(0.1465) | -0.4103(0.3192) | 19.8373(0.9442) | Excluded | |
| 20015 | 4.5379(0.3536) | 0.0759(0.0575) | 0.0999(0.1115) | 8.9874(0.3409) | | 0.3857 |
| 21001 | 4.6304(0.5666) | 0.1250(0.0825) | 0.5573(0.1746) | 17.9787(0.5791) | | 0.0040 |
| 21002 | 42.9272(4.9474) | 1.3950(0.7474) | 5.2355(1.5208) | 153.3097(5.0251) | | 0.0020 |
| 102 | 19.4977(3.1888) | 0.6285(0.4871) | 2.5351(0.9932) | 104.6823(3.3361) | | 0.0308 |
| 4079 | 13.2579(2.7734) | 0.1359(0.3432) | 0.7811(0.8411) | 95.4995(2.9324) | | 0.6326 |
| 4080 | 43.2231(8.1925) | 1.4060(1.1705) | 5.3977(2.4994) | 275.4511(8.6274) | | 0.0920 |
| 78 | 0.4017(0.0681) | -0.0072(0.0061) | -0.0179(0.0158) | 1.0985(0.0675) | Excluded | |
| 3143 | 2.7566(0.6796) | -0.1655(0.0539) | -0.3505(0.1637) | 12.9057(0.6917) | Excluded | |
| 3144 | 84.7201(16.1744) | -1.1356(1.5932) | -4.0006(3.8517) | 276.3461(16.1749) | Excluded | |
| 3147 | 126.0446(21.3600) | -2.2620(1.9058) | -5.6293(4.9698) | 344.6565(21.1687) | Excluded | |
| 3148 | 0.0048(0.0008) | -0.0001(0.0001) | -0.0003(0.0002) | 0.0139(0.0008) | Excluded | |
| 1558 | 0.1525(0.0528) | 0.0148(0.0096) | 0.0161(0.0171) | 2.1192(0.0581) | | 0.1938 |
| 1598 | 0.7563(0.9430) | 16.6010(1.0157) | 4.1934(0.4631) | 25.6094(1.1035) | Excluded | |
| 1309 | 0.0108(0.0329) | -0.0148(0.0035) | -0.0529(0.0098) | 1.9700(0.0379) | Excluded | |
| 1329 | 0.0669(0.0187) | -0.0010(0.0027) | 0.0028(0.0061) | 0.7423(0.0204) | Excluded | |
| 1369 | 0.0521(0.0167) | -0.0030(0.0027) | -0.0194(0.0055) | 0.6595(0.0182) | Excluded | |
| 1379 | 0.0273(0.0117) | 0.0053(0.0027) | -0.0060(0.0041) | 0.4768(0.0130) | | 2.73E-05 |
| 1458 | 0.9782(0.1847) | 0.0012(0.0250) | 0.0310(0.0576) | 6.6938(0.1961) | | 0.7890 |
| 1478 | 0.0796(0.0196) | 0.0007(0.0036) | -0.0145(0.0066) | 0.7393(0.0211) | Excluded | |
| 1488 | 0.4535(0.1724) | 0.0406(0.0290) | 0.1418(0.0556) | 7.3141(0.1924) | Excluded | |
| 1518 | 0.0423(0.0084) | -0.0013(0.0012) | -0.0090(0.0026) | 0.3069(0.0089) | Excluded | |
| 1528 | 0.7063(0.1312) | -0.0328(0.0212) | -0.3455(0.0422) | 4.9554(0.1371) | Excluded | |
| 1548 | 0.0247(0.0086) | 0.0005(0.0014) | -0.0020(0.0028) | 0.3392(0.0094) | | 0.4123 |
| 1120 | 0.1081(0.0425) | 0.0093(0.0072) | 0.0078(0.0135) | 1.4543(0.0461) | | 0.2894 |
| 2237 | 0.0256(0.0061) | 0.0004(0.0009) | 0.0035(0.0019) | 0.2333(0.0066) | Excluded | |
| 2139 | 1.4802(0.4017) | 0.3641(0.0929) | 0.4340(0.1365) | 13.2176(0.4311) | | 0.0001 |

| | | | | | | |
|---|---|---|---|---|---|---|
| 2149 | -6.7960(19.5429) | -8.5329(2.3998) | -38.0657(5.4242) | 1128.2480(23.51) | Excluded | |
| 2159 | 0.0012(0.0006) | -0.0003(0.0001) | -0.0016(0.0002) | 0.0381(0.0007) | Excluded | |
| 1160 | 0.0828(0.0267) | 0.0235(0.0067) | 0.0151(0.0097) | 1.0463(0.0293) | | 0.0001 |
| 1170 | 0.0616(0.0145) | -0.0048(0.0022) | -0.0250(0.0048) | 0.5562(0.0156) | Excluded | |
| 1180 | 0.0827(0.0243) | -0.0056(0.0030) | -0.0265(0.0073) | 0.8643(0.0260) | Excluded | |
| 1190 | 0.0285(0.0081) | 0.0023(0.0014) | 0.0064(0.0026) | 0.3064(0.0088) | | 0.0595 |
| 1200 | 0.0534(0.0124) | 0.0004(0.0018) | -0.0001(0.0040) | 0.4644(0.0132) | | 0.9543 |
| 1239 | 0.0146(0.0041) | 0.0014(0.0010) | -0.0135(0.0014) | 0.1765(0.0043) | Excluded | |
| 1249 | 0.2777(0.0417) | 0.0272(0.0075) | 0.0992(0.0133) | 1.4904(0.0443) | Excluded | |
| 20116 | 0.0699(0.0121) | 0.0054(0.0024) | 0.0056(0.0040) | 0.4232(0.0128) | | 0.0233 |
| 20160 | 0.0308(0.0058) | 0.0017(0.0009) | 0.0056(0.0018) | 0.1996(0.0062) | | 0.0072 |
| 1717 | 0.0460(0.0094) | 0.0010(0.0019) | -0.0100(0.0032) | 0.3453(0.0101) | Excluded | |
| 1687 | 0.0147(0.0029) | -0.0005(0.0004) | -0.0018(0.0009) | 0.1033(0.0031) | Excluded | |
| 1697 | 0.0270(0.0028) | -0.0001(0.0004) | -0.0004(0.0009) | 0.0840(0.0028) | Excluded | |
| 20022 | 0.0480(0.0184) | -0.0012(0.0012) | 0.0071(0.0041) | 0.3949(0.0192) | Excluded | |
| 2714 | 0.6329(0.1137) | 0.0120(0.0155) | 0.0264(0.0297) | 1.9899(0.1145) | | 0.6695 |
| 2744 | 0.2488(0.0864) | 0.0059(0.0127) | -0.0211(0.0211) | 1.3147(0.0884) | | 0.1341 |
| 2375 | 0.0385(0.0112) | -0.0018(0.0012) | -0.0096(0.0027) | 0.1670(0.0113) | Excluded | |
| 2385 | 0.0185(0.0063) | -0.0019(0.0004) | -0.0075(0.0013) | 0.0921(0.0064) | Excluded | |
| 2395 | 0.4137(0.0668) | -0.0003(0.0064) | 0.0056(0.0152) | 0.7728(0.0653) | Excluded | |
| 1920 | 0.0193(0.0058) | 0.0010(0.0009) | 0.0023(0.0018) | 0.2198(0.0063) | | 0.3940 |
| 1940 | 0.0174(0.0052) | -0.0012(0.0006) | -0.0037(0.0016) | 0.1918(0.0056) | Excluded | |
| 1950 | 0.0124(0.0057) | 0.0001(0.0009) | -0.0004(0.0018) | 0.2241(0.0062) | | 0.8834 |
| 1960 | 0.0162(0.0056) | 0.0007(0.0009) | 0.0022(0.0018) | 0.2155(0.0061) | | 0.4926 |
| 1970 | 0.0123(0.0044) | 0.0002(0.0006) | -0.0006(0.0014) | 0.1727(0.0048) | | 0.6717 |
| 1980 | 0.0247(0.0057) | 0.0005(0.0009) | 0.0013(0.0019) | 0.2111(0.0062) | | 0.7755 |
| 1990 | 0.0122(0.0039) | -0.0011(0.0006) | -0.0076(0.0013) | 0.1478(0.0042) | Excluded | |
| 2000 | 0.0326(0.0062) | 0.0003(0.0009) | 0.0008(0.0019) | 0.2112(0.0065) | | 0.9192 |
| 20127 | 1.7552(0.3175) | 0.0395(0.0438) | 0.0757(0.0920) | 8.5297(0.3276) | | 0.5963 |
| 2030 | 0.0205(0.0050) | -0.0008(0.0006) | -0.0015(0.0016) | 0.1839(0.0053) | Excluded | |
| 2040 | 0.0141(0.0046) | -0.0018(0.0006) | -0.0084(0.0015) | 0.1840(0.0050) | Excluded | |
| 2050 | 0.0415(0.0097) | 0.0020(0.0020) | -0.0285(0.0032) | 0.3642(0.0101) | Excluded | |
| 2060 | 0.0406(0.0108) | 0.0051(0.0028) | -0.0282(0.0038) | 0.4119(0.0115) | Excluded | |
| 2070 | 0.0403(0.0103) | -0.0032(0.0015) | -0.0191(0.0031) | 0.3799(0.0109) | Excluded | |
| 2080 | 0.0709(0.0174) | 0.0010(0.0029) | -0.0087(0.0058) | 0.6380(0.0187) | Excluded | |
| 2090 | 0.0134(0.0050) | 0.0011(0.0008) | 0.0033(0.0016) | 0.2003(0.0055) | | 0.1522 |
| 403 | 0.0011(0.0039) | -0.0022(0.0004) | -0.0060(0.0012) | 0.2660(0.0046) | Excluded | |

| | | | | | | |
|---|---|---|---|---|---|---|
| 2217 | 26.5141(7.4474) | -1.0862(0.8221) | 0.0062(2.1969) | 242.1076(7.9367) | Excluded | |
| 6138 | 0.1752(0.0237) | 0.0459(0.0063) | 0.0862(0.0084) | 0.7995(0.0250) | | 3.22E-23 |
| 728 | 0.0337(0.0165) | 0.0055(0.0031) | 0.0036(0.0055) | 0.6650(0.0182) | | 0.0897 |
| 20009 | 18.6624(6.7718) | 0.9099(1.0368) | 2.4535(1.9616) | 181.8959(7.1638) | | 0.4897 |

**Table S11**. **Genetic variance, two interaction variances and their covariance component estimates for ten UKBB phenotypes across POP1+POP2 with simultaneous covariates PC1 and PC2**. The estimates which were not within the valid parameter space were excluded for follow-up analyses as shown in the column "Note". SE denotes standard error. DF denotes degree of freedom.

| UKBB data field | $\text{var}(\mathbf{g}_0)$ (SE) | $\text{var}(\mathbf{g}_1)$ (SE) | $\text{cov}(\mathbf{g}_0, \mathbf{g}_1)$ (SE) | $\text{var}(\mathbf{g}_2)$ (SE) | $\text{cov}(\mathbf{g}_0, \mathbf{g}_2)$ (SE) | $\text{var}(\mathbf{e}_0)$ (SE) | Note | *P*-value by LRT comparing with null model (DF =4) |
|---|---|---|---|---|---|---|---|---|
| 102 | 19.6275(3.1920) | 1.6468(0.9220) | -4.5356(1.3220) | -0.1943(0.5102) | -0.9667(1.3264) | 103.7767(3.3897) | Excluded | |
| 78 | 0.4053(0.0676) | 0.0247(0.0216) | 0.0695(0.0255) | -0.0067(0.0074) | 0.0346(0.0232) | 1.0682(0.0679) | Excluded | |
| 3144 | 84.8875(16.0285) | 6.4843(5.3411) | 18.3183(6.1430) | -1.0605(1.8600) | 10.6128(5.5819) | 269.2638(16.2269) | Excluded | |
| 3147 | 127.1609(21.2069) | 7.7508(6.7659) | 21.8139(7.9958) | -2.0979(2.3355) | 10.8653(7.2771) | 335.1489(21.2930) | Excluded | |
| 3148 | 0.0048(0.0008) | 0.0003(0.0003) | 0.0009(0.0003) | -0.0001(0.0001) | 0.0004(0.0003) | 0.0135(0.0008) | Excluded | |
| 1160 | 0.0824(0.0267) | 0.0061(0.0084) | -0.0017(0.0119) | 0.0213(0.0073) | 0.0145(0.0130) | 1.0426(0.0297) | | 0.0003 |
| 20160 | 0.0304(0.0058) | 0.0043(0.0018) | -0.0076(0.0024) | 0.0005(0.0009) | 0.0009(0.0024) | 0.1971(0.0063) | | 0.0005 |
| 1379 | 0.0276(0.0116) | -0.0051(0.0030) | 0.0235(0.0048) | 0.0065(0.0028) | 0.0085(0.0053) | 0.4801(0.0129) | Excluded | |
| 2139 | 1.4745(0.4017) | 0.1370(0.1306) | 0.0349(0.1669) | 0.3112(0.1006) | 0.4877(0.1743) | 13.1333(0.4386) | | 0.0002 |
| 6138 | 0.1751(0.0223) | 0.0619(0.0115) | -0.0237(0.0111) | 0.0262(0.0064) | 0.0816(0.0105) | 0.7579(0.0246) | Excluded | |

**Table S12**. **Comparison of LRT significance results with/without additional confounders for ten phenotypes with significant interaction across POP1+POP2.** In bold we marked the significant ones by $P < 3.57E-4$ after Bonferroni correction. The estimates which were not within the valid parameter space were excluded. DF denotes degree of freedom.

| UKBB data-field | Phenotypes | *P*-value by LRT (G×P RNM with PC1 vs. null model) (DF =2) for phenotypes adjusted by basic confounders | *P*-value by LRT (G×P RNM with PC1 vs. null model) (DF=2) for phenotypes adjusted by basic and additional confounders | *P*-value by LRT (G×P RNM with PC2 vs. null model) (DF=2) for phenotypes adjusted by basic confounders | *P*-value by LRT (G×P RNM with PC2 vs. null model) (DF =2) for phenotypes adjusted by basic and additional confounders |
|---|---|---|---|---|---|
| 102 | Pulse rate, automated reading | **0.0002** | 0.0021 | 0.0308 | 0.0887 |
| 78 | Heel bone mineral density (BMD) T-score, automated | **9.39E-07** | **1.33E-08** | Excluded | Excluded |
| 3144 | Heel Broadband ultrasound attenuation, direct entry | **2.87E-07** | **1.70E-07** | Excluded | Excluded |
| 3147 | Heel quantitative ultrasound index (QUI), direct entry | **9.40E-07** | **1.33E-08** | Excluded | Excluded |
| 3148 | Heel bone mineral density (BMD) | **1.58E-07** | **2.19E-09** | Excluded | Excluded |
| 1160 | Sleep duration | 0.0073 | 0.2187 | **0.0001** | 0.0179 |
| 20160 | Ever smoked | **0.0001** | 0.6531 | 0.0072 | 0.9341 |
| 1379 | Lamb/mutton intake | Excluded | Excluded | **2.73E-05** | 0.0056 |
| 2139 | Age first had sexual intercourse | 0.0047 | 0.0010 | **0.0001** | **0.0001** |
| 6138 | Qualifications | **1.96E-18** | **1.21E-12** | **3.22E-23** | **3.76E-19** |

Notes: the basic confounders of fixed effects are sex, year of birth, age at recruitment, genotype measurement batch, UKB assessment centre and the first 20 ancestry PCs provided by UKB.

For phenotypes data-field 102, additional confounders are: “Townsend deprivation index at recruitment (data-field 189)”, “major dietary changes in the last 5 years (data-field 1538)”, “variation in diet (data-field 1548)”, “smoking status (data-field 20116)” and “alcohol drinker status (data-field 20117)”.

For phenotypes data-field 78, 3144, 3147, 3148, additional confounders are: “Townsend deprivation index at recruitment (data-field 189)”, “ankle spacing width (data-field 3143)”, “heel ultrasound method (data-field 19)” and “foot measured for bone density (data-field 3081)”.

For phenotypes data-field 1160, additional confounders are: “Townsend deprivation index at recruitment (data-field 189)”, “residential noise pollution (data-field: 24020-24024)”, “type of accommodation lived in (data-field 670)”, “number in household (data-field 709)”, “current employment status (data-field 6142)”, “morning/evening person (chronotype) (data-field 1180)”, “sleeplessness / insomnia (data-field 1200)” and “daytime dozing/sleeping (narcolepsy) (data-Field 1220)”.

For phenotype data-field 20160, additional confounder is: “Townsend deprivation index at recruitment (data-field 189)”, “exposure to tobacco smoke at home (data-field 1269)” and “exposure to tobacco smoke outside home (data-field 1279)”.

For phenotype data-field 1379, additional confounders are: “Townsend deprivation index at recruitment (data-field 189)”, “major dietary changes in the last 5 years (data-field 1538)”, “variation in diet (data-field 1548)”, “alcohol drinker status (data-field 20117)” and “average total household income before tax (data-field: 738)”.

For phenotype data-field 2139, additional confounders are: “Townsend deprivation index at recruitment (data-field 189)” and “qualifications (data-field 6138)”.

For phenotype data-field 6138, additional confounders are: “Townsend deprivation index at recruitment (data-field: 189)”, “current employment status (data-field: 6142)” and “average total household income before tax (data-field: 738)”.

**Table S13**. **Results of model comparisons (baseline, G×P/R×P and full) through LRT for two phenotypes across POP1+POP2 with the covariates PC1 and PC2**. The phenotypes were adjusted by basic plus additional confounders of fixed effects and transformed using rank-based INT. DF denotes degree of freedom.

| UKBB data field | Phenotype | Covariate | Model comparison | DF | P-value by LRT |
|---|---|---|---|---|---|
| 2139 | Age first had sexual intercourse | PC1 | Full versus R×P | 2 | 0.8770 |
| | | | Full versus G×P | 2 | 0.3778 |
| | | | Full versus GREML | 4 | 0.0002 |
| | | | R×P versus GREML | 2 | 2.22E-05 |
| | | | G×P versus GREML | 2 | 5.16E-05 |
| | | PC2 | Full versus R×P | 2 | 0.9247 |
| | | | Full versus G×P | 2 | 0.9901 |
| | | | Full versus GREML | 4 | 0.0484 |
| | | | R×P versus GREML | 2 | 0.0131 |
| | | | G×P versus GREML | 2 | 0.0097 |
| 6138 | Qualifications | PC1 | Full versus R×P | 2 | 4.15E-05 |
| | | | Full versus G×P | 2 | 5.09E-05 |
| | | | Full versus GREML | 4 | 2.35E-20 |
| | | | R×P versus GREML | 2 | 1.13E-17 |
| | | | G×P versus GREML | 2 | 9.21E-18 |
| | | PC2 | Full versus R×P | 2 | 0.0032 |
| | | | Full versus G×P | 2 | 3.39E-09 |
| | | | Full versus GREML | 4 | 5.64E-31 |
| | | | R×P versus GREML | 2 | 2.36E-30 |
| | | | G×P versus GREML | 2 | 2.22E-24 |

**Table S14**. **Genetic variance, interaction variance and their covariance component estimates for 58 UKBB phenotypes with significant heritabilities across POP2+POP3 with the covariate PC1.** The estimates which were not within the valid parameter space are excluded for follow-up analyses as shown in the column "Note" (in bold we marked the significant ones by P < 0.05/116 = 0.0004 after Bonferroni correction). SE denotes standard error. DF denotes degree of freedom.

| UKBB data-field | $\mathrm{var}(\mathbf{g}_0)$ (SE) | $\mathrm{var}(\mathbf{g}_1)$ (SE) | $\mathrm{cov}(\mathbf{g}_0, \mathbf{g}_1)$ (SE) | $\mathrm{var}(\mathbf{e}_0)$ (SE) | Note | *P*-value by LRT comparing with null model (DF=2) |
|---|---|---|---|---|---|---|
| 48 | 29.3531(3.4716) | 3.9198(1.0112) | -6.8481(1.1346) | 108.7908(3.6147) | | **6.92E-09** |
| 49 | 17.7361(2.1585) | 1.4819(0.5692) | -2.9198(0.6875) | 68.3066(2.2444) | | **0.0001** |
| 50 | 19.3685(1.0300) | 0.2300(0.2268) | -0.1071(0.3125) | 19.7821(0.9359) | | 0.4506 |
| 20015 | 4.7694(0.3326) | -0.0584(0.0696) | 0.1839(0.0995) | 8.3267(0.3186) | Excluded | |
| 21001 | 5.2187(0.5645) | 0.4888(0.1602) | -0.6964(0.1836) | 17.1518(0.5808) | | 0.0004 |
| 21002 | 50.4472(4.9668) | 5.9372(1.4506) | -9.2975(1.6187) | 142.5020(5.0576) | | **4.05E-08** |
| 102 | 14.4839(2.9487) | 1.3990(0.8005) | -2.6871(0.9507) | 104.7696(3.1923) | | 0.0166 |
| 4079 | 15.4457(2.7790) | -0.2765(0.6064) | 0.1227(0.8651) | 95.0684(2.9504) | Excluded | |
| 4080 | 40.1608(7.6905) | -2.7447(1.4987) | 2.9719(2.3585) | 272.5969(8.2202) | Excluded | |
| 78 | 0.3206(0.0671) | 0.0480(0.0204) | 0.0252(0.0187) | 1.1281(0.0692) | | **3.76E-07** |
| 3143 | 4.4681(0.7193) | -0.0833(0.1491) | 0.4781(0.1809) | 10.9806(0.7188) | Excluded | |
| 3144 | 71.8120(16.1704) | 9.3888(4.7703) | 8.6593(4.4874) | 279.7071(16.6966) | | **1.83E-07** |
| 3147 | 100.5781(21.0580) | 15.0530(6.3924) | 7.9177(5.8529) | 353.9731(21.7207) | | **3.77E-07** |
| 3148 | 0.0041(0.0008) | 0.0006(0.0003) | 0.0003(0.0002) | 0.0140(0.0009) | | **2.47E-07** |
| 1558 | 0.2557(0.0542) | 0.0039(0.0123) | 0.0041(0.0174) | 2.0676(0.0588) | | 0.7806 |
| 1598 | 2.5854(0.7757) | 0.4771(0.1750) | -1.4140(0.2662) | 29.4564(0.8877) | Excluded | |
| 1628 | 0.0160(0.0078) | 0.0019(0.0023) | -0.0096(0.0029) | 0.4396(0.0096) | Excluded | |
| 1309 | 0.0197(0.0364) | -0.0244(0.0078) | 0.0458(0.0123) | 2.1924(0.0423) | Excluded | |
| 1319 | 0.0391(0.0589) | -0.0319(0.0136) | 0.0850(0.0183) | 3.1448(0.0681) | Excluded | |
| 1329 | 0.0384(0.0180) | 0.0072(0.0050) | -0.0111(0.0061) | 0.7627(0.0204) | | 0.1723 |
| 1339 | 0.0299(0.0140) | 0.0053(0.0041) | -0.0032(0.0047) | 0.5921(0.0159) | | 0.3751 |
| 1349 | 0.0996(0.0269) | 0.0055(0.0062) | -0.0116(0.0088) | 1.0696(0.0296) | | 0.4266 |
| 1478 | 0.0820(0.0188) | -0.0020(0.0046) | 0.0304(0.0063) | 0.7340(0.0205) | Excluded | |
| 1518 | 0.0326(0.0084) | -0.0041(0.0014) | 0.0075(0.0025) | 0.3350(0.0091) | Excluded | |
| 1528 | 0.7051(0.1586) | 0.0826(0.0457) | -0.0074(0.0530) | 5.9796(0.1736) | | 0.0261 |

| | | | | | | |
|---|---|---|---|---|---|---|
| 1548 | 0.0222(0.0085) | 0.0002(0.0021) | -0.0015(0.0028) | 0.3542(0.0095) | | 0.8026 |
| 1110 | 0.1277(0.0347) | 0.0109(0.0089) | -0.0479(0.0121) | 1.5226(0.0383) | Excluded | |
| 1130 | 0.0419(0.0188) | 0.0106(0.0060) | 0.0244(0.0061) | 0.6486(0.0206) | Excluded | |
| 2237 | 0.0115(0.0057) | 0.0016(0.0017) | 0.0011(0.0019) | 0.2372(0.0064) | | 0.1631 |
| 2139 | 1.0829(0.3699) | 0.4007(0.1218) | -0.2017(0.1282) | 13.1538(0.4106) | | **0.0002** |
| 1160 | 0.0879(0.0262) | 0.0390(0.0098) | -0.0188(0.0093) | 1.0070(0.0292) | | **2.14E-05** |
| 1170 | 0.0772(0.0152) | 0.0028(0.0040) | 0.0007(0.0050) | 0.5661(0.0164) | | 0.5017 |
| 1180 | 0.1361(0.0252) | 0.0001(0.0058) | -0.0012(0.0079) | 0.8538(0.0269) | | 0.9831 |
| 1190 | 0.0164(0.0076) | 0.0052(0.0022) | -0.0057(0.0026) | 0.3099(0.0086) | | 0.0273 |
| 1200 | 0.0248(0.0116) | -0.0049(0.0025) | 0.0084(0.0037) | 0.4974(0.0130) | Excluded | |
| 1249 | 0.1317(0.0384) | -0.0043(0.0078) | 0.0063(0.0118) | 1.3732(0.0417) | Excluded | |
| 20116 | 0.0385(0.0114) | 0.0006(0.0029) | 0.0072(0.0037) | 0.4427(0.0125) | Excluded | |
| 1717 | 0.0740(0.0095) | 0.0018(0.0026) | 0.0272(0.0031) | 0.3165(0.0098) | Excluded | |
| 1687 | 0.0206(0.0029) | -0.0003(0.0007) | 0.0022(0.0009) | 0.0970(0.0030) | Excluded | |
| 1697 | 0.0292(0.0029) | 0.0010(0.0007) | -0.0017(0.0009) | 0.0828(0.0029) | | 0.1817 |
| 2714 | 0.5038(0.0998) | -0.0131(0.0172) | 0.0203(0.0266) | 2.0599(0.1023) | Excluded | |
| 2734 | 0.1687(0.0522) | 0.0196(0.0120) | -0.0232(0.0144) | 1.2134(0.0552) | | 0.1677 |
| 2744 | 0.2291(0.0878) | 0.0000(0.0153) | 0.0221(0.0215) | 1.4305(0.0908) | Excluded | |
| 2375 | 0.1770(0.0141) | 0.0431(0.0138) | 0.0028(0.0029) | -0.0003(0.0031) | | 0.3597 |
| 2395 | 0.4911(0.0708) | 0.0077(0.0129) | -0.0036(0.0154) | 0.6604(0.0689) | | 0.7761 |
| 1940 | 0.0108(0.0050) | -0.0004(0.0012) | 0.0018(0.0016) | 0.2029(0.0056) | Excluded | |
| 1960 | 0.0139(0.0053) | -0.0004(0.0012) | 0.0015(0.0017) | 0.2157(0.0059) | Excluded | |
| 2000 | 0.0117(0.0058) | -0.0004(0.0013) | 0.0008(0.0019) | 0.2336(0.0065) | Excluded | |
| 20127 | 0.5977(0.2965) | 0.0003(0.0669) | 0.0382(0.0910) | 9.7643(0.3257) | Excluded | |
| 2070 | 0.0295(0.0106) | -0.0022(0.0030) | 0.0201(0.0036) | 0.4233(0.0118) | Excluded | |
| 2080 | 0.0356(0.0169) | 0.0114(0.0053) | -0.0031(0.0059) | 0.6828(0.0191) | | 0.0148 |
| 404 | 12441.93(4683.00) | -1523.68(964.68) | 7665.23(1146.36) | 75961.29(4814.57) | Excluded | |
| 2217 | 47.6927(7.7861) | -1.8024(1.2492) | -2.3755(2.1523) | 222.6492(8.0704) | Excluded | |
| 2247 | 0.0099(0.0037) | 0.0000(0.0009) | -0.0034(0.0012) | 0.1617(0.0042) | Excluded | |
| 2443 | 0.0034(0.0010) | 0.0030(0.0005) | 0.0003(0.0004) | 0.0357(0.0011) | | **4.88E-29** |
| 6138 | 0.1327(0.0205) | 0.0398(0.0081) | -0.0001(0.0072) | 0.6825(0.0219) | | **1.81E-13** |
| 699 | 5.9094(1.9164) | 0.4690(0.4016) | -2.1491(0.6147) | 84.6077(2.1306) | Excluded | |
| 738 | 0.0738(0.0357) | -0.0053(0.0086) | 0.0348(0.0112) | 1.2983(0.0393) | Excluded | |

**Table S15. Genetic variance, interaction variance and their covariance component estimates for 58 UKBB phenotypes with significant heritabilities across POP2+POP3 with the covariate PC2.** The estimates which were not within the valid parameter space are excluded for follow-up analyses as shown in the column "Note" (in bold we marked the significant ones by P < 0.05/116 = 0.0004 after Bonferroni correction). SE denotes standard error. DF denotes degree of freedom.

| UKB data-field | $\text{var}(\mathbf{g}_0)$ (SE) | $\text{var}(\mathbf{g}_1)$ (SE) | $\text{cov}(\mathbf{g}_0, \mathbf{g}_1)$ (SE) | $\text{var}(\mathbf{e}_0)$ (SE) | Note | *P*-value by LRT comparing with null model (DF =2) |
|---|---|---|---|---|---|---|
| 48 | 29.2708(3.4812) | 0.7248(0.4715) | 4.0443(1.0811) | 112.0167(3.5482) | | **0.0002** |
| 49 | 17.6501(2.1581) | 0.5207(0.3017) | 2.3922(0.6738) | 69.3360(2.2038) | | 0.0011 |
| 50 | 19.4296(1.0298) | -0.0555(0.1421) | -0.3130(0.3197) | 20.0123(0.9239) | Excluded | |
| 20015 | 4.7441(0.3326) | 0.0380(0.0521) | -0.0340(0.1065) | 8.2553(0.3156) | | 0.3739 |
| 21001 | 5.2462(0.5654) | 0.0856(0.0771) | 0.4591(0.1765) | 17.5269(0.5703) | | 0.0203 |
| 21002 | 50.3179(4.9792) | 1.2034(0.7069) | 4.8923(1.5357) | 147.2704(4.9679) | | 0.0047 |
| 102 | 14.4908(2.9472) | 0.5656(0.4550) | 1.7582(0.9310) | 105.5717(3.1365) | | 0.1565 |
| 4079 | 15.3282(2.7745) | 0.3065(0.3380) | 1.5207(0.8435) | 94.6158(2.9111) | | 0.1976 |
| 4080 | 40.0402(7.6878) | -0.7162(0.8887) | -2.4181(2.4035) | 270.6507(8.1186) | Excluded | |
| 78 | 0.3100(0.0677) | -0.0042(0.0064) | -0.0094(0.0160) | 1.1911(0.0685) | Excluded | |
| 3143 | 4.4414(0.7217) | -0.1123(0.0607) | -0.2243(0.1658) | 11.0458(0.7144) | Excluded | |
| 3144 | 70.1760(16.3260) | -0.9640(1.5654) | -3.2945(3.8860) | 291.8513(16.5557) | Excluded | |
| 3147 | 97.2343(21.2319) | -1.3135(1.9993) | -2.9516(5.0193) | 373.7378(21.4770) | Excluded | |
| 3148 | 0.0040(0.0008) | -0.0001(0.0001) | -0.0001(0.0002) | 0.0148(0.0009) | Excluded | |
| 1558 | 0.2557(0.0540) | 0.0194(0.0095) | 0.0409(0.0179) | 2.0525(0.0581) | | 0.0473 |
| 1598 | 3.8801(0.7132) | 5.1373(0.5082) | -5.9721(0.3492) | 22.1056(0.6559) | Excluded | |
| 1628 | 0.0309(0.0126) | 0.0030(0.0019) | 0.0083(0.0040) | 0.4714(0.0138) | | 0.1290 |
| 1309 | 0.0410(0.0369) | -0.0179(0.0045) | -0.0739(0.0120) | 2.2874(0.0433) | Excluded | |
| 1319 | 0.2761(0.0907) | 0.3954(0.0490) | -0.1803(0.0353) | 3.5394(0.0993) | | **1.73E-78** |
| 1329 | 0.0381(0.0179) | 0.0035(0.0030) | 0.0118(0.0059) | 0.7667(0.0200) | Excluded | |
| 1339 | 0.0289(0.0140) | 0.0054(0.0029) | 0.0068(0.0048) | 0.5931(0.0157) | | 0.1734 |
| 1349 | 0.1006(0.0269) | -0.0105(0.0030) | -0.0254(0.0088) | 1.0845(0.0292) | Excluded | |
| 1478 | 0.0803(0.0189) | 0.0006(0.0033) | -0.0136(0.0064) | 0.7334(0.0204) | Excluded | |
| 1518 | 0.0318(0.0084) | -0.0006(0.0012) | -0.0037(0.0027) | 0.3324(0.0091) | Excluded | |
| 1528 | 0.7688(0.1583) | 0.1635(0.0367) | 0.2872(0.0542) | 5.8442(0.1708) | | **2.45E-07** |
| 1548 | 0.0222(0.0085) | 0.0006(0.0014) | 0.0012(0.0029) | 0.3538(0.0094) | | 0.9130 |

| | | | | | | |
|---|---|---|---|---|---|---|
| 1110 | 0.1398(0.0327) | 0.0191(0.0062) | 0.0634(0.0113) | 1.5020(0.0368) | Excluded | |
| 1130 | 0.0361(0.0189) | 0.0031(0.0035) | -0.0057(0.0060) | 0.6625(0.0206) | | 0.0558 |
| 2237 | 0.0120(0.0057) | -0.0012(0.0006) | -0.0016(0.0017) | 0.2397(0.0063) | Excluded | |
| 2139 | 1.0287(0.3688) | 0.3196(0.0861) | 0.3142(0.1295) | 13.2980(0.4027) | | **0.0001** |
| 1160 | 0.0903(0.0262) | 0.0302(0.0067) | 0.0249(0.0095) | 1.0139(0.0286) | | **2.26E-07** |
| 1170 | 0.0783(0.0152) | -0.0005(0.0024) | -0.0080(0.0051) | 0.5683(0.0162) | Excluded | |
| 1180 | 0.1367(0.0252) | -0.0025(0.0032) | -0.0089(0.0078) | 0.8557(0.0265) | Excluded | |
| 1190 | 0.0164(0.0076) | 0.0006(0.0012) | 0.0013(0.0025) | 0.3145(0.0084) | | 0.8662 |
| 1200 | 0.0250(0.0117) | 0.0002(0.0017) | -0.0000(0.0039) | 0.4922(0.0130) | | 0.9858 |
| 1249 | 0.1332(0.0384) | -0.0064(0.0041) | -0.0053(0.0116) | 1.3741(0.0413) | Excluded | |
| 20116 | 0.0395(0.0114) | 0.0031(0.0020) | -0.0018(0.0038) | 0.4393(0.0124) | | 0.0186 |
| 1717 | 0.0653(0.0096) | 0.0024(0.0020) | -0.0065(0.0033) | 0.3243(0.0100) | | 0.0020 |
| 1687 | 0.0208(0.0029) | -0.0005(0.0003) | -0.0015(0.0009) | 0.0970(0.0030) | Excluded | |
| 1697 | 0.0292(0.0029) | 0.0000(0.0004) | 0.0004(0.0009) | 0.0837(0.0029) | Excluded | |
| 2714 | 0.5008(0.0998) | 0.0003(0.0128) | -0.0142(0.0272) | 2.0492(0.1016) | Excluded | |
| 2734 | 0.1625(0.0517) | 0.0164(0.0074) | 0.0425(0.0144) | 1.2231(0.0541) | | 0.0153 |
| 2744 | 0.2304(0.0878) | 0.0110(0.0120) | 0.0131(0.0215) | 1.4189(0.0902) | | 0.6456 |
| 2375 | 0.0434(0.0138) | 0.0004(0.0015) | -0.0016(0.0031) | 0.1792(0.0139) | | 0.5309 |
| 2395 | 0.4938(0.0708) | -0.0002(0.0068) | 0.0026(0.0155) | 0.6658(0.0683) | Excluded | |
| 1940 | 0.0104(0.0050) | -0.0005(0.0006) | -0.0007(0.0016) | 0.2034(0.0055) | Excluded | |
| 1960 | 0.0134(0.0053) | 0.0004(0.0008) | 0.0019(0.0018) | 0.2153(0.0058) | | 0.5748 |
| 2000 | 0.0118(0.0057) | 0.0005(0.0009) | 0.0014(0.0019) | 0.2327(0.0064) | | 0.7379 |
| 20127 | 0.5881(0.2963) | 0.0198(0.0407) | 0.0543(0.0914) | 9.7560(0.3221) | | 0.8401 |
| 2070 | 0.0280(0.0106) | -0.0027(0.0016) | -0.0108(0.0033) | 0.4255(0.0117) | Excluded | |
| 2080 | 0.0357(0.0169) | 0.0062(0.0031) | 0.0078(0.0057) | 0.6881(0.0187) | | 0.0739 |
| 404 | 10582.50(4798.56) | 2663.16(877.46) | 1545.96(1166.36) | 73836.10(4949.27) | | 0.0004 |
| 2217 | 47.5311(7.7886) | -0.3026(0.8634) | 2.2033(2.2680) | 221.2215(8.0147) | Excluded | |
| 2247 | 0.0054(0.0025) | -0.0001(0.0003) | 0.0022(0.0008) | 0.1429(0.0031) | Excluded | |
| 2443 | 0.0037(0.0010) | 0.0015(0.0003) | 0.0006(0.0004) | 0.0368(0.0011) | | **1.15E-11** |
| 6138 | 0.1356(0.0206) | 0.0211(0.0049) | 0.0230(0.0073) | 0.6986(0.0216) | | **9.21E-06** |
| 699 | 3.3450(1.4937) | 0.3436(0.2089) | 1.9081(0.5026) | 83.7636(1.7889) | Excluded | |
| 738 | 0.0800(0.0358) | 0.0043(0.0061) | -0.0168(0.0112) | 1.2832(0.0390) | | 0.0239 |

**Table S16**. **The G×P interaction analysis results for 13 phenotypes across POP2+POP3.** The phenotypic values were adjusted by basic plus additional confounders of fixed effects and transformed by rank-based INT (in bold we marked the significant ones by P < 0.05/116 = 0.0004). The estimates which were not within the valid parameter space are excluded. SE denotes standard error. DF denotes degree of freedom.

| UKBB data-field | Phenotypes | *P*-value by LRT (G×P RNM with PC1 vs. baseline model) (DF =2) | *P*-value by LRT (G×P RNM with PC2 vs. baseline model) (DF =2) |
|---|---|---|---|
| 48 | Waist circumference | **2.92E-06** | 0.0004 |
| 49 | Hip circumference | 0.0070 | 0.0011 |
| 21002 | Weight | **0.0002** | 0.0252 |
| 78 | Heel bone mineral density (BMD) T-score, automated | 0.0852 | 0.8836 |
| 3144 | Heel Broadband ultrasound attenuation, direct entry | 0.0441 | 0.8292 |
| 3147 | Heel quantitative ultrasound index (QUI), direct entry | 0.0853 | 0.8836 |
| 3148 | Heel bone mineral density (BMD) | 0.0782 | 0.8718 |
| 1528 | Water intake | 0.7586 | 0.8449 |
| 1319 | Dried fruit intake | Excluded | Excluded |
| 1160 | Sleep duration | 0.0703 | 0.0840 |
| 2443 | Diabetes diagnosed by doctor | **6.65E-11** | **3.73E-08** |
| 2139 | Age first had sexual intercourse | **7.86E-05** | 0.0071 |
| 6138 | Qualifications | **1.06E-15** | 0.0162 |

Notes: the basic confounders are sex, year of birth, age at recruitment, genotype measurement batch, UKBB assessment centre and the first 20 ancestry PCs provided by UKBB.

For phenotypes data-fields 48, 49 and 21002, additional confounders are: “Townsend deprivation index at recruitment (data-field 189)”, “major dietary changes in the last 5 years (data-field 1538)”, “variation in diet (data-field 1548)”, “smoking status (data-field 20116)” and “alcohol drinker status (data-field 20117)”.

For phenotypes data-fields 78, 3144, 3147, 3148, additional confounders are: “Townsend deprivation index at recruitment (data-field 189)”, “ankle spacing width (data-field 3143)”, “heel ultrasound method (data-field 19)” and “foot measured for bone density (data-field 3081)”.

For phenotypes data-fields 1528 and 1319, additional confounders are: “Townsend deprivation index at recruitment (data-field: 189)”, “major dietary changes in the last 5 years (data-field: 1538)”, “variation in diet (data-field: 1548)” and “Vitamin and mineral supplements (data-field: 6155)”.

For phenotypes data-field 1160, additional confounders are: “residential noise pollution (data-field: 24020-24024)”, “type of accommodation lived in (data-field 670)”, “number in household (data-field 709)”, “current employment status (data-field 6142)”, “morning/evening person (chronotype) (data-field 1180)”, “sleeplessness / insomnia (data-field 1200)” and “daytime dozing/sleeping (narcolepsy) (data-field 1220)”.

For phenotypes data-fields 2443, additional confounders are: “Townsend deprivation index at recruitment (data-field 189)”, “major dietary changes in the last 5 years (data-field 1538)”, “variation in diet (data-field 1548)”, “smoking status (data-field 20116)” and “alcohol drinker status (data-field 20117)”.

For phenotype data-field 2139, additional confounders are: “Townsend deprivation index at recruitment (data-field 189)” and “qualifications (data-field 6138)”.

For phenotype data-field 6138, additional confounders are: “Townsend deprivation index at recruitment (data-field: 189)”, “current employment status (data-field: 6142)” and “average total household income before tax (data-field: 738)”.

**Table S17**. **Results of model comparisons (baseline, G×P/R×P and full) through LRT for two phenotypes across POP2+POP3 with the covariates PC1 and PC2**. The phenotypes were adjusted by basic plus additional confounders of fixed effects and transformed using rank-based INT. The estimates of full model for age first had sexual intercourse were not within the valid parameter space and thus were excluded. DF denotes degree of freedom.

| UKBB data field | Phenotype | Covariate | Model comparison | DF | P-value by LRT |
|---|---|---|---|---|---|
| 2139 | Age first had sexual intercourse | PC1 | Full versus R×P | 2 | Excluded |
| | | | Full versus G×P | 2 | Excluded |
| | | | Full versus GREML | 4 | Excluded |
| | | | R×P versus GREML | 2 | 0.0002 |
| | | | G×P versus GREML | 2 | 7.86E-05 |
| | | PC2 | Full versus R×P | 2 | 0.4374 |
| | | | Full versus G×P | 2 | 0.8363 |
| | | | Full versus GREML | 4 | 0.0363 |
| | | | R×P versus GREML | 2 | 0.0135 |
| | | | G×P versus GREML | 2 | 0.0071 |
| 6138 | Qualifications | PC1 | Full versus R×P | 2 | 0.0031 |
| | | | Full versus G×P | 2 | 0.2166 |
| | | | Full versus GREML | 4 | 8.53E-15 |
| | | | R×P versus GREML | 2 | 7.45E-14 |
| | | | G×P versus GREML | 2 | 1.06E-15 |
| | | PC2 | Full versus R×P | 2 | 0.0053 |
| | | | Full versus G×P | 2 | 0.0090 |
| | | | Full versus GREML | 4 | 0.0014 |
| | | | R×P versus GREML | 2 | 0.0273 |
| | | | G×P versus GREML | 2 | 0.0162 |

**Table S18**. **Genetic variance, interaction variance and their covariance component estimates for 70 UKBB phenotypes with significant heritability across POP1+POP3 with the covariate PC1.** The estimates which were not within the valid parameter space were excluded for follow-up analyses as shown in the column "Note" (in bold we marked the significant ones by P < 0.05/140 = 3.57E-4 after Bonferroni correction). SE denotes standard error. DF denotes degree of freedom.

| UKBB data-field | $\mathrm{var}(\mathbf{g}_0)$ (SE) | $\mathrm{var}(\mathbf{g}_1)$ (SE) | $\mathrm{cov}(\mathbf{g}_0,\mathbf{g}_1)$ (SE) | $\mathrm{var}(\mathbf{e}_0)$ (SE) | Note | *P*-value by LRT comparing with null model (DF =2) |
|---|---|---|---|---|---|---|
| 48 | 26.9732(3.4423) | -3.0381(1.0703) | -0.3651(0.8118) | 123.2641(3.7130) | Excluded | |
| 49 | 18.9254(2.0988) | -1.5711(0.6476) | 0.1017(0.4952) | 71.9214(2.2336) | Excluded | |
| 50 | 21.3636(0.9865) | -0.0227(0.2920) | -0.2788(0.2303) | 17.6890(0.9035) | Excluded | |
| 20015 | 5.0084(0.3254) | -0.0881(0.0982) | -0.2779(0.0776) | 8.3251(0.3215) | Excluded | |
| 21001 | 5.0703(0.5535) | -0.5280(0.1655) | -0.1188(0.1288) | 18.9252(0.5864) | Excluded | |
| 21002 | 49.9582(4.9475) | -4.6811(1.4725) | -1.4562(1.1500) | 162.5982(5.1971) | Excluded | |
| 102 | 19.3238(3.1377) | -1.1396(1.0436) | -0.4463(0.7590) | 112.5584(3.4496) | Excluded | |
| 4079 | 15.4216(2.6480) | -0.0335(0.8990) | -0.7379(0.6418) | 93.5503(2.9202) | Excluded | |
| 4080 | 44.1031(7.8598) | 1.9094(2.7565) | -1.6888(1.9399) | 277.0338(8.7000) | | 0.6187 |
| 78 | 0.4320(0.0586) | 0.0259(0.0167) | -0.0013(0.0116) | 0.9662(0.0597) | | 0.2742 |
| 3143 | 3.4326(0.6153) | -0.4373(0.1524) | 0.0428(0.1118) | 12.0823(0.6441) | Excluded | |
| 3144 | 105.0637(13.91) | 6.8594(4.03) | 0.9229(2.78) | 229.8112(14.19) | | 0.1690 |
| 3147 | 135.5514(18.39) | 8.1333(5.23) | -0.4124(3.63) | 303.1400(18.73) | | 0.2740 |
| 3148 | 0.0055(0.0007) | 0.0004(0.0002) | 0.0000(0.0001) | 0.0118(0.0007) | | 0.1049 |
| 1558 | 0.2076(0.0490) | -0.0113(0.0178) | 0.0099(0.0125) | 2.0548(0.0561) | Excluded | |
| 1309 | 0.1409(0.0531) | 0.2320(0.0261) | 0.1177(0.0160) | 2.1870(0.0613) | | **3.31E-47** |
| 1319 | 0.2023(0.0722) | -0.2698(0.0118) | 0.1448(0.0151) | 3.7380(0.0821) | Excluded | |
| 1329 | 0.0483(0.0172) | -0.0014(0.0064) | -0.0109(0.0046) | 0.7691(0.0201) | Excluded | |
| 1339 | 0.0278(0.0132) | -0.0032(0.0048) | -0.0030(0.0034) | 0.5913(0.0154) | Excluded | |
| 1349 | 0.0620(0.0229) | 0.0154(0.0091) | 0.0146(0.0061) | 0.9958(0.0266) | | 0.0037 |
| 1359 | 0.0438(0.0174) | -0.0038(0.0065) | 0.0146(0.0045) | 0.7825(0.0204) | Excluded | |
| 1369 | 0.0320(0.0152) | 0.0030(0.0057) | -0.0015(0.0039) | 0.6643(0.0176) | | 0.8446 |
| 1379 | 0.0228(0.0106) | -0.0036(0.0039) | 0.0002(0.0028) | 0.4814(0.0125) | Excluded | |
| 1408 | 0.0565(0.0261) | 0.0174(0.0102) | 0.0082(0.0068) | 1.1149(0.0305) | | 0.0534 |
| 1458 | 0.5171(0.1693) | 0.0346(0.0648) | 0.0042(0.0446) | 7.3911(0.1974) | | 0.8447 |

| | | | | | | |
|---|---|---|---|---|---|---|
| 1478 | 0.0627(0.0165) | -0.0003(0.0061) | 0.0015(0.0042) | 0.6861(0.0189) | Excluded | |
| 1518 | 0.0301(0.0073) | 0.0064(0.0029) | 0.0002(0.0019) | 0.2976(0.0083) | | 0.0544 |
| 1528 | 0.3581(0.1246) | 0.0089(0.0441) | 0.0939(0.0317) | 5.3408(0.1427) | Excluded | |
| 1110 | 0.1215(0.0381) | -0.0065(0.0139) | -0.0040(0.0097) | 1.6230(0.0438) | Excluded | |
| 2237 | 0.0175(0.0056) | -0.0004(0.0021) | -0.0037(0.0015) | 0.2411(0.0065) | Excluded | |
| 2139 | 2.1102(0.3636) | -0.0637(0.1198) | -0.1399(0.0873) | 12.4525(0.3993) | Excluded | |
| 1160 | 0.0885(0.0248) | -0.0125(0.0089) | -0.0169(0.0064) | 1.0712(0.0286) | Excluded | |
| 1170 | 0.0707(0.0134) | 0.0013(0.0048) | 0.0070(0.0034) | 0.5228(0.0150) | | 0.0893 |
| 1180 | 0.1505(0.0234) | -0.0002(0.0079) | 0.0053(0.0056) | 0.7792(0.0255) | Excluded | |
| 1190 | 0.0320(0.0076) | -0.0024(0.0027) | -0.0028(0.0019) | 0.3108(0.0086) | Excluded | |
| 1200 | 0.0499(0.0112) | -0.0008(0.0041) | -0.0028(0.0029) | 0.4608(0.0128) | Excluded | |
| 1210 | 0.0165(0.0052) | -0.0018(0.0018) | -0.0008(0.0013) | 0.2037(0.0059) | Excluded | |
| 1239 | 0.0098(0.0038) | 0.0021(0.0015) | 0.0018(0.0010) | 0.1643(0.0044) | | 0.0322 |
| 1249 | 0.1658(0.0433) | -0.0255(0.0141) | -0.0368(0.0105) | 1.6251(0.0489) | Excluded | |
| 20116 | 0.0564(0.0109) | -0.0015(0.0038) | -0.0059(0.0027) | 0.4305(0.0123) | Excluded | |
| 20160 | 0.0214(0.0053) | -0.0000(0.0019) | -0.0036(0.0014) | 0.2098(0.0060) | Excluded | |
| 1717 | 0.0606(0.0078) | 0.0020(0.0027) | 0.0046(0.0019) | 0.2723(0.0084) | | 0.0252 |
| 1687 | 0.0148(0.0026) | -0.0003(0.0009) | -0.0003(0.0006) | 0.0992(0.0029) | Excluded | |
| 1697 | 0.0318(0.0027) | -0.0005(0.0009) | -0.0000(0.0007) | 0.0826(0.0028) | Excluded | |
| 20022 | 0.0482(0.0160) | 0.0091(0.0050) | -0.0099(0.0034) | 0.3664(0.0172) | | 0.0052 |
| 2714 | 0.4597(0.0983) | -0.0166(0.0280) | -0.0154(0.0196) | 2.1025(0.1048) | Excluded | |
| 2734 | 0.1398(0.0520) | -0.0207(0.0151) | -0.0069(0.0101) | 1.2594(0.0564) | Excluded | |
| 2375 | 0.0424(0.0108) | -0.0003(0.0025) | 0.0012(0.0018) | 0.1605(0.0112) | Excluded | |
| 2395 | 0.5203(0.0653) | -0.0161(0.0138) | -0.0100(0.0108) | 0.6877(0.0641) | Excluded | |
| 1920 | 0.0277(0.0054) | -0.0008(0.0019) | -0.0006(0.0014) | 0.2103(0.0060) | Excluded | |
| 1930 | 0.0141(0.0051) | -0.0002(0.0019) | -0.0003(0.0013) | 0.2214(0.0059) | Excluded | |
| 1940 | 0.0199(0.0046) | 0.0015(0.0017) | 0.0007(0.0012) | 0.1801(0.0052) | | 0.4857 |
| 1950 | 0.0131(0.0052) | 0.0008(0.0020) | 0.0012(0.0014) | 0.2258(0.0061) | | 0.5554 |
| 1960 | 0.0228(0.0051) | -0.0010(0.0019) | 0.0001(0.0013) | 0.2091(0.0058) | Excluded | |
| 1970 | 0.0101(0.0037) | -0.0002(0.0014) | 0.0028(0.0010) | 0.1627(0.0044) | Excluded | |
| 1980 | 0.0227(0.0053) | 0.0004(0.0020) | -0.0001(0.0014) | 0.2165(0.0061) | | 0.9829 |
| 1990 | 0.0170(0.0033) | 0.0030(0.0013) | 0.0027(0.0009) | 0.1275(0.0037) | | **4.17E-05** |
| 2000 | 0.0187(0.0055) | 0.0000(0.0020) | -0.0001(0.0014) | 0.2248(0.0063) | Excluded | |
| 20127 | 1.1105(0.2743) | 0.0380(0.0951) | 0.0126(0.0658) | 9.1067(0.3058) | | 0.8802 |

| | | | | | | |
|---|---|---|---|---|---|---|
| 2030 | 0.0140(0.0045) | 0.0013(0.0017) | 0.0002(0.0012) | 0.1869(0.0051) | | 0.6694 |
| 2040 | 0.0146(0.0046) | 0.0039(0.0018) | 0.0032(0.0012) | 0.1872(0.0053) | | 0.0005 |
| 2080 | 0.0550(0.0156) | -0.0074(0.0055) | -0.0010(0.0039) | 0.6471(0.0179) | Excluded | |
| 2090 | 0.0101(0.0046) | -0.0015(0.0017) | 0.0000(0.0012) | 0.2103(0.0055) | Excluded | |
| 20023 | 799.12(238.06) | -175.87(84.96) | -191.32(60.37) | 10396.08(275.15) | Excluded | |
| 2217 | 38.2853(7.0677) | -0.6904(2.3170) | 0.4299(1.6681) | 234.4942(7.7601) | Excluded | |
| 2443 | 0.0022(0.0009) | -0.0016(0.0003) | -0.0007(0.0002) | 0.0437(0.0011) | Excluded | |
| 2463 | 0.0046(0.0020) | 0.0005(0.0007) | -0.0007(0.0005) | 0.0864(0.0023) | | 0.3925 |
| 6138 | 0.1751(0.0218) | -0.0111(0.0072) | -0.0335(0.0057) | 0.8129(0.0239) | Excluded | |
| 738 | 0.0742(0.0300) | 0.0068(0.0108) | 0.0106(0.0074) | 1.1533(0.0343) | | 0.2159 |
| 135 | 0.2616(0.0697) | -0.0234(0.0244) | -0.0641(0.0179) | 2.9658(0.0800) | Excluded | |

**Table S19. Genetic variance, interaction variance and their covariance component estimates for 70 UKBB phenotypes with significant heritability across POP1+POP3 with the covariate PC2.** The estimates which were not within the valid parameter space were excluded for follow-up analyses as shown in the column "Note" (in bold we marked the significant ones by P < 0.05/140 = 3.57E-4 after Bonferroni correction). SE denotes standard error. DF denotes degree of freedom.

| UKB data-field | $\text{var}(\mathbf{g}_0)$ (SE) | $\text{var}(\mathbf{g}_1)$ (SE) | $\text{cov}(\mathbf{g}_0, \mathbf{g}_1)$ (SE) | $\text{var}(\mathbf{e}_0)$ (SE) | Note | *P*-value by LRT comparing with null model (DF =2) |
|---|---|---|---|---|---|---|
| 48 | 26.7422(3.4436) | -0.1893(1.1229) | -0.1299(0.8292) | 120.6369(3.7013) | Excluded | |
| 49 | 18.8882(2.0995) | -0.1786(0.6798) | -0.0838(0.5078) | 70.5667(2.2352) | Excluded | |
| 50 | 21.4270(0.9869) | 0.2401(0.2877) | -0.2296(0.2334) | 17.3698(0.9027) | | 0.3257 |
| 20015 | 5.0044(0.3256) | -0.0912(0.0954) | 0.1491(0.0767) | 8.3308(0.3217) | Excluded | |
| 21001 | 5.0436(0.5541) | 0.0966(0.1820) | 0.0217(0.1359) | 18.3263(0.5879) | | 0.8630 |
| 21002 | 49.7739(4.9513) | 0.8844(1.6148) | 0.7226(1.2010) | 157.2047(5.1951) | | 0.7480 |
| 102 | 19.2398(3.1360) | -0.6202(1.0479) | 0.6924(0.7641) | 112.1195(3.4616) | Excluded | |
| 4079 | 15.4377(2.6475) | 0.7679(0.8959) | 0.1859(0.6480) | 92.7287(2.9115) | | 0.6815 |
| 4080 | 43.9619(7.8596) | 1.0961(2.6020) | 0.4277(1.8981) | 277.9730(8.6237) | | 0.9043 |
| 78 | 0.4317(0.0587) | 0.0325(0.0164) | -0.0007(0.0115) | 0.9599(0.0595) | | 0.1129 |
| 3143 | 3.4977(0.6161) | 0.0292(0.1624) | 0.0787(0.1188) | 11.5526(0.6400) | | 0.7992 |
| 3144 | 104.8254(13.91) | 7.4758(3.9089) | -1.1092(2.7636) | 229.4276(14.09) | | 0.1078 |
| 3147 | 135.4496(18.40) | 10.2016(5.1359) | -0.2191(3.6159) | 301.1764(18.65) | | 0.1129 |
| 3148 | 0.0055(0.0007) | 0.0005(0.0002) | -0.0000(0.0001) | 0.0117(0.0007) | Excluded | |
| 1558 | 0.2083(0.0490) | 0.0071(0.0174) | 0.0105(0.0127) | 2.0358(0.0557) | | 0.6863 |
| 1309 | 0.1198(0.0539) | 0.1287(0.0229) | -0.0185(0.0156) | 2.3122(0.0626) | | **3.55E-12** |
| 1319 | 0.3045(0.0757) | -0.1335(0.0212) | -0.2401(0.0191) | 3.5024(0.0849) | Excluded | |
| 1329 | 0.0479(0.0172) | 0.0015(0.0064) | 0.0093(0.0046) | 0.7665(0.0201) | Excluded | |
| 1339 | 0.0275(0.0132) | -0.0010(0.0048) | 0.0057(0.0034) | 0.5893(0.0154) | Excluded | |
| 1349 | 0.0655(0.0228) | 0.0201(0.0089) | -0.0217(0.0061) | 0.9877(0.0264) | | **8.22E-06** |
| 1359 | 0.0450(0.0175) | 0.0036(0.0065) | -0.0034(0.0046) | 0.7740(0.0203) | | 0.5708 |
| 1369 | 0.0330(0.0152) | -0.0036(0.0054) | -0.0053(0.0038) | 0.6699(0.0176) | Excluded | |
| 1379 | 0.0239(0.0106) | 0.0031(0.0040) | -0.0049(0.0028) | 0.4737(0.0124) | | 0.1112 |
| 1408 | 0.0589(0.0261) | 0.0135(0.0098) | -0.0192(0.0068) | 1.1164(0.0303) | | 0.0021 |
| 1458 | 0.5162(0.1692) | -0.0161(0.0610) | 0.0156(0.0441) | 7.4429(0.1962) | Excluded | |

| | | | | | | |
|---|---|---|---|---|---|---|
| 1478 | 0.0639(0.0164) | 0.0147(0.0062) | -0.0104(0.0043) | 0.6699(0.0187) | | 0.0016 |
| 1518 | 0.0302(0.0073) | -0.0008(0.0027) | -0.0007(0.0019) | 0.3048(0.0084) | Excluded | |
| 1528 | 0.3490(0.1250) | 0.0833(0.0472) | 0.0414(0.0326) | 5.2760(0.1440) | | 0.1311 |
| 1110 | 0.1214(0.0381) | 0.0006(0.0136) | 0.0055(0.0097) | 1.6160(0.0439) | | 0.8467 |
| 2237 | 0.0178(0.0056) | 0.0011(0.0020) | 0.0017(0.0014) | 0.2393(0.0065) | | 0.4698 |
| 2139 | 2.1346(0.3641) | 0.1096(0.1260) | -0.1346(0.0874) | 12.2547(0.3983) | | 0.1593 |
| 1160 | 0.0873(0.0248) | -0.0057(0.0087) | 0.0128(0.0064) | 1.0656(0.0286) | Excluded | |
| 1170 | 0.0702(0.0134) | -0.0016(0.0046) | -0.0068(0.0033) | 0.5262(0.0150) | Excluded | |
| 1180 | 0.1505(0.0234) | -0.0005(0.0076) | -0.0079(0.0055) | 0.7795(0.0254) | Excluded | |
| 1190 | 0.0317(0.0076) | -0.0017(0.0026) | -0.0001(0.0019) | 0.3104(0.0086) | Excluded | |
| 1200 | 0.0503(0.0112) | -0.0053(0.0038) | 0.0002(0.0028) | 0.4649(0.0127) | Excluded | |
| 1210 | 0.0166(0.0052) | -0.0004(0.0018) | 0.0015(0.0013) | 0.2021(0.0059) | Excluded | |
| 1239 | 0.0103(0.0038) | 0.0016(0.0014) | -0.0041(0.0010) | 0.1644(0.0044) | Excluded | |
| 1249 | 0.1658(0.0433) | -0.0182(0.0140) | 0.0322(0.0105) | 1.6177(0.0487) | Excluded | |
| 20116 | 0.0565(0.0109) | 0.0042(0.0039) | 0.0006(0.0028) | 0.4246(0.0122) | | 0.5405 |
| 20160 | 0.0214(0.0053) | -0.0009(0.0018) | 0.0021(0.0013) | 0.2107(0.0060) | Excluded | |
| 1717 | 0.0608(0.0078) | 0.0027(0.0026) | 0.0055(0.0020) | 0.2714(0.0084) | | 0.0181 |
| 1687 | 0.0148(0.0026) | -0.0005(0.0009) | 0.0000(0.0006) | 0.0993(0.0029) | Excluded | |
| 1697 | 0.0319(0.0027) | 0.0012(0.0009) | -0.0001(0.0007) | 0.0809(0.0028) | | 0.3524 |
| 20022 | 0.0494(0.0160) | 0.0051(0.0045) | 0.0035(0.0033) | 0.3692(0.0172) | | 0.3576 |
| 2714 | 0.4658(0.0983) | -0.0174(0.0268) | 0.0073(0.0194) | 2.0974(0.1042) | Excluded | |
| 2734 | 0.1358(0.0520) | -0.0088(0.0143) | 0.0016(0.0102) | 1.2515(0.0562) | Excluded | |
| 2375 | 0.0427(0.0107) | 0.0033(0.0026) | -0.0027(0.0019) | 0.1566(0.0111) | | 0.0858 |
| 2395 | 0.5182(0.0654) | -0.0055(0.0146) | 0.0025(0.0109) | 0.6791(0.0649) | Excluded | |
| 1920 | 0.0278(0.0054) | 0.0013(0.0019) | 0.0013(0.0014) | 0.2082(0.0060) | | 0.5842 |
| 1930 | 0.0141(0.0051) | 0.0012(0.0019) | 0.0009(0.0013) | 0.2199(0.0059) | | 0.7024 |
| 1940 | 0.0198(0.0046) | 0.0023(0.0017) | -0.0001(0.0012) | 0.1794(0.0052) | | 0.3347 |
| 1950 | 0.0129(0.0052) | 0.0008(0.0019) | -0.0012(0.0013) | 0.2260(0.0060) | | 0.5622 |
| 1960 | 0.0229(0.0051) | 0.0008(0.0018) | 0.0010(0.0013) | 0.2072(0.0058) | | 0.6997 |
| 1970 | 0.0097(0.0037) | 0.0005(0.0014) | -0.0008(0.0010) | 0.1623(0.0044) | | 0.5919 |
| 1980 | 0.0226(0.0053) | 0.0000(0.0019) | -0.0004(0.0014) | 0.2168(0.0061) | Excluded | |
| 1990 | 0.0167(0.0033) | 0.0014(0.0012) | -0.0027(0.0009) | 0.1294(0.0037) | | 0.0011 |
| 2000 | 0.0186(0.0055) | 0.0007(0.0020) | -0.0003(0.0014) | 0.2242(0.0063) | | 0.9084 |
| 20127 | 1.1075(0.2742) | 0.0482(0.0926) | 0.0165(0.0652) | 9.0996(0.3043) | | 0.8651 |

| | | | | | | |
|---|---|---|---|---|---|---|
| 2030 | 0.0140(0.0045) | 0.0001(0.0016) | -0.0007(0.0011) | 0.1881(0.0051) | | 0.8209 |
| 2040 | 0.0144(0.0047) | 0.0032(0.0017) | -0.0019(0.0012) | 0.1881(0.0053) | | 0.0249 |
| 2080 | 0.0541(0.0156) | 0.0020(0.0053) | 0.0092(0.0040) | 0.6386(0.0178) | | 0.0659 |
| 2090 | 0.0102(0.0046) | -0.0009(0.0017) | 0.0002(0.0012) | 0.2096(0.0054) | Excluded | |
| 20023 | 772.3533(238.59) | -280.1147(75.61) | -74.0616(57.83) | 10523.65(274.88) | Excluded | |
| 2217 | 38.3644(7.0684) | 0.9720(2.3483) | -0.5732(1.6921) | 232.7557(7.7706) | | 0.8279 |
| 2443 | 0.0024(0.0008) | -0.0016(0.0002) | 0.0005(0.0002) | 0.0418(0.0009) | Excluded | |
| 2463 | 0.0045(0.0020) | -0.0020(0.0007) | -0.0002(0.0005) | 0.0890(0.0023) | Excluded | |
| 6138 | 0.1678(0.0218) | -0.0061(0.0073) | 0.0226(0.0056) | 0.8148(0.0239) | Excluded | |
| 738 | 0.0758(0.0300) | 0.0187(0.0108) | -0.0200(0.0075) | 1.1398(0.0342) | | 0.0015 |
| 135 | 0.2529(0.0698) | -0.0129(0.0244) | 0.0502(0.0178) | 2.9637(0.0800) | Excluded | |

**Table S20. Genetic variance, interaction variance and their covariance component estimates for the two phenotypes with significant interaction for the covariate PC1 across POP1+POP3 as shown in Table S18.** Here the estimates were based on phenotypes adjusted by basic plus additional confounders of fixed effects and transformed by rank-based INT. The significant level is by P < 0.05/140 = 3.57E-4 after Bonferroni correction (as shown in Table S18). SE denotes standard error. DF denotes degree of freedom.

| UKBB data-field | Phenotype | $\mathrm{var}(\mathbf{g}_0)$ (SE) | $\mathrm{var}(\mathbf{g}_1)$ (SE) | $\mathrm{cov}(\mathbf{g}_0, \mathbf{g}_1)$ (SE) | $\mathrm{var}(\mathbf{e}_0)$ (SE) | P-value by LRT (G×P RNM with PC1 vs. null model) (DF=2) for phenotypes adjusted by basic confounders | P-value by LRT (G×P RNM with PC1 vs. null model) (DF=2) for phenotypes adjusted by basic and additional confounders | P-value by LRT (G×P RNM with PC1 vs. null model) (DF=2) for phenotypes adjusted by basic and additional confounders as well as rank-based INT |
|---|---|---|---|---|---|---|---|---|
| 1309 | Fresh fruit intake | 0.0326(0.0590) | -0.0111(0.0147) | 0.0158(0.0095) | 0.9784(0.0634) | **3.31E-47** | **3.33E-10** | 0.2505 |
| 1990 | Tense / highly strung | 0.1383(0.0532) | 0.0099(0.0141) | 0.0034(0.0096) | 0.8514(0.0561) | **4.17E-05** | 0.0266 | 0.6467 |

Notes: the basic confounders are sex, year of birth, age at recruitment, genotype measurement batch, UKBB assessment centre and the first 20 ancestry PCs provided by UKBB.

For phenotypes data-field 1309, additional confounders are: “Townsend deprivation index at recruitment (data-field 189)”, “major dietary changes in the last 5 years (data-field 1538)”, “variation in diet (data-field 1548)” and “Vitamin and mineral supplements (data-field 6155)”.

For phenotypes data-field 1990, additional confounders are: “Townsend deprivation index at recruitment (data-field 189)”, “seen doctor (GP) for nerves, anxiety, tension or depression (data-field 2090)”, “seen a psychiatrist for nerves, anxiety, tension or depression (data-field 2100)”, “Illness, injury, bereavement, stress in last 2 years (data-field 6145)”, “current employment status (data-field: 6142)” and “average total household income before tax (data-field: 738)”.

**Table S21. Genetic variance, interaction variance and their covariance component estimates for the two phenotypes with significant interaction for the covariate PC2 across POP1+POP3 as shown in Table S19.** Here the estimates were based on phenotypes by basic plus additional confounders and rank-based INT. The significant level is by P < 0.05/140 = 3.57E-4 after Bonferroni correction (as shown in Table S18). SE denotes standard error. DF denotes degree of freedom.

| UKB data-field | Phenotype | $\mathrm{var}(\mathbf{g}_0)$ (SE) | $\mathrm{var}(\mathbf{g}_1)$ (SE) | $\mathrm{cov}(\mathbf{g}_0, \mathbf{g}_1)$ (SE) | $\mathrm{var}(\mathbf{e}_0)$ (SE) | P-value by LRT (G×P RNM with PC1 vs. null model) (DF=2) for phenotypes adjusted by basic confounders | P-value by LRT (G×P RNM with PC1 vs. null model) (DF=2) for phenotypes adjusted by basic and additional confounders | P-value by LRT (G×P RNM with PC1 vs. null model) (DF=2) for phenotypes adjusted by basic and additional confounders as well as rank-based INT |
|---|---|---|---|---|---|---|---|---|
| 1309 | Fresh fruit intake | 0.0239(0.0592) | 0.0251(0.0152) | -0.0118(0.0101) | 0.9507(0.0632) | **3.55E-12** | **1.86E-14** | 0.0467 |
| 1349 | Processed meat intake | 0.0459(0.0567) | 0.0209(0.0150) | -0.0233(0.0100) | 0.9330(0.0608) | **8.22E-06** | 0.0016 | 0.0065 |

Notes: the basic confounders are sex, year of birth, age at recruitment, genotype measurement batch, UKBB assessment centre and the first 20 ancestry PCs provided by UKBB.

For phenotypes data-fields 1309 and 1349, additional confounders are: “Townsend deprivation index at recruitment (data-field 189)”, “major dietary changes in the last 5 years (data-field 1538)”, “variation in diet (data-field 1548)” and “Vitamin and mineral supplements (data-field 6155)”.

**Table S22**. **Phenotypic measure of the UKBB variable qualifications by Okbay et al.'s education year mapping.** Subjects who selected several response categories were assigned the higher qualification (data fields: 6138.0.0 to 6138.0.5).

| UKBB coding | UKBB category | Education years |
|---|---|---|
| 1 | College or University degree | 20 |
| 2 | A levels/AS levels or equivalent | 13 |
| 3 | O levels/GCSEs or equivalent | 10 |
| 4 | CSEs or equivalent | 10 |
| 5 | NVQ or HND or HNC or equivalent | 19 |
| 6 | Other professional qualifications eg: nursing, teaching | 15 |
| -7 | None of the above | 7 |
| -3 | Prefer not to answer | Excluded |

**Table 23. Genetic variance, interaction variance and their covariance component estimates for qualifications (mapping by Okbay et al.'s education years) across three data designs with the covariates PC1 and PC2**. The phenotypic values were adjusted by basic plus additional confounders of fixed effects and transformed by rank-based INT. SE denotes standard error. DF denotes degree of freedom.

| Phenotype (continuous) | Data design | Covariate | $\text{var}(\mathbf{g}_0)$ (SE) | $\text{var}(\mathbf{g}_1)$ (SE) | $\text{cov}(\mathbf{g}_0, \mathbf{g}_1)$ (SE) | $\text{var}(\mathbf{e}_0)$ (SE) | *P*-value by LRT comparing with baseline model (DF = 2) |
|---|---|---|---|---|---|---|---|
| Qualifications | POP1+POP2 | PC1 | 0.1149(0.0195) | 0.0833(0.0107) | -0.0881(0.0090) | 0.8040(0.0230) | 5.19E-23 |
| | | PC2 | 0.1746(0.0201) | 0.0394(0.0062) | 0.0855(0.0083) | 0.7880(0.0224) | 1.38E-25 |
| | POP2+POP3 | PC1 | 0.0740(0.0242) | 0.0397(0.0100) | 0.0285(0.0090) | 0.8813(0.0268) | 8.76E-29 |
| | | PC2 | 0.0984(0.0249) | 0.0186(0.0062) | 0.0084(0.0084) | 0.8822(0.0271) | 0.0007 |
| | POP1+POP3 | PC1 | 0.1060(0.0229) | -0.0320(0.0073) | -0.0311(0.0059) | 0.9257(0.0257) | Excluded |
| | | PC2 | 0.0959(0.0228) | -0.0032(0.0079) | 0.0313(0.0061) | 0.9075(0.0254) | Excluded |

**Table S24**. **Genetic variance, interaction variance and their covariance component estimates for qualifications (binary mapping by Gazal et al.) across three data designs with the covariates PC1 and PC2**. The phenotypic values were adjusted by basic plus additional confounders of fixed effects and transformed by rank-based INT. SE denotes standard error. DF denotes degree of freedom.

| Phenotype ( binary) | Data | Covariate | $\mathrm{var}(\mathbf{g}_0)$ (SE) | $\mathrm{var}(\mathbf{g}_1)$ (SE) | $\mathrm{cov}(\mathbf{g}_0, \mathbf{g}_1)$ (SE) | $\mathrm{var}(\mathbf{e}_0)$ (SE) | *P*-value by LRT comparing with baseline model (DF = 2) |
|---|---|---|---|---|---|---|---|
| Qualifications “college or university degree” (1) versus other six categories (0) | POP1+POP2 | PC1 | 0.1467(0.0242) | -0.0026(0.0068) | 0.0242(0.0084) | 0.8500(0.0260) | Excluded |
| | | PC2 | 0.1462(0.0242) | 0.0068(0.0053) | -0.0276(0.0085) | 0.8411(0.0258) | 2.71E-07 |
| | POP2+POP3 | PC1 | 0.1045(0.0258) | 0.0170(0.0078) | 0.0048(0.0088) | 0.8758(0.0280) | 1.66E-05 |
| | | PC2 | 0.1184(0.0257) | 0.0108(0.0056) | -0.0041(0.0084) | 0.8696(0.0276) | 0.0043 |
| | POP1+POP3 | PC1 | 0.0928(0.0240) | 0.0207(0.0092) | 0.0272(0.0062) | 0.8855(0.0270) | 8.82E-08 |
| | | PC2 | 0.0957(0.0242) | 0.0108(0.0088) | -0.0171(0.0060) | 0.8929(0.0271) | 0.0031 |
| Qualifications “none of the above” (1) versus other six categories (0) | POP1+POP2 | PC1 | 0.2032(0.0216) | 0.0637(0.0098) | -0.0808(0.0089) | 0.7227(0.0236) | 7.77E-22 |
| | | PC2 | 0.2509(0.0221) | 0.0379(0.0063) | 0.0837(0.0086) | 0.7032(0.0232) | 1.91E-22 |
| | POP2+POP3 | PC1 | 0.1552(0.0234) | 0.0350(0.0100) | 0.0277(0.0089) | 0.7882(0.0256) | 5.67E-26 |
| | | PC2 | 0.2114(0.0235) | 0.0104(0.0053) | 0.0019(0.0078) | 0.7623(0.0246) | 0.0449 |
| | POP1+POP3 | PC1 | 0.2336(0.0233) | -0.0062(0.0078) | -0.0444(0.0063) | 0.7709(0.0245) | Excluded |
| | | PC2 | 0.2123(0.0228) | -0.0085(0.0078) | 0.0303(0.0060) | 0.7934(0.0243) | Excluded |

**Table S25. The distributions of original phenotypic values across POP1, POP2 and POP3 for the trait "age first had sexual intercourse".**

| | Min. | 1st Quartile | Median | Mean | 3rd Quartile | Max. | Variance | *N* |
|---|---|---|---|---|---|---|---|---|
| POP1 | 5.00 | 17.00 | 18.00 | 19.03 | 21.00 | 66.00 | 15.6130 | 6647 |
| POP2 | 3.00 | 17.00 | 19.00 | 19.27 | 21.00 | 62.00 | 15.5495 | 5924 |
| POP3 | 3.00 | 17.00 | 18.00 | 19.11 | 21.00 | 63.00 | 14.6723 | 7102 |

**Table S26. The distributions of original phenotypic values across POP1, POP2 and POP3 for the trait "qualifications".** Level 1: none; Level 2: O-levels or CSEs; Level 3: A-levels, NVQ, HND, HNC or other professional qualification; Level 4: college or university degree.

| | Level 1 | Level 2 | Level 3 | Level 4 | Mean | Variance | *N* |
|---|---|---|---|---|---|---|---|
| POP1 | 1303 | 2148 | 1769 | 2216 | 2.6587 | 1.1715 | 7436 |
| POP2 | 636 | 807 | 1394 | 3927 | 3.2732 | 1.0015 | 6764 |
| POP3 | 536 | 992 | 1719 | 4518 | 3.3160 | 0.8859 | 7765 |

**Table S27. The results of linear regressions including 1 (representing POP1) and 2 (representing POP2) as independent variable across POP1+POP2.** For real data of the trait qualifications, we used educational levels as dependent variable, which was adjusted by basic and additional confounders and rank-based INT (but not adjusted by PCs). For 100 replicates simulated with different selection odds ratio combinations, we used the simulated phenotypes as dependant variable, which was adjusted by rank-based INT.

| | R-squared | | Adjusted R-squared | | -log10(P-value) | |
|---|---|---|---|---|---|---|
| Real data of qualifications | 0.0045 | | 0.0044 | | 12.9914 | |
| Selection scenarios for 100 simulation replicates | Mean | SE | Mean | SE | Mean | SE |
| $OR_{\text{POP1, Y}} = 2$, $OR_{\text{POP2, Y}} = 3$ | 0.0059 | 0.0004 | 0.0058 | 0.0004 | 10.1497 | 0.6567 |
| $OR_{\text{POP1, Y}} = 2$, $OR_{\text{POP1, }Z} = 2$, $OR_{\text{POP2, Y}} = 3$, $OR_{\text{POP2, }Z} = 2$ | 0.0056 | 0.0004 | 0.0055 | 0.0004 | 9.6210 | 0.6509 |
| $OR_{\text{POP1, Y}} = 2$, $OR_{\text{POP1, }Z} = 2$, $OR_{\text{POP2, Y}} = 3$, $OR_{\text{POP2, }Z} = 3$ | 0.0035 | 0.0003 | 0.0034 | 0.0003 | 6.2513 | 0.5188 |
| $OR_{\text{POP1, Y}} = 2$, $OR_{\text{POP1, }Z} = 3$, $OR_{\text{POP2, Y}} = 3$, $OR_{\text{POP2, }Z} = 3$ | 0.0052 | 0.0004 | 0.0051 | 0.0004 | 8.9677 | 0.6478 |

**Table S28**. **SNP-based heritabilities estimated by GREML and G×P RNM methods for two traits across POP1+POP2, POP2+POP3 and POP1+POP3.** Here the phenotypes were adjusted by basic plus additional confounders of fixed effects and transformed by rank-based INT. SE denotes standard error.

| Phenotype | Estimate method | POP1+POP2 | | POP2+POP3 | | POP1+POP3 | |
|---|---|---|---|---|---|---|---|
| | | $h^2_{SNP}$ | SE | $h^2_{SNP}$ | SE | $h^2_{SNP}$ | SE |
| Qualifications | GREML | 0.0998 | 0.0262 | 0.1156 | 0.0265 | 0.1031 | 0.0247 |
| | G×P RNM with PC1 | 0.1281 | 0.0250 | 0.0971 | 0.0272 | 0.1151 | 0.0237 |
| | G×P RNM with PC2 | 0.1840 | 0.0219 | 0.1156 | 0.0269 | 0.1090 | 0.0241 |
| Age first had sexual intercourse | GREML | 0.1000 | 0.0266 | 0.0933 | 0.0258 | 0.1582 | 0.0250 |
| | G×P RNM with PC1 | 0.1015 | 0.0267 | 0.0962 | 0.0264 | 0.1574 | 0.0249 |
| | G×P RNM with PC2 | 0.1023 | 0.0268 | 0.0947 | 0.0261 | 0.1591 | 0.0252 |

**Table S29**. **Genetic correlation estimates between population groups (POP1, POP2 and POP3) by bivariate GREML for qualifications.** Here the qualifications were reclassified into binary measures, adjusted by basic plus additional confounders of fixed effects and transformed by rank-based INT. SE denotes standard error. P-value was obtained through a Wald test under a null hypothesis that genetic correlation equals to 1.

| Binary phenotype | Genetic correlation between POP1 and POP2 | | | Genetic correlation between POP2 and POP3 | | | Genetic correlation between POP1 and POP3 | | |
|---|---|---|---|---|---|---|---|---|---|
| | Estimate | SE | P value | Estimate | SE | P value | Estimate | SE | P value |
| “college or university degree” versus other six educational categories | 0.2940 | 0.3353 | 0.0352 | 0.7111 | 0.4226 | 0.4942 | 0.9551 | 0.6020 | 0.9405 |
| “none of the above” versus other six educational categories | -0.0232 | 0.2539 | 5.58E-05 | 0.3134 | 0.1735 | 7.59E-05 | 0.4315 | 0.3045 | 0.0619 |
